# Autism genes converge on three functional programs organized by neuronal subclass, developmental timing, and cortical patterning

**DOI:** 10.64898/2026.09.22.753508

**Authors:** Rachel L. Smith, Yundan Liao, Lambertus Klei, Lujing Zhang, Yunlong Ma, Miao Tang, F. Kyle Satterstrom, Chiara Auwerx, Jack M. Fu, Joseph D. Buxbaum, Mark J. Daly, Michael E. Talkowski, Luis de la Torre-Ubieta, Yevgenia Kozorovitskiy, Matthew L. MacDonald, Kathryn Roeder, Bernie Devlin, Michael J. Gandal

**Affiliations:** Departments of Psychiatry, Genetics, and Pediatrics, Perelman School of Medicine, University of Pennsylvania, Philadelphia, PA, USA; The Lurie Autism Institute at Penn Medicine and the Children’s Hospital of Philadelphia, University of Pennsylvania, Philadelphia, PA, USA; The Lifespan Brain Institute at Penn Medicine and the Children’s Hospital of Philadelphia, University of Pennsylvania, Philadelphia, PA, USA; Department of Psychiatry, University of Pittsburgh School of Medicine, Pittsburgh, PA, USA; Department of Statistics and Data Science, Carnegie Mellon University, Pittsburgh, PA, USA; Stanley Center for Psychiatric Research, Broad Institute of MIT and Harvard, Cambridge, MA, USA; Analytic and Translational Genetics Unit, Department of Medicine, Massachusetts General Hospital, Boston, MA, USA; Program in Medical and Population Genetics, Broad Institute of MIT and Harvard, Cambridge, MA, USA; Center for Genomic Medicine, Massachusetts General Hospital, Boston, MA, USA; Department of Neurology, Massachusetts General Hospital and Harvard Medical School, Boston, MA, USA; Seaver Autism Center for Research and Treatment, Icahn School of Medicine at Mount Sinai, New York, NY, USA; Department of Psychiatry, Icahn School of Medicine at Mount Sinai, New York, NY, USA; Department of Genetics and Genomic Sciences, Icahn School of Medicine at Mount Sinai, New York, NY, USA; Friedman Brain Institute, Icahn School of Medicine at Mount Sinai, New York, NY, USA; The Mindich Child Health and Development Institute, Icahn School of Medicine at Mount Sinai, New York, NY, USA; Department of Neuroscience, Icahn School of Medicine at Mount Sinai, New York, NY, USA; Institute for Molecular Medicine Finland, University of Helsinki, Helsinki, Finland; Department of Medicine, Harvard Medical School, Boston, MA, USA; Program in Bioinformatics and Integrative Genomics, Harvard Medical School, Boston, MA, USA; Department of Psychiatry and Biobehavioral Sciences, David Geffen School of Medicine, University of California Los Angeles, Los Angeles, CA, USA; Intellectual and Developmental Disabilities Research Center, Semel Institute for Neuroscience and Human Behavior, University of California Los Angeles, Los Angeles, CA, USA; Department of Neurobiology, Northwestern University, Evanston, IL, USA; Computational Biology Department, Carnegie Mellon University, Pittsburgh, PA, USA

**Keywords:** Autism, single-cell genomics, gene regulatory networks, cortical development

## Abstract

Our companion sequencing study uncovered 253 genes robustly associated with autism spectrum disorder (ASD), yet the biological programs they impact, and the cellular, developmental, and spatial contexts in which they converge, remain unresolved. Here we systematically contextualize genes associated with ASD across neurodevelopment and cortical areas, integrating developmental single-cell and spatial atlases with gene- and isoform co-expression, regulatory, proteomic, and synaptic networks. Genetic burden concentrates within temporally-resolved neuronal subclasses: newborn excitatory neurons, immature interneurons, and maturing intratelencephalic lineages. Genes associated with ASD converge on three functional programs—gene regulation, neuronal morphogenesis, and synaptic transmembrane signaling machinery—resolved from 28 ASD-associated networks, several of which are directly regulated by ASD genes including ***MEF2C*, *SOX11***, and ***FOXP2***. These programs are spatially patterned, with risk genes exhibiting an increasing anterior-to-posterior cortical expression gradient, anchored in the primary visual cortex and driven by excitatory neuron gene-regulatory programs. Finally, ASD risk genes associated with more severe developmental phenotypes show broader excitatory neuron enrichment and less interneuron involvement. Together, these findings anchor ASD genetic vulnerability to specific neurodevelopmental lineages, epochs, and molecular substrates, and delineate the features that distinguish ASD from comorbidities with broader developmental impacts.

## Introduction

Autism spectrum disorder (ASD) is a complex neurodevelopmental condition associated with broad phenotypic heterogeneity and clinical comorbidity. Genetic factors contribute substantially to the population-level liability for ASD, with hundreds of genetic loci identified to date ^1–4^. Most ASD-associated genes have been identified by whole exome sequencing and harbor an excess of rare, damaging ***de novo*** mutations ^2,5–7^. The Autism Sequencing Consortium (ASC) has now identified 253 rare variant-associated genes achieving false discovery rate (FDR) < 0.001 ^8^. Despite this progress, our understanding of the specific underlying developmental, cellular, and molecular programs through which genetic effects converge remains limited. Consequently, a major current focus for ASD genomics is to move beyond a list of individual genes to a concrete set of convergent mechanistic insights ^4,9^.

As gene function is highly context-dependent, a central challenge is to determine the cellular, developmental, and spatial contexts in which ASD-associated genes act. Early integrative analyses of genetic risk and developmental brain transcriptomics implicated the mid-fetal cerebral cortex and cortical projection neurons as major points of convergence ^10,11^. Case-control postmortem studies have provided a complementary view, revealing widespread cortical transcriptomic dysregulation in ASD with prominent effects in excitatory neurons ^12–14^. These changes are spatially patterned: molecular differences among cortical areas established through developmental arealization ^15,16^ are attenuated in ASD, with dysregulation increasing along an anterior-to-posterior (A-P) gradient and greatest in primary visual cortex ^12^. Yet these studies provide only limited resolution of the developmental and cellular contexts of genetic risk. Single-cell, multiomic, and spatially resolved atlases of human brain development ^17–20^ now allow ASD genetic association to be localized across neuronal lineages, developmental trajectories, and cortical regions at substantially finer resolution.

A second challenge is to resolve the specific biological programs in which genes associated with ASD converge. Rare variant studies have consistently implicated broad functional categories including gene and chromatin regulation and synaptic function ^5,7,21^, but these categories encompass diverse molecular mechanisms. Data-driven transcriptomic studies have identified reproducible ASD-associated co-expression and regulatory networks ^12–14^, while large developmental datasets have defined gene- and isoform-level co-expression modules and transcriptional regulons across human neurodevelopment ^20,22–24^. Together, these resources allow rare-variant association to be mapped systematically onto data-driven programs and molecular networks, while expert-curated ontologies such as SynGO can further resolve broad synaptic convergence into specific functions and subcellular compartments ^25^.

Finally, it remains unknown whether these cellular contexts and biological programs are shared across the heterogeneous phenotypic spectrum of ASD, such as in the context of comorbid developmental delay and intellectual disability (DD/ID). Probands with DD/ID carry a higher burden of damaging *de novo* mutations (DNMs) than those without ^2,26^. Stratifying genetic association by DD/ID comorbidity may therefore distinguish cellular and developmental contexts associated with broader neurodevelopmental impacts from those associated with ASD in the absence of DD/ID.

Here, we integrate the most recent ASD rare-variant association (RVAS) results from the ASC ^8^, including 253 high-confidence ASD risk genes, with developmental single-cell and spatial atlases of the human brain ^17–20^ and a curated set of 1,151 gene co-expression modules and transcriptional regulons ^12,14,20,24^. We address two complementary questions: in which cell types, developmental windows, and cortical regions does ASD rare-variant association concentrate, and which convergent biological programs do these ASD genes collectively disrupt? We further examine the degree to which these patterns differ by comorbid DD/ID. Together, our findings reveal a coherent biological architecture underlying ASD, and provide a framework for prioritizing genes, cell types, functional programs, and developmental windows for future investigation (**Fig. 1A**).

**Figure 1.**
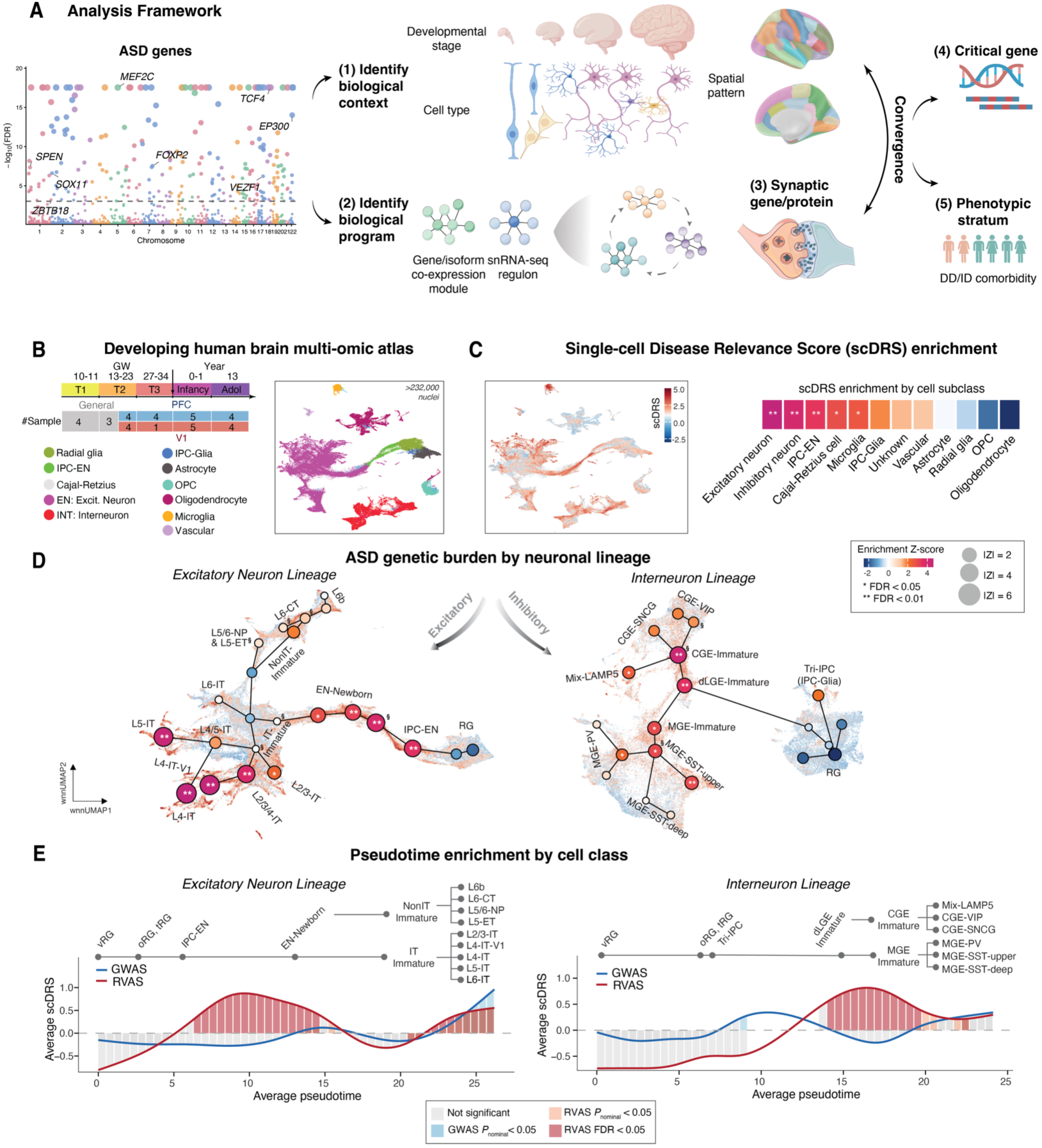
ASD genes localize within specific developmental windows and neuronal sub-classes. **(A)** Analysis framework to contextualize ASD rare-variant genes by (1) identifying their biological context (developmental stage, cell type, and spatial pattern) using single-cell and spatial atlases of the human brain ^18–20,30^, and (2) identifying the biological programs they collectively disrupt using 1,151 gene/isoform co-expression modules and transcriptional regulons ^12,14,20,24^, and (3) resolving the broad synaptic signal through SynGO ontology to pinpoint the most enriched synaptic functions and compartments. Convergence across these analyses prioritized (4) critical genes and enabled (5) stratification of these patterns by DD/ID comorbidity. **(B)** Single-cell multi-omic atlas of 232,328 nuclei from the developing human neocortex, spanning first-trimester to early adolescence from ^20^; GW = gestational week. UMAP shows major progenitor, neuron, and glial cell-types. Significance: ** FDR < 0.01; * FDR < 0.05. **(C)** ASD genetic enrichments within cell types calculated with scDRS. Left: UMAP colored by normalized scDRS scores. Right: scDRS group-level enrichment *Z*-scores for each subclass. Significance: ** FDR < 0.01; * FDR < 0.05. **(D)** scDRS enrichment along excitatory (EN; left) and inhibitory (IN; right) lineage trajectories. Node size reflects enrichment magnitude (|*Z*-score|) and color reflects group-level enrichment *Z*-score. Edges denote inferred developmental relationships between lineage clusters. § indicates a lineage cluster which reflects a mixture of constituent cell types (reference annotations as in **B**); the group-level enrichment *Z*-score is computed on the pooled population (see **Fig. S1** for per-cluster composition). **(E)** Temporal characterization of ASD genetic enrichment along excitatory and inhibitory neuron lineage pseudotime trajectories. ASD rare variant association is observed for IPC, newborn, and IT lineage excitatory neurons (left), and immature and specific mature interneurons (right). Common variant association from GWAS is shown in blue for comparison ^31^. Average scDRS per pseudotime bin for RVAS and GWAS, curve color indicates bin-level significance.

## Results

### ASD rare genetic burden concentrates in immature interneurons, newborn excitatory, and intratelencephalic lineages

To determine the specific cellular and developmental contexts in which 253 high-confidence ASD genes converge ^8^, we mapped RVAS statistics to single cells within a comprehensive multi-omic atlas of human neurodevelopment spanning prenatal first-trimester to postnatal adolescent stages ^20^. We adopted the single-cell disease relevance score (scDRS) framework to systematically characterize genetic enrichments across multiple neurodevelopmental hierarchies—from individual cells to major cell subclasses, lineages, subtypes, laminae, and regional axes. We observed pronounced ASD genetic enrichments within neuronal (versus non-neuronal) classes, with excitatory neurons (ENs) exhibiting the strongest signals followed by interneurons (IN; **Fig. 1B-C**; **Table S1A**). Our primary analyses used the exome-wide significant genes (Transmission and *De Novo* Association [TADA] FDR < 0.001), although the observed cell-type-specific vulnerability profile remained robust across expanded gene sets at relaxed FDR thresholds (< 0.01 [*n* = 416 genes], < 0.05 [*n* = 696], < 0.1 [*n* = 951]; **Methods**; **Fig. S2A-D**).

Driven by these prominent neuronal signals, we next tracked genetic enrichments along the EN lineage developmental trajectory, which revealed a bimodal enrichment pattern (**Fig. 1D**; **Fig. S1**; **Table S1B**). The first peak coincided with transition from intermediate progenitor cells (IPC) to newborn ENs, whereas the second peak captured ENs within the intratelencephalic (IT) lineage. Within the IT branch, enrichment spanned cortical layers 2–5 and was strongest in L4-IT neurons. This signal was notably lineage-specific, with preferential enrichment of the IT lineage over extratelencephalic (ET) projecting, corticothalamic (CT), and near-projecting (NP) lineages. IT neurons receive thalamocortical input, engage in broad cortico-cortical connectivity, and represent the overwhelming majority of cortico-striatal projections, whereas ET and CT populations primarily target subcortical structures ^27–29^. Thus, ASD genetic signals are concentrated in circuits mediating intracortical processing and downstream cortico-striatal communication.

The inhibitory neuron (IN) lineage also exhibited significant associations (**Fig. 1D**; **Fig. S1**; **Table S1B**), particularly within immature subtypes. INs originate from the ganglionic eminence, an ephemeral brain structure with three subdivisions: the caudal ganglionic eminence (CGE), dorsolateral ganglionic eminence (dLGE), and medial ganglionic eminence (MGE). The strongest enrichment was observed in CGE-derived immature INs, followed by significant associations in dLGE and MGE immature INs. Specific mature IN subtypes were also enriched, including CGE-derived LAMP5-expressing and MGE-derived Somatostatin-expressing (SST) clusters. MGE-SST neurons are of special interest because somatostatin produced by these neurons has been implicated in EN maturation ^20^. MGE-SST neurons have also been shown to signal to multiple mature EN types, and interact extensively with mature IT neurons ^20^, motivating a closer examination of the internal heterogeneity of the MGE-SST lineage.

Within the MGE-SST lineage, scDRS scores were non-uniformly distributed suggesting additional subtype specificity (**Fig. 1D, right**). The uniform manifold approximation and projection (UMAP) of the SST neuron subset revealed a segregation into two subtypes with differential ASD association (**Fig. S3A**). The enriched subtype was characterized by higher expression of upper-layer SST marker genes (e.g., *SYTL5*) based on the Allen Brain reference atlases ^32,33^. In contrast, the depleted subtype highly expressed the deep-layer markers (e.g., *B3GAT2*). These annotations were further supported by characterization by MapMyCell ^34^ and differential expression analyses between subtypes (**Supplement**; **Figs. S3B, S4**; **Tables S2, S3**). We hereafter refer to these as the upper-layer and deep-layer MGE-SST subtypes, respectively. MGE-derived SST interneurons exert layer- and subtype-specific control over pyramidal-cell dendrites: SST inhibitory neurons gate the integration of intracortical and higher-order thalamic inputs by superficial L2/3 IT neurons, while deep-layer SST subtypes are more likely to regulate dendritic compartments of L5 IT and ET/pyramidal tract (PT) pyramidal neurons ^35^.

### Genes associated with ASD show distinct trajectories across excitatory neuron and interneuron lineages

To further dissect the temporal dynamics underlying observed cell-type-specific enrichments, we integrated RVAS results with single-cell pseudotime trajectories across EN and IN lineages (**Methods**; **Fig. 1E**), which corresponded closely with underlying developmental stages (**Fig. S5A**). We additionally integrated common variant association results from ASD GWAS using scDRS to examine how rare and common variant signals were distributed across developmental stages (**Methods**) ^31^. Across developmental stages, enrichment was greater for rare variant associations (RVAS; **Table S1C**) than for common variant associations (GWAS; **Table S4A**) in both EN and IN lineages (**Fig. S5A**). Reflecting the underpowered nature of current ASD GWAS, no significant common variant enrichment was observed in either lineage, although a marginal signal was detected within second trimester ENs (*P* = 0.052).

In the EN lineage, RVAS enrichment was significant in the first and second trimesters, infancy, and adolescence (**Fig. S5A**). At finer pseudotime resolution, these broad-stage enrichments elicited a bimodal pattern, with the stage-level enrichments corresponding to two pseudotime peaks along the continuous trajectory: one at the newborn EN stage and the other at the terminal pseudotime stage of excitatory IT neuron development (**Fig. 1E**; **Tables S1D-E, S4B-C**). Notably, this terminal window represented a point of convergence between rare- and common-variant signals. GWAS enrichment rose to nominal significance at this stage, aligning with the secondary RVAS peak. In the IN lineage, RVAS enrichment spanned the second trimester through adolescence, peaking in the third trimester (**Fig. S5A**). At finer pseudotime resolution, this enrichment resolved into a single pseudotime window concentrated at the immature IN stage, whereas GWAS showed no significant enrichment along the trajectory. Sensitivity analyses yielded highly consistent developmental enrichment patterns for ASD genes defined across TADA FDR thresholds (**Fig. S5B-C**).

### Posterior-predominant regional patterning of ASD genetic signal

Human brain development and maturation proceed along regionally distinct trajectories, producing a spatially patterned neocortex with specialized cytoarchitectural and connectivity profiles ^16,17,20,36^. Transcriptomic studies have similarly revealed regional variation in ASD-associated molecular pathology, including attenuated cortical patterning and an anterior-to-posterior gradient in differential expression effect size ^12^. To test whether rare variants that contribute risk to ASD exhibit region-specific association, we computed group-level scDRS *Z*-scores separately for single cells across anterior (prefrontal cortex; PFC) and posterior (primary visual cortex; V1) brain regions for 11 donors with paired samples. scDRS enrichments were highly divergent across regions for EN—but not IN—lineages (**Fig. 2A**; **Table S1F**). Pooled across all developmental stages, EN enrichments were significantly greater in V1 than PFC (paired t-test, *P* = 0.026), whereas INs exhibited no regional differences (*P* = 0.535). To further resolve this pattern, we repeated scDRS group-level analyses with cluster-level resolution, using both the full dataset and the paired-donor subset. Here again, we observed greater V1 association for EN lineage clusters across developmental stages, particularly for IT neurons (**Fig. S6A-B**). In contrast, IN subtypes did not exhibit consistent regional associations across stages (**Fig. S6A-B**). Together, these results suggest that, within the excitatory neuron lineage, ASD genes may have a greater developmental impact in the posterior brain regions—closely mirroring post-mortem transcriptomic findings ^12^.

**Figure 2.**
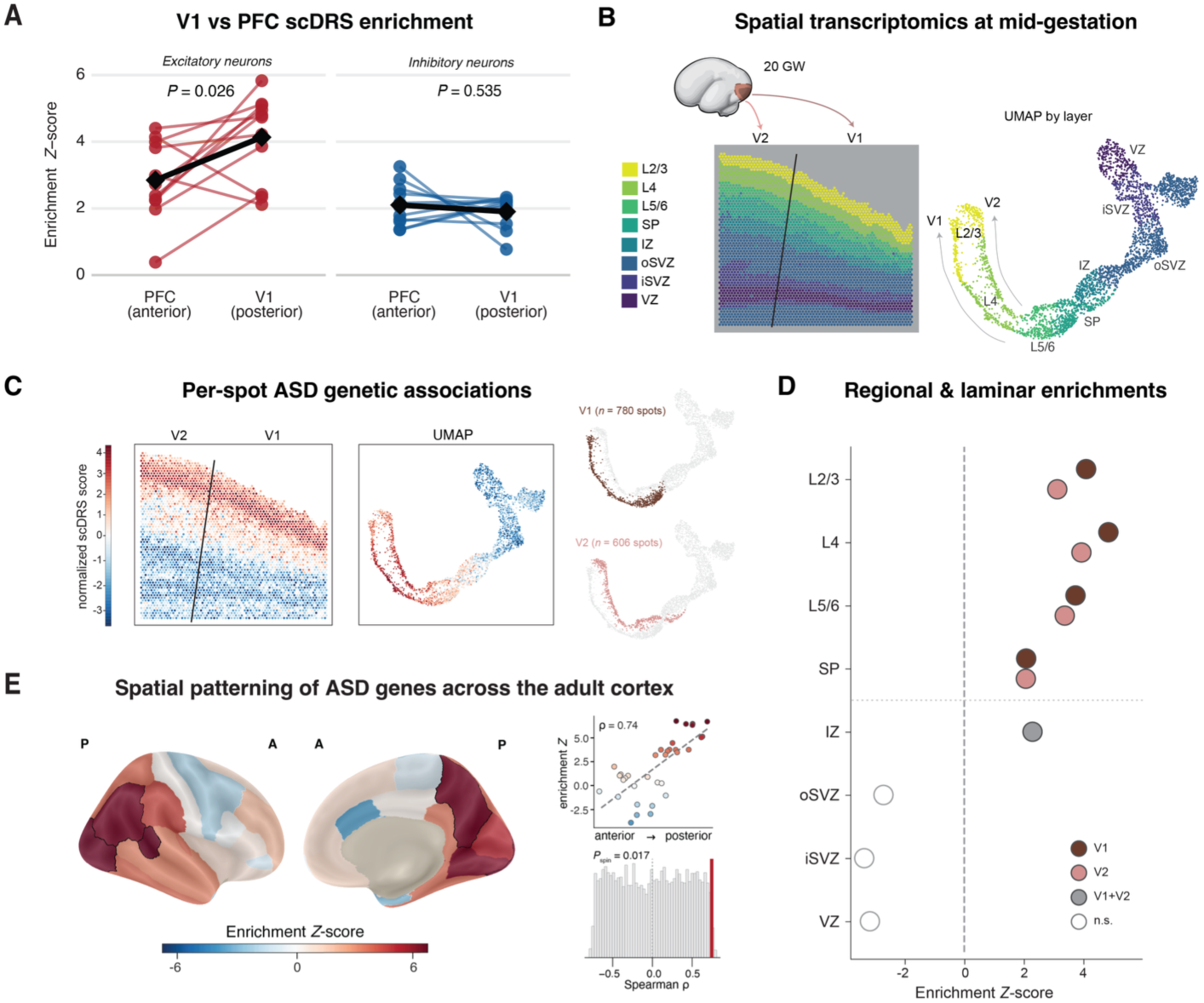
Genes associated with ASD show posterior cortical expression patterns, anchored in V1. **(A)** Group-level scDRS enrichment *Z*-scores computed for 11 donors with paired PFC and V1 samples, pooling excitatory neurons and interneuron classes across developmental stages. Colored lines, individual donors; black line and diamond, cohort mean. *P* values, two-sided paired t-test; *n* = 11 donors. **(B)** Left: laminar annotation of 10X Visium spatial transcriptomics dataset spanning V1 and V2 at 20 weeks gestation ^17^. The black line indicates the V1/V2 boundary. Right: UMAP of spots colored by tissue compartment show clear transcriptomic separation of V1 and V2 regions of the cortical plate. *N* = 1 section from 1 donor; 3,591 spots. **(C)** ASD scDRS genetic association normalized scores are shown for each individual spatial transcriptomic spot, as well as for the joint V1 and V2 UMAP. **(D)** ASD scDRS enrichments are shown for cortical layers and germinal zones across V1 and V2, quantified as group-level enrichment *Z*-score. Below the cortical plate (dotted line), V1 and V2 spots were pooled as the areal boundary cannot be reliably assigned from anatomical position in these compartments. Point color indicates region; filled circles indicate FDR < 0.05. **(E)** Spatial enrichment patterning of ASD genes across the adult cortex. Enrichment significantly increases along an anterior-posterior axis (*ρ* = 0.74; *P*_spin_ = 0.017), with posterior regions exhibiting significant enrichment compared to spatial autocorrelation-corrected background (FDR < 0.05). Individual regions whose enrichment is FDR significant above the spin null are outlined in black. The histogram shows the spin null distribution along with the observed correlation.

V1, the primary recipient of thalamic visual inputs into the cortex, is a cytoarchitecturally and transcriptomically distinct region with an expanded population of upper-layer neurons ^17,30^. As V1 is also among the most posterior regions of the human cortex, we could not readily distinguish whether observed ASD genetic enrichments in V1 reflect its distinct cytoarchitecture or its posterior positioning. We therefore next leveraged a spatial transcriptomics atlas of the developing visual cortex at gestational week 20, at which point the unique V1 laminar structure has begun to emerge ^17^ (**Fig. 2B**). We quantified the spatial distribution of ASD genetic enrichments across the cortical wall for V1 and the adjacent secondary visual cortex (V2), applying the scDRS framework for spot-level spatial transcriptomic data (**Methods**). ASD genetic enrichment was significant (FDR < 0.05) across both V1 and V2, particularly within the post-mitotic neuronally enriched layers of the cortical plate and subplate—but not in the germinal zones (**Fig. 2C,D**). Within these compartments, V1-L4 showed the strongest laminar association with ASD. V1 enrichments were consistently higher than V2, although individual laminar associations were highly significant within V2 itself (**Fig. 2C,D**). Like the within-donor PFC-V1 paired single-cell analyses above, the differences here are unlikely to be explained by technical factors or subject demographics. Rather, results suggest that both posterior cortical patterning and inferred thalamocortical primary sensory inputs contribute to the observed regionality of ASD genetic associations. Further, these processes are likely contributing as early as 20 gestational weeks.

Motivated by this heightened V1 posterior association, we next asked whether genes associated with ASD continue to show spatially patterned enrichment in the adult cortex, using the adult multi-region Allen Human Brain Atlas (AHBA), with samples mapped to the Desikan-Killiany (DK) cortical parcellation ^30,37^. ASD genes continued to demonstrate a non-uniform spatial expression pattern, with an increasing expression gradient along the anterior-posterior (A-P) axis (*ρ* = 0.74; *P*_spin_ = 0.017; **Fig. 2E**; **Fig. S7A**). This gradient was most pronounced across 5 posterior regions—the lateral occipital, banks of the superior temporal sulcus, inferior parietal, precuneus, and lingual regions—each individually enriched above both the gene-set resampling null (*Z* = 6.3-6.8, FDR < 0.05) and the spatial autocorrelation (spin) null (all *P*_spin_ < 0.05; **Methods**; **Table S5**). As the DK parcellation does not delineate V1 as a distinct parcel, we additionally tested a focal occipital cluster (pericalcarine, lateral occipital, lingual, and cuneus); this cluster was likewise significant (*Z* = 5.8; *P*_spin_ = 0.037; **Table S5**), consistent with the occipital enrichments observed in the single-cell and spatial transcriptomic data (**Fig. 2A-D**). These findings were robust across sensitivity analyses: occipital enrichment remained significant against a null matched on gene expression level, constraint, and coding sequence length, and was not driven by a small subset of genes with high occipital expression (**Fig. S7B-C**). Furthermore, the A-P gradient association persisted even after excluding the four occipital regions defined above, indicating a broad cortical pattern (**Fig. S7D**). These findings demonstrate that the posterior- and V1-predominant ASD enrichments observed within developmental single-cell and spatial datasets persist well into adulthood and are reproduced by bulk regional cortical transcriptomics. Additionally, results suggest that ASD gene patterning encompasses both a continuous A-P gradient and a discrete V1/V2 difference, mirroring the two modes of cortical arealization ^17^.

### ASD rare variant burden converges on 28 distinct regulatory networks

We next asked which molecular programs underlie the cellular, developmental, and spatial patterns of ASD associations described above. As established gene-set annotations incompletely capture the molecular programs governing human brain development and function, we adopted an unsupervised data-driven approach, assembling a broad compendium of 1,151 networks from large-scale genomic studies profiling neurodevelopment and ASD. These included co-expression defined network modules, as well as single-cell gene regulatory networks, with individual transcriptional ‘regulons’ defined based on transcription factor (TF) co-expression with motif-inferred target genes (collectively, “modules”). These modules were drawn from four independent sources: (1) developmental gene- and isoform-level co-expression networks ^24^; (2) ASD-associated pancortical gene- and isoform-level co-expression networks ^12^; (3) developmental transcriptional regulons ^20^; and (4) ASD-associated regulons ^14^ (**Fig. 3A**).

**Figure 3.**
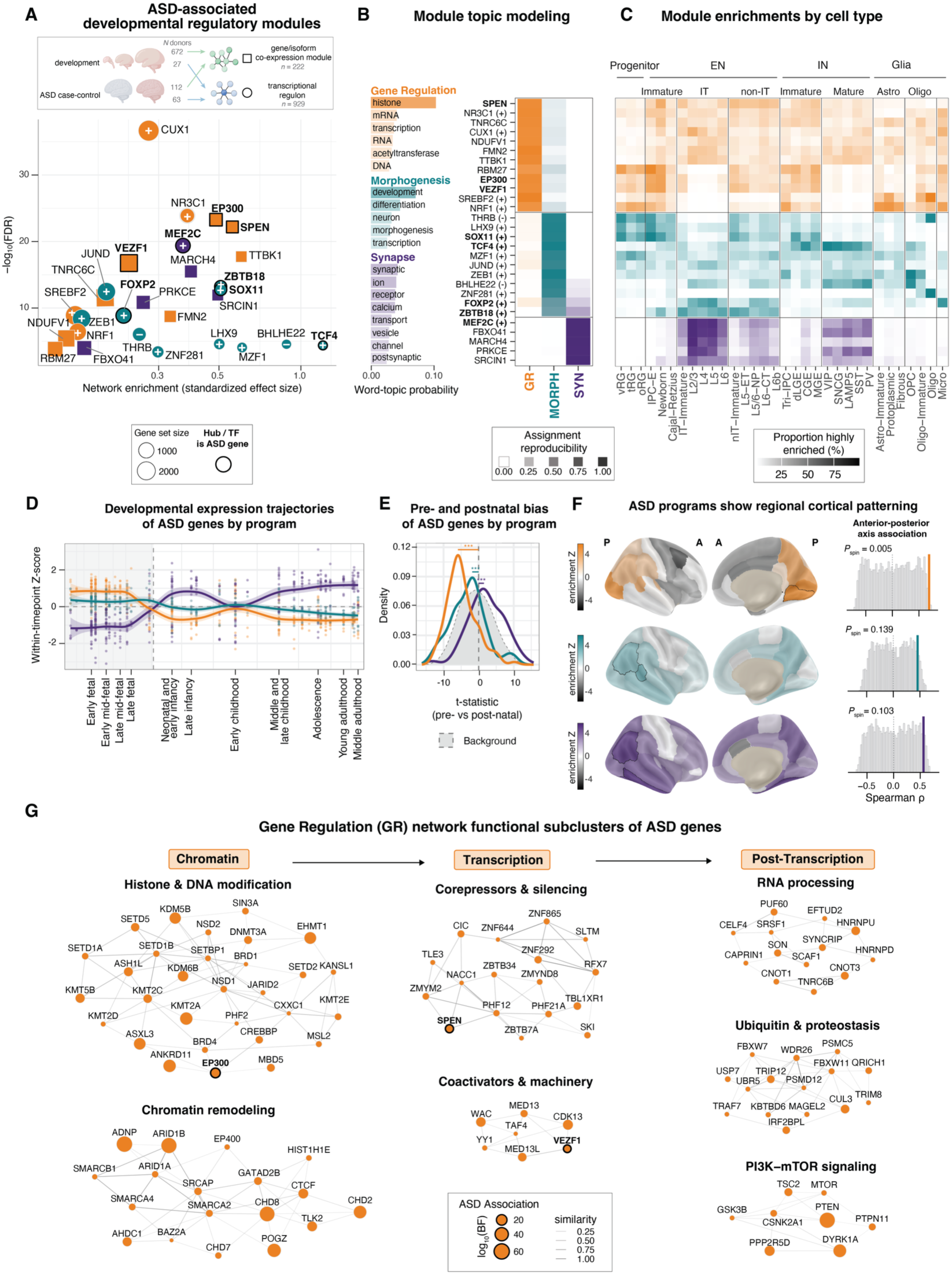
ASD-associated modules converge on gene-regulation, morphogenesis, and synaptic pathways. **(A)** Top: Modules included in this analysis were collated across four studies: 1. Developmental gene and isoform co-expression modules (*n* = 124) from *N* = 672 donors ^24^; 2. Developmental transcriptional regulons (*n* = 583) from *N* = 27 donors ^20^; 3. ASD case-control gene and isoform co-expression modules (*n* = 98) from *N* = 112 donors ^12^; 4. ASD case-control transcriptional regulons (*n* = 346) from *N* = 63 donors ^14^. Bottom: Twenty-eight gene co-expression modules and transcriptional regulatory networks are associated with ASD rare variant burden. Each point is a module; x-axis, standardized effect size (*β*); y-axis, −log_10_(FDR). Color indicates biological program; size indicates number of genes or isoforms. Bold indicates modules whose representative gene (the co-expression hub or regulon driver TF) is itself a high-confidence ASD gene (TADA FDR < 0.001). **(B)** Left, topic modeling identifies three distinct biological programs underlying ASD regulatory network associations, characterized by gene regulation (GR), neuronal morphogenesis (MORPH), and synapse-related (SYN) pathways. Barplot shows the probability of each term within the program, all significantly enriched relative to cross-program mean (FDR < 0.05). Right, ASD-associated regulatory modules show program-level clustering. Opacity indicates the number of times the module was assigned to the program across 1000 topic model solutions. As in panel A, modules in which the representative gene is itself an ASD gene are in bold. **(C)** Cell-type associations for ASD regulatory networks, using AUCell ^40^ and data from Fig 1. SYN modules show strongest enrichments for IT neurons and interneurons, whereas GR and MORPH networks exhibit broader, less specific patterns. **(D, E)** Developmental expression trajectories are shown for program-assigned ASD genes. GR and MORPH genes exhibit prenatally-biased expression patterns (FDR = 2.42×10⁻¹⁹ and 3.81×10⁻⁴, respectively), whereas SYN network expression is postnatally predominant (FDR = 3.88×10⁻⁵). Plots show mean within-timepoint expression *Z*-scores across 11 developmental stages from BrainSpan ^23^. **(F)** Left: Spatial enrichment gradients of program-assigned ASD genes across the adult cortex. Regions outlined in black are significantly enriched after accounting for spatial autocorrelation (FDR < 0.05). Right: GR exhibits a significant A-P gradient (*ρ* = 0.69, *P*_spin_ = 0.005); SYN and MORPH do not reach significance after accounting for spatial autocorrelation (*ρ* = 0.56, *P*_spin_ = 0.10 and *ρ* = 0.46, *P*_spin_ = 0.14, respectively). **(G)** The GR-assigned ASD genes (*n* = 128) cluster into functional subgroups of gene regulation from chromatin remodeling to transcription to post-transcriptional processing, assessed by GO term membership cosine similarity. Gene size indicates strength of ASD association; bold indicates genes that are also the representative gene of an ASD-associated module (as in A).

As modules across studies likely contain overlapping genes or capture related latent biological processes, we performed forward stepwise selection to identify a parsimonious subset of modules that jointly and independently explain variation in gene-level ASD RVAS scores (**Methods**). The selected model identified 28 gene modules that collectively best account for ASD RVAS scores, with module size ranging from 14 to 3205 genes (median = 708) and each containing between 4 and 127 high-confidence ASD genes (median = 31.5; **Fig. 3A**; **Table S6**). Results were robust across alternative regression model specifications and ASD-gene thresholds (**Supplement**; **Figs. S8, S9**). Modules are labeled by their representative gene: transcriptional regulons are labeled by the driver TF; co-expression networks are labeled by their “hub” gene, defined as the gene whose expression is most strongly correlated with the overall expression pattern of the module. For regulons, a superscript (+) or (-) suffix denotes whether the TF is predicted to activate or repress its target genes, respectively.

Notably, nearly one-third of the associated modules (8 of 28 total) were centered on a high-confidence ASD gene, either as the TF defining a regulon or as the hub of a co-expression network (**Fig. 3A-B**). This organization suggests that certain ASD genes occupy central positions within broader transcriptional programs that themselves converge on rare variant burden. For instance, *MEF2C* and *ZBTB18* are both high-confidence ASD genes and TFs whose associated developmental regulons exhibited broad enrichment for ASD genetic liability, consistent with previous work ^23,38^. *SOX11*, another risk gene TF driving an associated regulon, has been previously linked to altered A-P cortical patterning in ASD ^12,39^. Co-expression modules with hub genes *SPEN* and *EP300* were also highly enriched and have been previously linked to rare variants in ASD and developmental delay ^24^.

### Modules converge on genomic-regulatory, morphogenesis, and synaptic programs

To identify recurring biological themes across these 28 ASD modules, we next conducted unsupervised topic modeling on associated Gene Ontology (GO) terms. Three topics, hereafter referred to as biological programs, were selected by the tradeoff between semantic coherence and exclusivity (**Fig. 3B**; **Fig. S10**; **Tables S7A, S8A-B**): (1) genomic regulation, including chromatin remodeling and post-transcriptional control (GR), (2) neuronal development and morphogenesis (MORPH), and (3) synaptic transmission and ion transport (SYN). Each of the 28 modules was assigned to its dominant program by consensus across 1000 topic model solutions (**Methods**; **Fig. 3B**). To characterize the cellular contributions of modules within each program, we used AUCell to quantify the proportion of cells with high module activity within each major developmental cell type ^20,40^ (**Methods**; **Fig. 3C**; **Table S7B**).

GR comprised 12 modules enriched for broad genomic regulatory processes spanning epigenetic (e.g., histone and chromatin remodeling), transcriptional, and post-transcriptional (e.g., RNA processing) regulation (**Table S8**). Three were centered on a high-confidence ASD gene: *EP300*, *SPEN*, and *VEZF1* (**Fig. 3B**). GR modules displayed the least cell-type specificity, consistent with the broad expression of gene-regulatory machinery across cell types (**Fig. 3C**). Nevertheless, several GR modules showed prominent activity in developing neuronal populations, including *NRF1^(+)^*, *SREBF2^(+)^*, *VEZF1*, *EP300*, and *RBM27*.

MORPH comprised 11 modules enriched for neuronal development and differentiation, morphogenesis, and axon projection (**Table S8**). Four were centered on a high-confidence ASD gene as the TF driver: *SOX11^(+)^, TCF4^(+)^, ZBTB18^(+)^*, and *FOXP2*^(+)^ (**Fig. 3B**). Like GR, MORPH modules exhibited broad cell-type distributions, especially among immature neuronal populations (**Fig. 3C**). *FOXP2^(+)^* and *ZBTB18^(+)^* were strongly enriched in mature IT ENs.

Finally, SYN comprised 5 modules enriched for processes related to synaptic transmission, ion transport, and vesicle trafficking (**Table S8**), one of which was centered on a high-confidence ASD gene, the *MEF2C^(+)^* regulon (**Fig. 3B**). Compared to MORPH and GR, SYN modules showed the most specific cell type enrichments, as they all demonstrated strong activity in mature neuronal populations, most prominently within the excitatory IT lineage, consistent with the IT-neuron enrichment identified in the scDRS analyses (**Fig. 3C**). ***MEF2C^(+)^*, *PRKCE***, and ***FBXO41*** also demonstrated strong activity in mature interneuron subclasses whereas ***MARCH4*** and ***SRCIN1*** showed more modest interneuron associations. Together, these findings organize ASD-associated modules into three biologically distinct programs with graded cellular specificity: genomic regulation and morphogenesis distributed more broadly across developmental stages and cell types, and synaptic networks concentrated in mature neurons, particularly the IT lineage.

### ASD genes show program-specific developmental timing and spatial signatures

We next sought to characterize the broader developmental timing and regional expression patterns of the ASD-associated genes within each of the three biological programs. This required a gene-level assignment, separate from the unsupervised module assignment above. Each of the 253 high-confidence ASD genes was assigned to its best-matching program based on overlap between the GO terms annotated to that gene and the GO terms enriched in the program’s modules, followed by manual review (**Methods**; **Table S9A**). Because the two assignments were made separately, a module and its representative gene (i.e., hub or TF) were not necessarily in the same program— a divergence we return to below.

These ASD-gene programs showed distinct developmental trajectories in BrainSpan. GR- and MORPH-program ASD genes exhibited significantly higher prenatal expression—a pattern most pronounced for GR (**Methods**; **Fig. 3D-E**). In contrast, SYN genes were more highly expressed postnatally. Consistent with these trajectories, cell-level GR and MORPH activity correlated most strongly with ASD scDRS in immature neuronal populations, whereas SYN activity correlated most strongly in mature neurons (**Methods**; **Fig. S11**); GR activity also remained associated with scDRS in mature neurons, although more weakly than in immature populations.

The three programs also showed distinct and non-uniform spatial organization (**Fig. S12**). GR genes robustly recapitulated the A-P gradient of the full ASD gene set (*ρ* = 0.69; *P*_spin_ = 0.005; **Fig. 3F**), with strongest enrichment in posterior occipital regions (**Supplement**; **Table S10A**), and were not spatially correlated with MORPH or SYN beyond that expected from spatial autocorrelation (**Fig. S13**). This GR pattern was robust across sensitivity analyses, directionally concordant with regional ASD transcriptomic dysregulation from case-control data ^12^, and evident during mid-gestation, when GR genes showed progenitor-biased expression and preferential enrichment in the V1 cortical plate (**Supplement**; **Figs. S14-S16**; **Tables S10-S11**). Given its functional breadth, we further clustered GR-assigned ASD genes by GO-term membership, identifying seven functional groups spanning chromatin regulation, transcription, and post-transcriptional regulation, including RNA processing and protein homeostasis (**Methods**; **Fig. 3G**; **Table S9B**). Thus, GR represents a broad genomic-regulatory program with a prenatal progenitor bias and a persistent posterior cortical signature.

MORPH and SYN comprised a second spatial regime, with strongly correlated regional patterning (*ρ* = 0.85, *P*_spin_ = 2×10^-4^; **Fig. S13**), but, unlike GR, neither exhibiting a significant A-P gradient after accounting for spatial autocorrelation (**Fig. 3F**). Both were most strongly enriched across temporal and parietal association regions and did not show GR’s occipital-predominant enrichment, indicating that the posterior-occipital gradient of the full ASD gene set is largely driven by GR (**Fig. S14**; **Table S10B-C).** This temporal-parietal patterning suggested alignment with the sensorimotor-association (S-A) axis of cortical specialization ^36^. Consistent with this, several MORPH-program ASD genes— including *SATB2, ZBTB18, BCL11A, BHLHE22,* and *RORB* at the sensorimotor pole and *THRB* at the association pole—are themselves established areal-patterning factors (**Fig. S17**; **Table S12**). Their developmental spatial patterns diverged, however, with MORPH genes enriched in germinal zones and SYN genes preferentially localized to the cortical plate and thalamus (**Fig. S16**; **Table S11A-B**). Together, these findings distinguish a posteriorly patterned GR program from MORPH and SYN programs organized along complementary axes of cortical development and specialization.

### De novo mutations implicate specific synaptic compartments

The synapse is a reproducible and long-standing point of convergence in ASD genetics ^5,7^. However, the synapse encompasses a diverse and highly complex set of molecular processes and specialized subcompartments, and ASD genetic associations have not been systematically resolved at this finer scale. Using the expert-curated SynGO ontology framework ^25^, we scored each synaptic compartment for size-adjusted enrichment of damaging *de novo* mutations (DNMs) in ASD probands versus unaffected siblings (*Z*_adj_; **Methods**) and applied Hierarchical HotNet ^41^ to identify connected subnetworks of coordinated enrichment **(Fig. 4A**; **Fig. S18**; **Supplement**; **Table S13A-B**). This analysis identified five subnetworks: three presynaptic (the active zone, presynaptic membrane, and presynaptic ion channels) and two postsynaptic (postsynaptic specialization and postsynaptic organization), indicating that ASD rare-variant burden preferentially localizes to pre- and post-synaptic compartments rather than uniformly across the synapse (**Fig. 4A**; **Fig. S18**). Presynaptic ion channels and the active zone had the highest ASD-gene density, and, consistent with our aggregation approach, enrichment within these subnetworks was broadly distributed across constituent genes rather than driven by a few recurrently mutated ones (**Supplement**; **Fig. S19**). Interestingly, we found that ASD genetic burden was further concentrated in genes with multiple synaptic functions. High-confidence genes such as *GRIN2A*, *GRIN2B*, and *SCN2A* recurred across several subnetworks (**Fig. 4B**), and genes shared across multiple compartments tended to have the highest burden (**Supplement**; **Fig. S21**).

**Figure 4.**
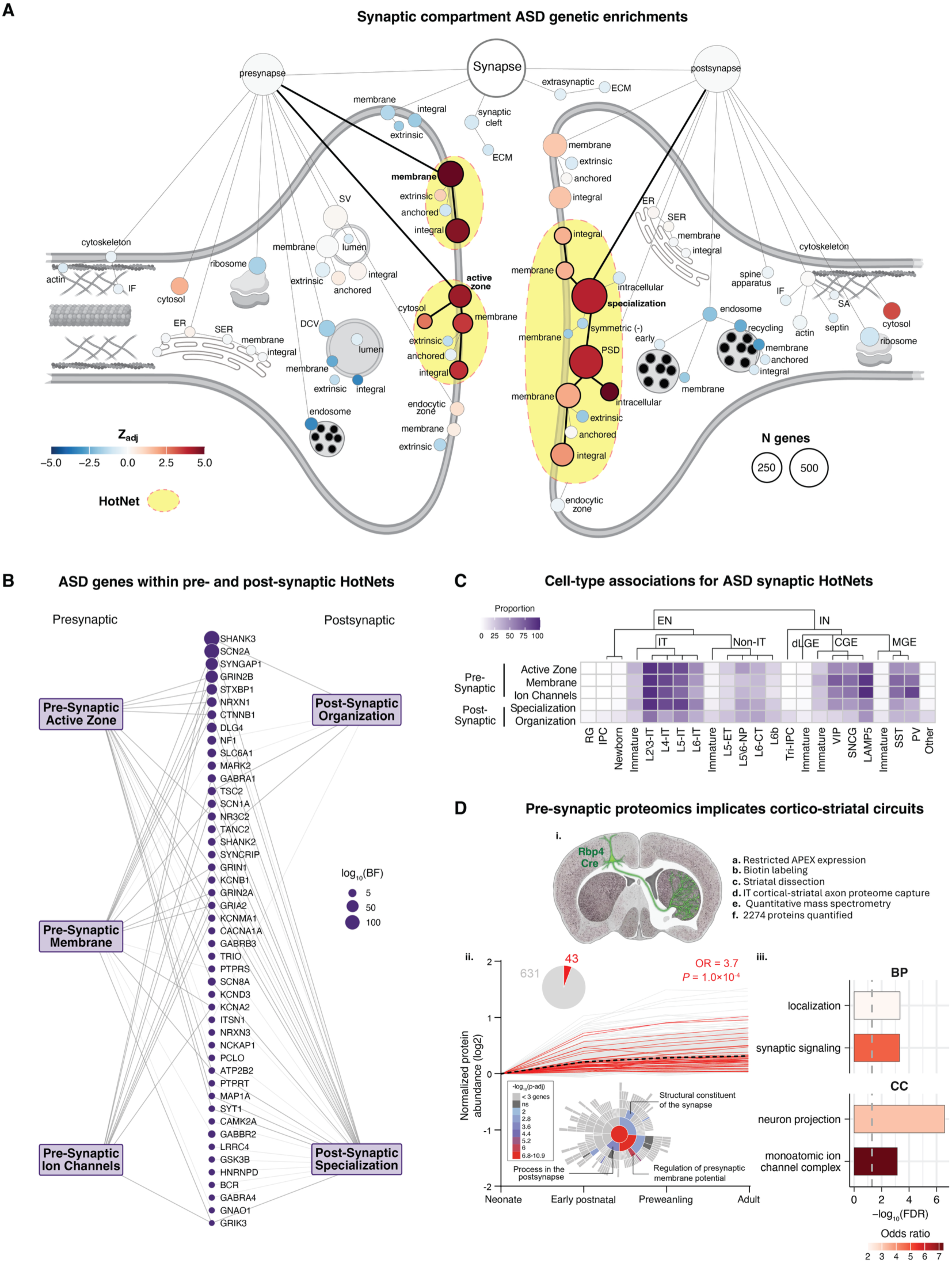
ASD *de novo* mutations cluster within trans-synaptic transmembrane subcompartments. **(A)** Enrichment for damaging DNMs in ASD probands compared to unaffected siblings within the SynGO cellular component (CC) gene ontology network. Each node represents a CC term colored by *Z*_adj_, a gene-count-corrected binomial enrichment statistic comparing damaging DNMs in ASD probands vs unaffected siblings. Node size indicates the number of genes per compartment. Outlined nodes indicate Hierarchical HotNet subnetworks with coordinated elevated signal, including 3 enriched pre-synaptic and 2 post-synaptic compartments. **(B)** SYN ASD genes (TADA FDR < 0.001) organized by SynGO HotNet neighborhood. **(C)** SynGO HotNet cell-type enrichments assessed by AUCell ^40^, highlighting IT neuron and interneuron associations. Tile fill represents the proportion of cells with high AUC scores. **(D)** i. APEX-Cre based approach for capture of IT-enriched cortico-striatal axon and presynaptic proteomes across neurodevelopment in mice using data from ^42^. ii. Co-regulated IT axon proteins showing the greatest increase in expression and axonal localization between neonatal and early postnatal periods are enriched for ASD genes. iii. This IT axonal subproteome is defined by many of the same transynaptic processes implicated in ASD by the analysis reported in Fig. 4A. Abbreviations: DCV = dense core vesicle; ECM = extracellular matrix; (S)ER = (smooth) endoplasmic reticulum; IF = intermediate filament; PSD = postsynaptic density; SA = spectrin-associated; SV = synaptic vesicle.

To characterize the cellular context of each HotNet, we quantified HotNet activity across cell types (AUCell) and tested whether cells with higher activity carried greater ASD genetic vulnerability (as measured by scDRS). HotNet activity increased along the pseudotime trajectory and was enriched among mature EN-IT neurons and, to a lesser extent, mature IN subtypes (**Fig. 4C**; **Fig. S22A-C**; **Supplement**). Presynaptic HotNets (active zone and membrane) most strongly correlated with ASD vulnerability in L4-IT neurons, an association that emerged postnatally and strengthened from infancy to adolescence (**Fig. S23A-B**). In contrast, postsynaptic organization tracked vulnerability in IPC-EN and EN-Newborn subclasses, with these associations strongest prenatally (first and second trimester). Together, these analyses reveal compartment-specific patterns of ASD vulnerability, with presynaptic HotNet programs tracking ASD burden in postnatal mature IT neurons.

These HotNets implicate neurodevelopmentally dynamic, cell-type- and compartment-specific synaptic gene sets in ASD, such as pre- and post-synaptic machinery in mature EN-IT cells, which project to both the cortex and the striatum. To confirm the fidelity of this ASD genetic risk convergence at the spatial and temporal proteome level, we intersected all 253 ASD genes with a recent survey of corticostriatal axonal proteomes across neurodevelopment in mice ^42^. This study, which utilized APEX-Cre-based proximity labeling and quantitative mass spectrometry (**Fig. 4Di**), identified distinct co-regulated protein groups, defined by unique developmental trajectories in corticostriatal axons. A co-regulated sub-proteome (**Methods**) displaying a large increase in axonal protein levels between neonatal and early postnatal periods, followed by leveling during preweaning and adulthood, was significantly enriched for ASD genetic burden (P = 1×10^-4^; **Fig. 4Dii**). In line with the SYN and HotNet analyses above, this developmental axonal sub-proteome was enriched for transsynaptic processes including “Regulation of presynaptic membrane potential”, “Processes in the postsynapse”, and “Synaptic signaling” (**Fig. 4Dii-iii**).

Together, these results indicate that ASD genetic vulnerability is localized to specific synaptic compartments, especially among mature IT neurons, a signature that is reflected in their corticostriatal axonal proteome.

### Neurodevelopmental transcription factors link MORPH to synaptic networks

Synaptic and genomic-regulatory pathways have been the two most reproducible points of convergence for ASD rare variants, but the regulatory relationships between them remain unresolved ^5,7^. The preceding results implicate MORPH as a candidate intermediary: its adult cortical patterning is highly correlated with that of SYN (**Fig. S13**), yet like GR, it is prenatally biased and composed in part of genomic regulators— neurodevelopmental TFs whose MORPH assignment reflects their roles in neuronal differentiation and cortical areal identity (rather than their molecular activity). We therefore asked whether these regulators couple developmental patterning to downstream synaptic machinery. We first tested this model at the network level by asking whether MORPH-associated modules were enriched for synaptic genes. Modules from all 3 programs showed synaptic associations (5 of 5 SYN, 9 of 11 MORPH, and 7 of 12 GR), but enrichment magnitude differed by program (**Fig. 5A**; **Table S13C-D**). As expected from its functional definition, SYN showed the strongest enrichment (OR 4.3, *P* = 6.0×10^-26^). Importantly, MORPH modules were more enriched than GR modules (OR 2.3, *P* = 1.0×10^-6^ vs OR 1.2, *P* = 0.14), despite MORPH not being defined by synaptic annotation. Enrichment for individual pre- and post-synaptic HotNets was more restricted, reaching significance only among SYN modules and preferentially involving presynaptic compartments (**Fig. 5A**). Thus, MORPH extends broadly into synaptic biology, whereas localization to specific synaptic subcompartments is concentrated within SYN.

**Figure 5.**
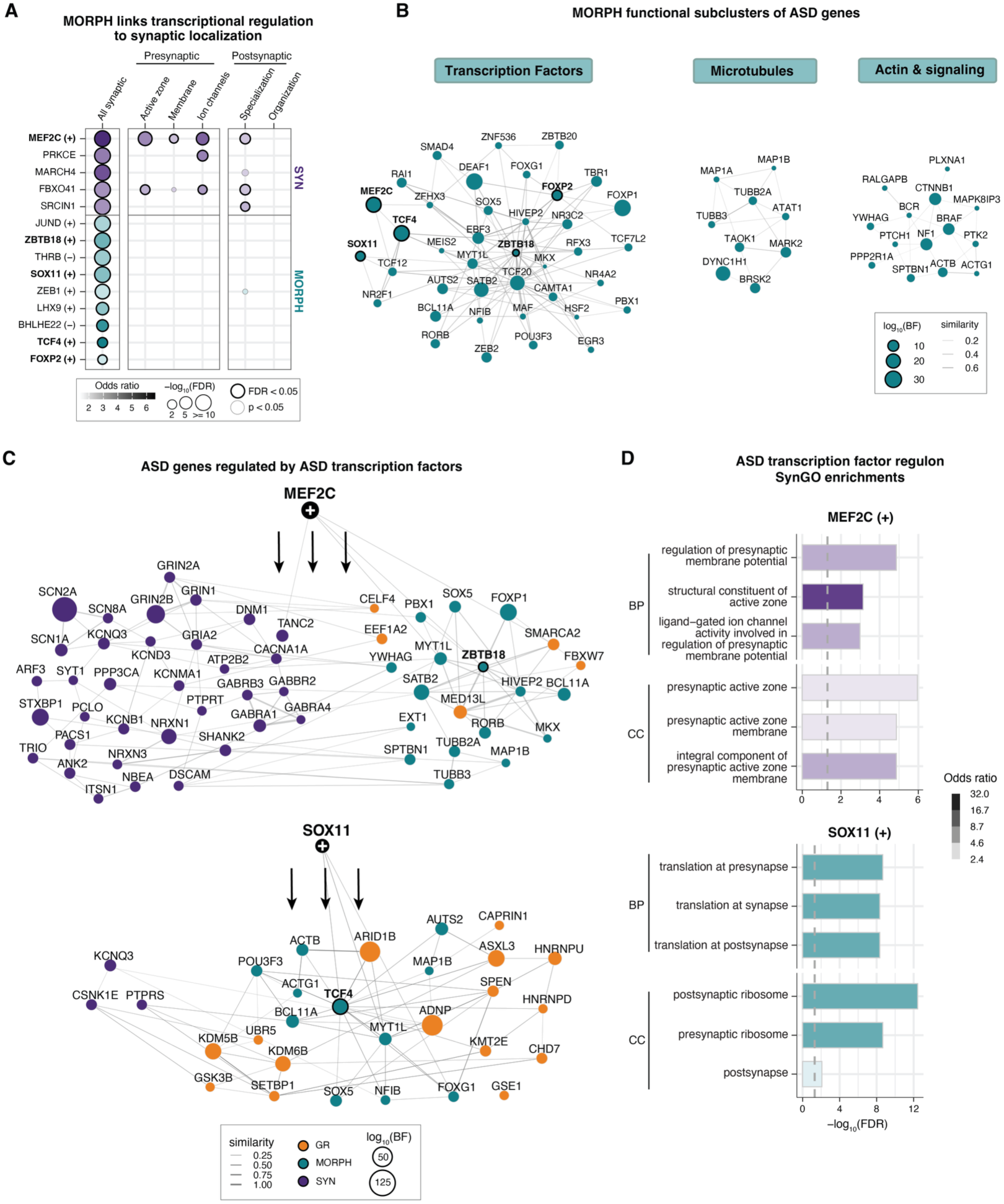
Neurodevelopmental TFs connect ASD genes across neuronal morphogenesis and synaptic networks. **(A)** Enrichment of ASD-associated modules for overall synaptic genes, as well as pre- and postsynaptic SynGO HotNets. Each row represents a module with at least one nominally significant enrichment; circle fill indicates biological program; size encodes odds ratio; opacity indicates -log_10_(FDR). **(B)** Functional subclustering of MORPH-assigned ASD genes (TADA FDR < 0.001; *n* = 69) by GO term membership cosine similarity. Legend as in Fig. 3D. **(C)** Functional similarity among ASD genes in *MEF2C^(+)^* and *SOX11^(+)^*regulons, colored by SYN, MORPH, and GR assignments. Legend as in Fig. 3D, with nodes colored by biological program assignment. **(D)** SynGO enrichments of the *MEF2C^(+)^*and *SOX11^(+)^* regulons within the biological process (BP) and cellular component (CC) ontologies. X-axis indicates - log_10_(FDR); opacity indicates odds ratio.

Many of the 28 ASD-associated modules are TF-driven regulons (*n* = 16), including all 11 in MORPH. In five, the driver TF is itself a high-confidence ASD gene, so these regulons carry two program assignments: one for the driver TF, from its own annotation, and one for the regulon, from the annotation of its targets (**Methods**; **Fig. 3B**). We asked whether any of these five regulons showed divergence between TF- and regulon-level assignments, reasoning that this could indicate direct cross-program connections. *MEF2C*, a high-confidence ASD gene, provided the clearest example. The *MEF2C*^(+)^ regulon was assigned to SYN and showed the strongest enrichment across presynaptic HotNets of any ASD-associated module (**Fig. 5A**), whereas *MEF2C* itself is a MORPH- assigned TF with peak brain expression during mid-fetal stages ^23^. This positions *MEF2C* as a potential common link between developmental transcriptional regulation and synaptic functions. We therefore asked whether *MEF2C* belongs to a broader transcription-factor branch of MORPH. Applying the same GO-based clustering used above resolved four MORPH subclusters: neurodevelopmental transcription factors and chromatin regulators, microtubule cytoskeleton and neuron polarity, cytoskeletal signaling and axon guidance, and a residual low-annotation group (**Fig. 5B**; **Table S9C**). *MEF2C* clustered with other neurodevelopmental regulators, including *SOX11*, *ZBTB18*, *FOXP2*, and *TCF4* (**Fig. 5B**; **Fig. S24**), identifying a subset of MORPH-assigned ASD genes positioned to regulate downstream developmental programs.

To define the downstream architecture of this connection, we examined these neurodevelopmental regulators and their targets in greater detail. The *MEF2C^(+)^* regulon contains 56 high-confidence ASD genes among its 765 targets, including *SCN2A*, *STXBP1*, *GRIN2A*, and *CACNA1A*; of these 34 are assigned to SYN, 17 to MORPH, and 5 to GR (**Fig. 5C**). The SYN-assigned targets map to an interconnected protein-protein interaction network with other regulon members (**Fig. S25**), and the regulon itself showed SynGO enrichments for active zone organization and presynaptic ion channel activity (**Fig. 5D**). *SOX11* provides a complementary example. Both *SOX11* and its regulon are assigned to MORPH, yet its downstream targets also extend into synaptic biology, with enrichment centered on synaptic translational machinery (**Fig. 5D**)—demonstrating that MORPH transcription factors can reach the synapse through distinct functional routes. Collectively, these results position MORPH’s transcription factors as a putative link between GR and SYN, as nuclear transcriptional regulators whose downstream targets converge on synaptic architecture and function.

### Developmental disorder comorbidity stratifies ASD-associated cellular and developmental contexts

As many high-confidence ASD genes are also associated with DD/ID ^2,8,43^, we next asked whether their cellular and developmental enrichment patterns differ between ASD probands with recorded DD/ID (ASD_DDID_, abbreviated to ASD_D_) and those without (ASD_DDID-Absent_, abbreviated to ASD_DA_). We stratified high-confidence ASD genes with at least one damaging DNM (defined as PTV, Mis2, and DEL; **Methods**) among all probands (*N* = 38,680) into three groups based on their mutation burden in ASD_D_ versus ASD_DA_ probands: ASD_DA_, ASD_DH_ (ASD_DDID-High_), and ASD_DL_ (ASD_DDID-Low_) genes. ASD_DA_ genes had a higher per-proband mutation rate in ASD_DA_ than ASD_D_ probands (*n* = 44); the remaining genes were split by whether the fraction of damaging DNMs from ASD_D_ probands was above (ASD_DH_; *n* = 121) or below (ASD_DL_; *n* = 84) the background expectation (p0) (**Methods**; **Fig. 6A-B**; **Table S14A**). As the ASD_DA_ group was relatively small, we first analyzed it together with ASD_DL_ genes (*n* = 128), then restricted the analysis to ASD_DA_ genes to identify signals specific to ASD without DD/ID.

**Figure 6.**
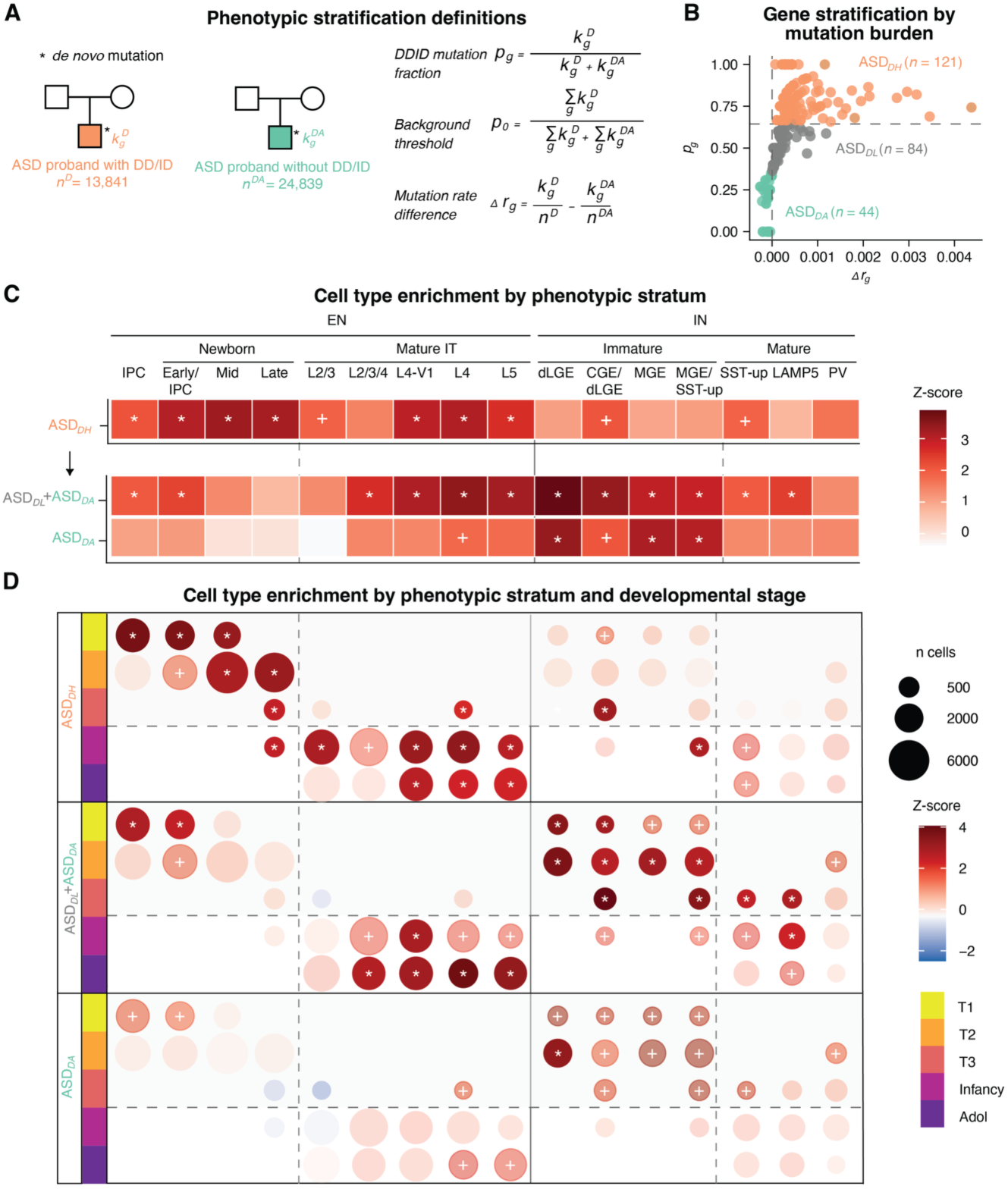
ASD genetic burden across cell types and developmental stages differs by DD/ID comorbidity status. **(A)** Phenotypic stratification framework for DD/ID comorbidity analysis. ASD probands are classified by presence (ASD_D_, *n* = 13,841) or absence (ASD_DA_, *n* = 24,839) of comorbid developmental delay or intellectual disability. For each gene *g*, 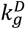 and 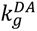 denote the observed *de novo* mutation counts in each cohort; *p* quantifies the relative mutation burden in ASD_D_ probands; *p*_0_ is the cohort-wide background expectation; Δ*r_g_* captures the per-proband mutation rate excess in ASD_D_ relative to ASD_DA_. **(B)** Gene stratification by DD/ID mutation burden. X-axis shows mutation rate difference (*Δr_g_*); y-axis shows mutation fraction (*p*_g_) for high-confidence ASD genes (TADA FDR < 0.001; *n* = 253). The horizontal dashed line indicates *p*_0_; genes are colored by stratum. **(C)** Cell type enrichment by phenotypic stratum. Color indicates scDRS enrichment *Z*-scores for ASD_DH_, ASD_DL_ and ASD_DA_ combined, and ASD_DA_ gene sets across lineage clusters with significant ASD enrichment (Fig. 1D; FDR < 0.05). **(D)** scDRS enrichment *Z*-scores across developmental stages and lineage clusters for each gene stratum. Each dot represents a cluster-stage combination with at least 150 cells; size encodes the number of cells; color encodes the enrichment *Z*-score. The horizontal dashed line separates prenatal from postnatal stages; vertical lines separate EN and IN lineages and subtypes as in (C). T1, First trimester; T2, Second trimester; T3, Third trimester; Adol, Adolescence. Significance: * FDR < 0.05; ^+^ *P* < 0.05, uncorrected. Nominally significant results are shown with reduced opacity, non-significant results are displayed with the lowest opacity.

Among lineage clusters with significant enrichment identified using scDRS (**Fig. 1D**, FDR < 0.05), mid- and late-pseudotime newborn EN subpopulations were specifically enriched in ASD_DH_ genes. ASD_DH_ and the combined ASD_DL_ and ASD_DA_ set showed largely overlapping EN-IT enrichment, with ASD_DH_ genes showing less IN enrichment. (**Fig. 6C**; **Table S14B**). For ASD_DA_ genes alone, EN-IT cells demonstrated weaker enrichment, likely reflecting both attenuation of ASD_DL_ gene signals and reduced statistical power from the smaller gene set. In contrast, the ASD_DA_ gene set alone demonstrated immature IN enrichment: dLGE-immature *Z*-scores remained high (3.89 to 3.38) and MGE-immature *Z*-scores rose (2.89 to 3.05; 2.96 to 3.15). This pattern was replicated when the TADA FDR threshold for defining ASD_DA_ genes was relaxed to FDR < 0.01 (*n* = 105 genes; **Fig. S26**).

We next asked if these cell type enrichments were specific to developmental stages by quantifying the cell-type enrichments across trimesters 1-3, infancy, and adolescence. ASD_DH_ genes were enriched in prenatal ENs across all trimesters and in L4-IT neurons across both prenatal and postnatal stages (**Fig. 6D**; **Table S14C**). The combined ASD_DL_ and ASD_DA_ gene set showed predominant enrichment in prenatal INs (second and third trimesters), with L4-IT enrichment restricted to postnatal stages. ASD_DA_ genes retained significant enrichment in prenatal dLGE-immature INs while EN enrichment remained nominal.

Given that the cell-type enrichment analyses above were limited to cortical cells, we additionally examined DNM burden (here defined as PTV, Mis1, or Mis2 for gene set enrichment analysis; **Method**; **Supplement**) in germinal zone (GZ), cortical plate (CP), and thalamus (THL) gene sets from a prenatal spatial transcriptomic atlas ^18^, the latter representing a major source of thalamocortical input ^28^. In each region, ASD_D_ probands demonstrated higher DNM burden than ASD_DA_ probands, and CP- and THL-enriched genes showed significantly higher burden than GZ-enriched genes (**Fig. S27A-D**; **Table S15**). In contrast, no regional differences were observed in ASD_DA_ probands (**Supplement**).

Finally, we explored whether ASD-associated modules and SynGO HotNets demonstrated differential enrichment between ASD_D_ and ASD_DA_ probands (**Supplement**). As with the spatial gene sets, almost all modules and pathways were enriched in both ASD_D_ and ASD_DA_ probands relative to siblings, with rare variant burden broadly elevated in ASD_D_ probands (**Figs. S28A-D,S29A-B**; **Tables S16-S17**). The only exception was synapse adhesion between pre- and postsynapse (GO:0099560), enriched in ASD_DA_ but not ASD_D_ probands relative to siblings (**Fig. S29A-B**; **Table S17A-B**), with mutations concentrated in *NRXN1* (12 DNMs), *PTPRS* (5), *PTPRD* (5), *LRFN4* (4), and *LRRC4* (4). Notably, *PTPRD* and *PTPRS* encode presynaptic proteins that interact directly with the postsynaptic adhesion molecule *LRFN4* ^44^, suggesting amplified vulnerability across synaptic connections. These findings indicate that trans-synaptic adhesion may represent a distinct axis of ASD vulnerability independent of DD/ID comorbidity.

Together, these findings indicate that the biological programs disrupted by rare variants in ASD are broadly shared across the phenotypic spectrum, although the cell-type and developmental contexts in which they operate differ by DD/ID comorbidity status.

## Discussion

The findings presented here refine our understanding of ASD rare-variant vulnerability beyond the cell types and broad molecular pathways established by prior exome studies ^2,5,7,21^. Integrating the largest exome sequencing dataset to date with developmental single-cell and spatial atlases, regulatory networks, and synaptic ontologies, we resolved this convergence into three biological programs—genomic regulation (GR), neuronal morphogenesis (MORPH), and synaptic transmission (SYN)—with distinct developmental, cellular, and spatial signatures. Genetic liability was concentrated in newborn excitatory neurons, immature interneurons, and mature intratelencephalic (IT) neurons, and, within the synapse, in the pre- and post-synaptic membrane-associated machinery. Genes associated with ASD were also spatially patterned across the cortex, following an anterior-to-posterior gradient anchored at the primary visual cortex in the occipital pole, and driven predominately by GR. Finally, these patterns differed with DD/ID comorbidity, indicating that broadly shared molecular programs can confer ASD liability through distinct cellular and developmental contexts.

Previous co-expression network analyses have consistently implicated mid-fetal cortical excitatory neurons in ASD vulnerability ^7,10,11,13,14^. Our data both confirm and extend these prior results. Rare-variant association first peaked around the transition from intermediate progenitors to newborn excitatory neurons—consistent with this fetal signal—but resolved into a bimodal trajectory with a second peak in mature IT neurons, while interneuron vulnerability concentrated in immature populations. The postnatal IT signal coincides with the prolonged maturation and experience-dependent refinement of intracortical excitatory circuits. Consistent with a synaptic locus for this later vulnerability, damaging DNMs concentrated in presynaptic active zone and membrane compartments and postsynaptic specializations rather than uniformly across the synapse, refining the synaptic hypothesis of ASD ^45–47^ toward the machinery of synaptic signaling across pre- and postsynaptic membranes. This interpretation is further supported by enrichment of ASD genes within a developmentally regulated corticostriatal axonal proteome in the mouse that rises sharply during early postnatal maturation ^42^. The apparent disagreement of earlier studies over deep-versus superficial-layer vulnerability also becomes more coherent when reframed by projection identity: enrichment favored IT neurons across cortical layers 2-5 over ET, CT, and NP lineages. Because IT neurons mediate broad cortico-cortical, cross-callosal, and cortico-striatal connectivity ^28,29^, these findings implicate circuits supporting communication within the telencephalon rather than a single cortical layer. Enrichment was strongest in L4-IT neurons, the principal recipients of thalamocortical input and a population particularly prominent in primary sensory cortices ^16,17,20,36^.

Phenotypic stratification further indicates that these cellular contexts are not uniform across ASD. Genes carrying greater mutation burden in probands with DD/ID showed broader excitatory-neuron involvement, including prenatal and newborn excitatory populations, whereas genes with higher per-proband mutation rates in ASD without DD/ID retained prominent enrichment in immature interneurons. Most regulatory modules and synaptic pathways remained enriched in both groups, arguing against a simple partition of ASD into distinct molecular etiologies by DD/ID comorbidity; rather, shared biological programs appear to differ in the developmental stages and neuronal populations through which genetic vulnerability is expressed. Selective enrichment of trans-synaptic adhesion in ASD without DD/ID provides a more pointed exception, nominating synaptic connectivity itself as one axis of vulnerability less tightly coupled to broader developmental impairment. These distinctions remain provisional, given the smaller ASD-without-DD/ID gene set and the heterogeneity inherent in clinical DD/ID ascertainment, but they illustrate how growing genetic sample sizes can begin to connect molecular convergence with phenotypic variation.

The developmental and spatial signatures of the three programs suggest a working model linking ASD genetic vulnerability to successive phases of cortical arealization. GR showed the strongest prenatal bias and was enriched in germinal-zone progenitors, where molecular morphogen gradients contribute to the cell-intrinsic cortical protomap ^15,16,48^. GR also accounted for most of the posterior A-P gradient in the adult cortex and showed preferential developmental enrichment toward V1, the most molecularly distinct area of the neocortex ^17,49^. Together, these observations position the GR-program chromatin remodeling and transcriptional machinery as a candidate regulatory substrate linking early cortical patterning to the persistent molecular specialization of posterior sensory cortex. Consistent with this, the GR regulon *CUX1^(+)^*, a predicted driver of ASD-associated transcriptomic downregulation ^14^, marks V1-specific L4-IT neurons in the developing cortex ^20^. SYN showed a complementary profile: postnatally biased expression, developmental localization to the cortical plate and thalamus, and preferential activity in mature IT neurons. Together, these features are consistent with a role in the later assembly and activity-dependent refinement of cortical circuitry envisioned by the protocortex model ^50^, although its adult spatial enrichment extended predominantly across temporal and parietal association cortices. V1 may therefore represent a particularly sensitive point of convergence between these developmental mechanisms: it occupies the posterior extreme of the intrinsic A-P protomap while also representing a primary sensory area strongly shaped by first-order thalamocortical input. Its prominent ASD genetic association may thus reflect vulnerability at the intersection of intrinsic areal specification and extrinsic circuit refinement. More broadly, these results suggest that ASD genetic risk intersects both early intrinsic patterning and later activity-dependent circuit maturation rather than implicating a single developmental epoch.

MORPH provides a potential connection between these regimes. Like GR, MORPH genes were prenatally biased and enriched in developmental cell populations, yet their adult spatial pattern resembled SYN’s. MORPH also contains neurodevelopmental transcription factors whose downstream programs extend into synaptic biology. This link is exemplified by *MEF2C:* although the transcription factor is itself assigned to MORPH, its regulon is strongly enriched for SYN genes and presynaptic functions, including active-zone organization and ion-channel activity. More broadly, MORPH contains several established areal-patterning transcription factors spanning the sensorimotor and association poles (**Table S12**), while MORPH regulons as a group extend substantially into synaptic biology. This architecture—developmental regulators of neuronal identity whose downstream programs reach the synapse—may link early neuronal differentiation to later areal and synaptic specialization. We speculate that it participates in the transcriptional response to extrinsic thalamocortical signals described by induction-exclusion models of cortical development ^36^, although direct regulation by thalamic input remains to be established. In this framework, GR, MORPH, and SYN represent overlapping stages rather than strictly independent pathways: early genomic regulation, neuronal differentiation and morphogenesis, and the subsequent assembly and refinement of synaptic circuitry.

Several limitations qualify this framework. These analyses are primarily associational and integrate cross-sectional human genomic and postmortem molecular data; the developmental trajectories therefore represent inferred rather than directly observed longitudinal processes; and these atlases sample a limited number of donors and cortical regions. The gene-level analyses aggregate rare-variant classes with potentially distinct functional consequences, and DD/ID status compresses substantial phenotypic heterogeneity into a binary clinical classification. Most importantly, the proposed directional relationships—including GR contributing to the establishment or maintenance of posterior cortical identity and MORPH coupling developmental patterning to synaptic programs—require experimental validation in model systems. Nevertheless, convergence across genetic association, developmental cell states, cortical patterning, regulatory networks, and synaptic compartments provides a framework for testing how ASD rare-variant vulnerability propagates from early molecular regulation through neuronal differentiation to the assembly and refinement of cortical circuits.

## Data availability

Rare variant association results are available as supplementary tables in ^8^. Specifically, gene-level counts for ASD probands are in Table S7, TADA results are in Table S9, and gene-level counts for DDID probands are in Table S17. The present study analyzed the following publicly available datasets: developing snMultiome human brain atlas (https://doi.org/10.5061/dryad.2280gb612); μBrain laser-microdissection microarray of the mid-fetal human cortex (https://zenodo.org/records/10622337); prenatal Visium spatial transcriptome data (https://zenodo.org/records/14422018); pre-synaptic mass spectrometry data (https://www.ebi.ac.uk/pride/archive?keyword=PXD030864), and the SynGO database (v1.2, https://www.syngoportal.org/). Gene assignments to each of the co-expression modules, transcription regulons, and prenatal spatial transcriptomic factor-enriched gene sets analyzed were accessed from the supplementary material of their respective studies. GWAS summary statistics of ASD was downloaded from: https://bitbucket.org/steinlabunc/spark_asd_sumstats.

## Code availability

Scripts for analysis described in this manuscript are available on GitHub at: https://github.com/gandallab/asd_rarevar_anno

## Acknowledgements

This research was supported, in part, by grants from the Simons Foundation Autism Research Initiative (SF1018804 to M.L.M., B.D., and K.R.), the National Science Foundation (NSF RECODE 2225624 to M.J.G.), and the National Institutes of Health (R01MH129725 and R01MH123184 to K.R.; R01MH137578, R01MH123922, and R01MH121521 to M.J.G.; R01MH143222 to M.L.M. and M.J.G.; R01MH125235 to M.L.M. and B.D.). R.L.S. was supported by the National Human Genome Research Institute of the National Institutes of Health under Award Number T32 HG009495. The content is solely the responsibility of the authors and does not necessarily represent the official views of the National Institutes of Health.

## Author contributions

M.J.G., B.D. and K.R. supervised the study. M.J.G., B.D., K.R., and M.L.M. acquired funding. M.J.G., B.D., R.L.S. and Y.L. conceptualized the study. R.L.S. and Y.L. performed the primary analyses, developed the methodology, created the figures and wrote the manuscript, with M.J.G. contributing to analysis, methodology, visualization, and writing throughout. B.D. and L.K. performed the SynGO analyses. K.R. and L.Z. performed the HotNet analysis. M.L.M. performed the proteomics analysis. B.D., K.R., M.L.M. and Y.K. contributed to visualization and writing and provided expert input on interpretation of results throughout. Y.M. and M.T. contributed to methodological discussion and interpretation of results. F.K.S., C.A. and J.M.F. curated the ASC genetic data; C.A. also revised the manuscript. J.D.B., M.J.D. and M.E.T. contributed the ASC cohort data and revised the manuscript. L.d.l.T.-U. provided expert input and revised the manuscript. All authors reviewed and approved the final manuscript.

## Methods

### Definition of the ASD rare variant gene set

We utilized the ASD rare variant association results from 62,429 individuals with recorded autism (38,680 probands and 23,749 cases) and 33,316 individuals without recorded autism (9,567 siblings and 23,749 controls) derived from the aggregation of multiple fundamental research and clinical datasets described in our companion genetic association study ^8^. Rare variant associations of protein-truncating variants (PTVs), missense variants with MPC ≥ 2 and AlphaMissense estimated pathogenicity (AM path) ≥ 0.97 (Mis2), and missense variants with either MPC ≥ 2 or AM path ≥ 0 (Mis1), deletions (DELs), and duplications were jointly modelled by the Transmission and *De Novo* Association (TADA) Bayesian framework ^2,51^ to obtain the gene-level associations ^8^. High-confidence ASD genes were defined at a false discovery rate (FDR) threshold of < 0.001 (*n* = 253). This gene set, together with the continuous gene-level associations, forms the basis of all downstream analyses (**Figs. 1-5**). Sensitivity analyses used expanded gene sets at relaxed TADA thresholds: FDR < 0.01 (*n* = 416), < 0.05 (*n* = 696), and < 0.1 (*n* = 951).

### Single-cell disease relevance score framework

To investigate the cellular manifestation of ASD genetics across human brain development, we utilized a single-nucleus multiome (snMultiome) atlas from 38 human neurotypical samples covering developmental stages across the first, second, and third trimesters, infancy, and adolescence ^20^. The released processed data comprised 232,328 quality-controlled nuclei profiled by paired snRNA-seq and snATAC-seq together with curated subclass and cell-type annotations, and associated metadata including sample identity, age, sex, developmental stage, and brain region. For visualization of the complete snMultiome atlas, we used the precomputed weighted nearest-neighbor (WNN)-based uniform manifold approximation and projection (UMAP) coordinates (**Fig. 1B**).

We mapped genetic signals to individual cells using the scDRS (single-cell Disease Relevance Score) framework ^52^ to evaluate both rare and common variant contributions. In the context of ASD, we refer to the resulting scDRS as Condition Relevance Score. For rare variant association study (RVAS)-based scDRS computation, the input gene-level *P* values of the 253 high-confidence ASD genes were transformed from FDR values using the formula *P* = FDR*rank(FDR) / (max(FDR)*length(FDR)). We also performed sensitivity analyses by applying the same *P*-value transformation to the expanded gene sets. For common variant-based scDRS computation, the input gene-level *Z*-scores were derived from ASD GWAS summary statistics ^31^ using MAGMA ^53^. A total of 61 genes reached gene-level significance in ASD GWAS at FDR < 0.05, and 127 genes at FDR < 0.1. Because scDRS requires a minimum input gene set size of 100 genes, we used the top 100 genes ranked by MAGMA *Z*-score for the primary analysis and the top 200 genes for sensitivity analysis.

Individual cell raw scDRSs were computed based on the aggregate weighted expression of these prioritized gene sets. To provide a statistical baseline, scDRS computed 1,000 Monte Carlo (MC) samples of raw control scores for each cell, each derived from a control gene set matched to the input gene set in size, mean expression, and expression variance. Raw scores were adjusted for technical and biological covariates, including sample ID, sex, and number of genes detected per cell. Cell-level p-values were estimated by normalizing raw scores through gene set alignment and cell-wise standardization, then comparing each cell’s normalized scDRS against the empirical distribution of pooled normalized control scores across all cells and all 1,000 control gene sets ^52^. To identify key cell populations, we performed downstream group-level analysis at the subclass (**Fig. 1C**) and cluster (**Fig. 1D**) levels. For group-level association testing, we applied the scDRS downstream analysis using a Monte Carlo (MC) test in which the top 5% quantile of normalized scDRSs of cells within each group was used as the test statistic. Multiple-testing-corrected MC *P* values (assoc_mcp) were used to determine statistical significance, and MC *Z*-scores (assoc_mcz) were used to further prioritize associations whose MC *P* values reached the MC limit of 1/(1 + 1000). All analyses were performed on the RNA assay data that were size-factor normalized and transformed as ln(1 + normalized counts) using the *LogNormalize* method in *Seurat* (v5.3.1) ^54^. scDRS analyses were performed using the default parameters of *compute-score* and *perform-downstream* scDRS functions. These analyses produced the subclass- and cluster-level ASD enrichment results in **Fig. 1C-D**, with the common variant and expanded gene set sensitivity analyses in **Fig. S5**.

### Developmental trajectory inference

To reconstruct the developmental trajectories of excitatory (EN) and inhibitory (IN) neurons, we subsetted relevant cells from the whole dataset. For the EN lineage, we included radial glia (RG), intermediate progenitor cells (IPC-ENs), and glutamatergic neurons. For the IN lineage, we included RG, Tri-IPC (IPC-Glia), and GABAergic neurons. For each lineage, we reconstructed a weighted nearest-neighbour (WNN) graph using *Seurat* (v5.3.1) to integrate 1-50 principal components and 2-40 latent semantic indexing (LSI) components, and used this graph to generate an eight-dimensional (8D) UMAP embedding. Clustering was performed in this 8D space using *mclust* R package (v6.1.2) ^55^, yielding 23 EN-lineage clusters and 21 IN-lineage clusters after removal of outlier populations. For each lineage cluster, we calculated the proportion of cells assigned to each snMultiome atlas curated cell-type annotation ^20^. Clusters in which a single type accounted for >70% of cells were named accordingly; the remainder were assigned a combined label listing all constituent types (**Fig. S1A**). These composition-based labels were further examined using cluster-average expression of marker genes of snMultiome atlas curated cell types. Clusters assigned a combined label co-expressed marker genes corresponding to each constituent type, consistent with the proportion-based assignment (**Fig. S1B**). Consistent with findings of the snMultiome atlas ^20^, EN-L4-IT neurons diverged into V1-specific and shared (PFC and V1) populations based on their region of origin.

We modeled developmental trajectories using *Slingshot* (v2.7.0) ^56^. A global lineage structure was first established via a cluster-based Minimum Spanning Tree (MST), designating RG-vRG as the root (starting) cluster and terminally differentiated neurons as leaves (ending clusters). We then fitted nine simultaneous principal curves to the MST for each lineage. Individual cell weights and pseudotime values were calculated based on orthogonal projections onto these curves. Branch-specific shrinkage was applied to the curves for better convergence. Cell-level pseudotime values were then visualized on a 2D UMAP embedding generated from the same WNN graph.

To investigate the temporal progression of genetic liability, we mapped scDRS onto the inferred trajectories. For cluster-level association, we performed scDRS group-level analysis on the *mclust*-defined clusters (**Fig. 1D**). Resulting p-values were adjusted for multiple testing across all evaluated clusters. Cells were further partitioned into 50 equal-width bins along the pseudotime axis. The EN lineage occupied all 50 bins, whereas the IN pseudotime distribution contained a region with no observed cells, resulting in only 41 occupied bins. We calculated the average scDRS per bin for visualization, and performed scDRS group-level analysis treating each bin as a group to test for enrichment (**Fig. 1E**). These analyses produced the cluster- and pseudotime-resolved enrichment results in **Fig. 1D-E**, with supporting analyses in **Figs. S1-S2** and **S5-S6** (**Tables S1,S4**).

### Spatial transcriptomic ASD genetic enrichment

To assess the spatial enrichment of the 253 high-confidence ASD genes (244 of which were present in the spatial feature space), we analyzed one 10x Visium section from GW20 human occipital cortex (V1 and V2, 3,591 spots) ^17^, using the scDRS framework. We compute per-spot condition relevance scores from raw counts, with the number of detected genes per spot as a covariate. scDRS internally performed library-size normalization and log-transformation, with 1,000 matched control gene sets defining the null distribution per spot. Occipital layer annotations were derived from the original cluster-to-layer mapping over Louvain clusters (resolution 1.5), with the V1-specific L2/L3 subdivision collapsed into a single upper-plate band, giving 6 germinal layers (not region-split) and 6 cortical-plate layer-by-region (V1/V2) combinations. ASD genetic group-level enrichment was tested for each of the resulting 12 groups using the same Monte Carlo test as above, with FDR correction across all 12 groups.

### Adult cortical enrichment of gene sets

Regional gene expression data with associated spatial coordinates were derived from the Allen Human Brain Atlas (AHBA) adult post-mortem microarray dataset ^30^ using abagen v0.1.3 ^57^. Five of the six total AHBA donors were retrievable; these data were processed through abagen’s standard workflow: differential-stability probe-to-gene selection, intensity-based filtering, assignment of tissue samples to the volumetric Desikan-Killiany (DK) atlas ^37^, within-donor scaled normalization, and cross-donor aggregation ^58^. This resulted in a matrix of 68 cortical (plus subcortical and brainstem regions) x 15563 genes; left- and right-hemisphere homologues were then averaged to a single value per region (resulting in *N* = 34 cortical regions). The brain-expressed gene set was defined as all genes with mean cross-regional expression at or above the 10th percentile (*n* = 14,006).

To test for spatial enrichment patterns of a given gene set, each gene’s expression was first *Z*-scored across regions. The per-region enrichment score of the gene set was calculated as the mean standardized expression of its members (restricted to the background; *n* = 239 of 253 ASD genes total). To quantify the per-region enrichment above chance, each region’s observed score was compared to a null distribution built from 10000 random size-matched gene sets drawn without replacement from the background, and expressed as a *Z*-score relative to that null (*Z* vs background = [observed - null mean] / null SD). Statistical significance was assessed as the empirical one-sided (enrichment) *P* and corrected across regions using the Benjamini-Hochberg method.

We additionally tested whether enrichment was organized along the cortical anterior-posterior axis by correlating (Spearman’s ρ) each region’s enrichment *Z* with its mean anterior-posterior coordinate on the fsaverage surface ^59^. Anterior-posterior position was defined as each region’s mean y-axis coordinate of its vertices on the fsaverage pial surface. Significance was assessed using a null model that preserves spatial autocorrelation ^60^ by generating parcel-wise spin permutations (*n* = 5000) using the Cornblath method ^61,62^, leveraging neuromaps tools ^63^ at 41k fsaverage density, following the implementation of ^64^. The spin *P* value was the proportion of rotated maps with |*ρ*| >= the observed value. We used this same spin permutation framework to identify individual regions (and the occipital cluster, *N* = 4 DK regions [pericalcarine, lateraloccipital, lingual, cuneus]) whose enrichment exceeded the spatially rotated null. These analyses produced the regional enrichment maps and anterior-posterior gradients for the full ASD gene set (**Fig. 1G**) and for each functional program (**Fig. 3F**), with additional sensitivity analyses reported in the **Supplement**. Details on the gene set spatial enrichment in the developing cortex (μbrain analysis ^19^) can also be found in the **Supplement**.

### Gene module curation and conditional association testing

To identify biological mechanisms potentially impacted by rare genetic variation in ASD, we curated a comprehensive set of gene co-expression modules and transcriptional regulons (collectively referred to as “modules”) from four independent sources: (1) developmental gene- and isoform-level co-expression modules (*n* = 124 total) ^24^; (2) ASD-associated pancortical gene- and isoform-level co-expression modules (*n* = 36 gene-level, *n* = 62 isoform-level) ^12^; (3) development regulons (*n* = 583) ^20^; and (4) ASD-associated regulons (*n* = 346) ^14^. This resulted in a total of 1,151 modules. All analyses were restricted to the 11,324 genes shared across all four studies and the present study.

To test each module for an individual association with ASD, we fit a linear regression model (**Equation 1**) in which the outcome variable (*Z*_ASD_) was defined as the gene-level *Z*-score derived from the TADA FDR-transformed *P* value, and module membership (*M*) was a binary predictor (0 = not in module, 1 = in module). Standardized transcript length (the sum of all non-overlapping exons per gene) and its log_10_ transform was included as a covariate to control for gene-length bias. All *P* values were corrected for multiple comparisons using the Benjamini-Hochberg (BH) method.

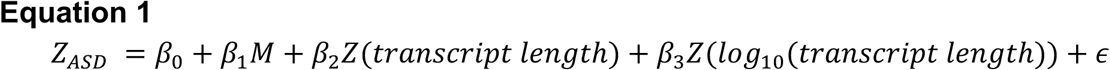

Because many modules share substantial gene content and thus represent redundant biological information, we used conditional forward selection to identify a parsimonious set of significantly associated modules which carry independent information. For this analysis, we included all modules that were significantly marginally associated at FDR < 0.001 as candidates, a stringent threshold to prioritize high-confidence modules given the large number of tests; the final selected set was robust to this choice (**Fig. S8**). Beginning with the most strongly associated module (lowest marginal *P* value), we iteratively tested all remaining candidate modules by re-running **Equation 1**, conditioning on all previously selected modules as covariates. At each step, the module that most improved model fit (as assessed by a likelihood ratio test) was retained and added as a covariate for the next iteration. Selection continued until no remaining module improved model fit (conditional likelihood-ratio *P* < 0.001, a more stringent threshold to limit false inclusions from the repeated testing inherent to forward selection). Modules that survived this conditional forward selection and were individually associated (FDR < 0.001) were carried forward into enrichment testing (**Fig. 3A**); marginal association statistics for all modules are provided in **Table S6B**. Co-expression modules are referenced in the text by their top hub gene (highest module eigengene correlation, kME ^65^); refer to **Table S6A** for module labels used here and module names from the original studies.

To confirm that results were not sensitive to the use of a continuous TADA score outcome or to potential enrichment of DDID-associated genes at the FDR < 0.001 threshold, we repeated the analysis using logistic regression with a binary ASD gene outcome across four TADA FDR thresholds (FDR < 0.001, 0.01, 0.05, and 0.1).

Together, these analyses identified the significantly associated modules shown in **Fig. 3A** (**Tables S6A-B**); threshold-robustness and continuous-versus-binary sensitivity analyses are reported in **Figs. S8** and **S9A-B**.

### Functional annotations of ASD-associated modules

#### Biological pathways and topic modeling

To characterize the biological processes captured by the ASD-associated modules, we performed overrepresentation analyses using Gene Ontology (GO) terms ^66^, restricted to terms annotating between 5 and 1000 genes. Overrepresentation was assessed by Fisher’s exact test (one-sided) against a background of GO-annotated genes within each module’s study network, with BH correction across terms within each module. The same approach was applied to select modules to test SynGO ^25^ term enrichment. To synthesize GO results across modules, we applied unsupervised topic modeling ^67^ via the R packages *tidytext* (v0.4.3) ^68^ and *stm* (v1.3.8) ^69^, with the number of topics set to *k* = 3 to capture distinct, high-level biological themes with high semantic coherence (**Supplement**; **Fig. S10**; **Table S7A**). Each module was assigned to its most frequent topic across 1000 topic model fits with random initialization; the proportion of fits supporting each assignment is reported as assignment reproducibility (**Supplement**; **Table S7B**). We refer to each topic as a “[biological] program” (**Fig. 3B**).

#### Assignment of ASD genes to programs and functional subclustering

We next assigned genes to each unique program using a GO overlap scoring approach. First, a program-specific GO vocabulary was built by taking all significantly enriched Biological Process (BP) terms (FDR < 0.05) from the modules belonging to each program. For each ASD gene (FDR < 0.001), we then counted how many of its GO terms overlapped each program’s vocabulary, and assigned the gene to the program with the greatest overlap. A curated set of genes whose top-scoring program conflicted with their established primary biology was manually overridden to the better-fitting program, and ASD genes with no scorable GO overlap were manually assigned by known function. This yielded a single, mutually exclusive program label per gene (**Table S9A**).

We also explored subclusters within each primary program’s gene set, in this case using an unsupervised, exploratory approach, as the number and identity of the subgroups were not specified a priori (in contrast with the three primary programs, which were defined by the topic model). For each gene, we retrieved all BP GO terms, this time without filtering by term size to maximize the number of genes assigned to at least one pathway. We then constructed a gene x GO-term binary membership matrix and computed the pairwise cosine similarity between all genes. Genes were clustered by Ward’s D2 hierarchical clustering on the cosine distance (1 – similarity). Genes with no GO annotation (*n* = 3) were retained in the matrix as zero-vectors and placed by variance minimization. The resulting clusters were assigned biological labels post hoc and manually refined to improve functional coherence (**Figs. 2G, 4B**; **Table S9B-D**).

#### Developmental expression trajectories and pre- or post-natal bias

Finally, to visualize the developmental trajectory of the genes uniquely assigned to each biological program, we used bulk RNA-seq data from the BrainSpan Atlas of the Developing Human Brain ^23^, restricted to the 11 neocortical regions and processed following ^7^. Briefly, genes were retained if RPKM >= 0.5 in >= 80% of samples from at least one neocortical region x epoch (prenatal/postnatal) combination, and expression was normalized using log_2_(RPKM + 1) and *Z*-scored per gene across all samples. For each sample, we computed the mean *Z*-score of ASD genes assigned to each program (as described above). To center trajectories relative to background expression, we first subtracted the genome-wide per-sample mean, then *Z*-scored the resulting values within each post-conception week (pcw) across all samples and programs, such that y = 0 represents the average expression level across programs at that timepoint. Developmental trajectories were smoothed using LOESS regression across the 11 developmental periods defined by ^23^, spanning early fetal development through middle adulthood, with full-term birth defined as 40 pcw (**Fig. 3D**).

From the same BrainSpan data, we also quantified each gene’s prenatal versus postnatal expression bias (**Fig. 3E**). Samples were first collapsed to a single donor-level mean per gene (averaging across regions to avoid collinearity between donor identity and prenatal status), and for each gene we fit a linear model of expression on prenatal status with sex as a covariate, taking the sign-flipped *t*-statistic of the prenatal term as the bias score (negative = higher prenatal, positive = higher postnatal). For each program, this distribution of bias scores was then compared against the background of non-ASD genes using a two-sided Wilcoxon rank-sum test (Benjamini-Hochberg corrected), with Cliff’s delta as the effect size (bounded [–1, 1]).

Together, these analyses defined the three functional programs (**Fig. 3B**), their developmental trajectories and prenatal bias (**Fig. 3D-E**), and their functional subclusters (**Fig. 5B**); topic model outputs and gene-to-program assignments are provided in the **Supplement** (**Fig. S10**; **Tables S7A, S8A-B, S9A-D**).

### Single-cell gene set enrichment scoring (AUCell)

Gene set enrichment at the single-cell level was quantified using AUCell (v1.26.0) ^40^. Input expression counts were derived from Seurat’s SCT assay, in which raw counts were normalized using SCTransform to correct for sequencing depth differences. For each cell, genes were ranked by expression level, and the area under the recovery curve (AUC) was calculated for each gene set using the top 5% of ranked genes. The AUC reflects the proportion of gene set members falling within this top-ranked fraction and their relative expression compared to all other genes in the same cell, providing a cell-level gene set enrichment score that is robust to differences in library size and normalization. Cells with AUC scores exceeding Global_k1 were classified as high-scoring, where Global_k1 is defined as the quantile corresponding to probability 1 − (*t*ℎ*rP*/*N* + 0.25) of a normal distribution fitted to the global AUC score distribution, with default parameters (*t*ℎ*rP* = 0.01, *N* = number of cells) and with mean and standard deviation estimated across all cells. For default parameters, this approximates the 75th percentile of the fitted normal distribution. The proportion of high-scoring cells per annotated cell type was computed to assess cell type-level enrichment across modules and SynGO HotNets.

We further investigated the association between gene set program activity and ASD relevance as quantified by scDRS. For each lineage cluster with significant enrichment found in **Fig. 1D** (FDR < 0.05), the Spearman correlation between cell-level AUCs and normalized scDRSs was used as the test statistic *r_condition*. Significance was assessed using a two-sided MC test. The same Spearman correlation was computed between each of the 1000 matched normalized control scores and AUCs within the same set of cells, generating an empirical null distribution of control test statistics. The MC *P* value was computed as (1 + number of control correlations whose absolute value equals or exceeds |*r_condition*|) / (1 + 1000), where the use of absolute values implements a two-sided test. The MC test does not require the assumption of cell independence, which is particularly important in single-cell analyses for which cells within the same cluster are likely to be correlated. Multiple testing correction was applied across all gene set and lineage cluster combinations. These analyses produced the AUCell results shown in **Figs. 2C, 3C, S11, S22-S23,** and **Table S7B.**

### Synaptic gene ontology enrichment analysis (SynGO)

To characterize the synaptic functions disrupted by rare genetic variation in ASD, we mapped *de novo* mutation (DNM) counts onto the SynGO gene ontology framework ^25^, which assigns 1602 unique genes to biological processes (BP; 1126 genes) and cellular components (CC; 1487 genes), organized as nested tree structures of synaptic subdomains (hereafter referred to as “compartments”). Compartment assignments are curated by domain experts from published experimental evidence for the localization and function of synaptic proteins. After mapping to our cohort data, we identified overlap for 1533 (all SynGO), 1076 (BP), and 1425 (CC) genes. Because DNM rates in ASD probands exceed those in siblings even in genes below the threshold of individual TADA test significance, we included all SynGO genes in compartment-level analyses to capture the full distribution of synaptic vulnerability (**Supplement**). Analyses were restricted to compartments with >= 10 genes. Damaging DNMs were defined as PTVs, Mis2, and Mis1. Mis1 variants were included as they carry information at the gene set level, though they are less informative at the single-gene level. For each SynGO compartment, we quantified DNM burden using *Z_adj_*, a gene-set-size-adjusted statistic that enables identification of compartments with unusually strong signal relative to their gene count. Briefly, *Z_adj_* is defined as the residual from a regression of *Z_all_*—a binomial test statistic quantifying total per-compartment mutation counts in probands relative to siblings (**Supplement**; **Equation 2**)—on gene count per compartment. We corrected for compartment size as gene count per compartment explained 61.5% of variance in *Z_all_* for CC and 41.9% for BP. We additionally quantified compartment-level DNM enrichment as the rate ratio of damaging DNMs in ASD probands (*n* = 38,680) relative to unaffected siblings (*n* = 9,567), defined as the per-subject mutation rate in probands divided by the per-subject mutation rate in siblings (**Equation 3**).

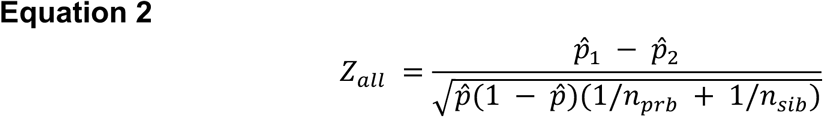

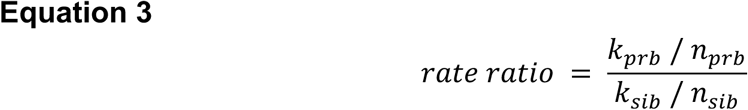

To identify subnetworks of compartments that were highly associated with ASD, we applied Hierarchical HotNet ^41^ to *Z_adj_* values within the SynGO tree structure. This method assigns scores to network edges based on the *Z_adj_* values of connected compartments and their topological proximity, then removes low-scoring edges to identify densely-connected subnetworks of compartments with elevated signal per gene.

Because many SynGO compartments share genes, we further partitioned genes into non-overlapping sets defined by the major child nodes of the root compartment (BP or CC), and computed the rate ratio separately for each independent gene set to distinguish contributions of distinct synaptic subdomains. We additionally computed per-gene mutation rates in probands by dividing the total number of *de novo* mutations observed in each gene set by the number of genes in that set, and contrasted these rates across independent gene sets under a binomial assumption.

These analyses produced the compartment-level DNM enrichment and HotNet subnetworks in **Figs. 3A-C,4A** and **Table S13A-B** with the independent-gene-set rate ratios, LOEUF constraint, and supporting analyses in the **Supplement** (**Figs. S18-23**).

### Enrichment of ASD-associated genes for developmental corticostriatal axonal proteome

The axonal proteomics data set ^42^ consisted of corticostriatal axonal proteomes quantified across four developmental stages in mice: neonate, early postnatal, preweaning, and adult. The original data set consisted of measurements on 2274 proteins, the developmental trajectory cluster to which they were assigned, and the mouse gene symbol when available. Because 9 proteins had no gene symbol, the remaining 2265 were processed as follows: retrieve mouse Ensembl gene id using *biomaRt* ^70^, which were available for 2213 of the proteins; retrieve a human Ensembl gene id matching the mouse Ensembl gene id, of which 2169 could be matched and gene names assigned; and, for the remaining 96 unmatched proteins, 43 could be mapped onto human genes by hand curation and *biomaRt*, whereas 53 were removed from further analyses. For our analysis, human Ensembl gene ids were available for 2212 of the 2265 mouse proteins. We then matched 2128 of these genes to the rare variant association results in ^8^.

In the axonal proteomics ^42^, proteins were assigned to one of six developmental clusters. However, clusters 1 and 5 showed very similar patterns in their rising developmental trajectory. Thus, these trajectories were combined to a set consisting of 634 genes. Of these, 43 had FDR < 0.001 in the ASC data. This combined set of proteins/genes is highly enriched for these ASD genes (OR = 2.342; 95% CI: 1.488-3.682; *P* = 0.0001) relative to the remaining set of 1,494 proteins/genes. Next, PANTHER (https://pantherdb.org/ ^71^) was used to obtain gene set enrichment for these 43 genes relative to the universe of 2128 genes by using Fisher exact test and FDR correction for multiple testing.

### Enrichment of ASD-associated modules for synaptic gene sets

We tested each of the ASD-associated modules for over-representation of synaptic genes, both overall (using the SynGO synapse cellular component root term [GO:0045202; 1,487 genes]) and within individual SynGO HotNet neighborhoods. All enrichments used Fisher’s exact test with two-sided *P* values, BH-corrected across modules within each synaptic gene set. Background was defined as brain-expressed genes ^72^ within the module’s study network for the overall test, and as SynGO-annotated genes within that network for the HotNet tests.

### Developmental disorder/intellectual disability comorbidity stratification of cell-type enrichment and gene set burden

Finally, we examined how cell-type enrichments differed between ASD probands with recorded DD/ID (ASD_DDID_, hereafter referred to as ASD_D_) and those without (ASD_DDID-Absent_, now ASD_DA_). Unlike the SynGO analyses, here we included *de novo* variant classes PTV, Mis2, and DEL, both to align with prior analyses showing the clearest proband-over-sibling enrichment in the ASC cohort ^8^, and because Mis1 variants are less reliable for single-gene-level analyses compared to gene-set level. For each of the 253 high-confidence ASD genes, we first computed the per-proband mutation rate difference 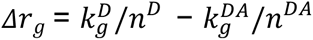, where 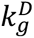 and 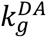 denote the number of damaging DNMs in ASD_D_ and ASD_DA_ probands, respectively, and *n^D^* = 13,841 and *n^DA^* = 24,839 the corresponding proband sample size. Gene with *Δr_g_* < 0 (indicating a higher per-proband mutation rate in ASD_DA_ than ASD_D_ probands) were designated ASD_DA_ genes, the remainder were designated ASD_D_ genes. ASD_D_ genes were then subdivided by their mutation fraction 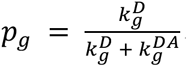 the proportion of damaging DNMs attributable to ASD_D_ probands (**Fig. 6A**). Genes with *p*_g_ exceeding the cohort-wide background expectation *p*_0_ = 64.37% (computed across all 253 genes) were defined as having a high level of DD/ID-associated mutation burden (ASD_DDID-High_, now ASD_DH_); genes below the background expectation were considered to have a low level of DD/ID-associated mutation burden (ASD_DDID-Low_, now ASD_DL_) (**Fig. 6B**).

To balance power with resolution, scDRS enrichment was assessed separately for ASD_DH_ genes, the combined ASD_DL_ and ASD_DA_ set for power, and ASD_DA_ genes for finer resolution. Within each lineage cluster with significant enrichment in **Fig. 1D** (FDR < 0.05), enrichment was evaluated using the same group-level scDRS framework described above (**Fig. 6C**). To characterize the developmental dynamics, we performed scDRS group-level analysis on cells stratified by both lineage cluster and developmental stage, treating each cluster-stage combination as an independent group (**Fig. 6D)**. Groups with fewer than 150 cells were excluded, and enrichment significance was assessed as described in the Methods (scDRS framework) section. These analyses underlie the DD/ID comorbidity stratification cell-type enrichment results shown in **Figs. 6**, **S26**, and **Table S14.**

We additionally quantified how gene set (modules, SynGO, and spatial) enrichment differed between probands with and without recorded DD/ID by computing observed-to-expected ratios ^73^ for damaging DNMs across all ASD probands, ASD_D_ probands, ASD_DA_ probands, and unaffected siblings as a negative control. Spatial gene sets (germinal zone, cortical plate, thalamus) were derived from a prenatal spatial transcriptomic factor analysis ^18^. Expected DMN counts were derived from per-gene, per-variant-class mutation rates based on gnomAD v4.1. Pairwise differences between gene sets were assessed using a two-sided conditional binomial test to estimate relative burden of gene sets ^73^. Full details are provided in the **Supplement**. These analyses underlie the DD/ID comorbidity stratification gene set enrichment results shown in **Figs. S27-S29,** and **Tables S15-S17.**

All analyses were performed in R version 4.4.0. All *P* values were corrected for multiple testing using the Benjamini–Hochberg procedure ^74^.

## References

1. Bai, D. et al. Association of genetic and environmental factors with autism in a 5-country cohort. JAMA Psychiatry 76, 1035–1043 (2019).

2. Fu, J. M. et al. Rare coding variation provides insight into the genetic architecture and phenotypic context of autism. Nat. Genet. 54, 1320–1331 (2022).

3. Grove, J. et al. Identification of common genetic risk variants for autism spectrum disorder. Nat. Genet. 51, 431–444 (2019).

4. Margolis, M. P. et al. From variants to mechanisms: Neurogenomics in the post-GWAS era. Neuron 113, 3509–3529 (2025).

5. De Rubeis, S. et al. Synaptic, transcriptional and chromatin genes disrupted in autism. Nature 515, 209–215 (2014).

6. Iossifov, I. et al. The contribution of de novo coding mutations to autism spectrum disorder. Nature 515, 216–221 (2014).

7. Satterstrom, F. K. et al. Large-scale exome sequencing study implicates both developmental and functional changes in the neurobiology of autism. Cell 180, 568–584.e23 (2020).

8. Satterstrom, F. K. et al. Rare variation illuminates the distinct and pleiotropic genetic architecture of autism across neuropsychiatric traits. medRxiv 2026.08.24.26360398 (2026) doi:10.64898/2026.08.24.26360398.

9. Willsey, H. R., Willsey, A. J., Wang, B. & State, M. W. Genomics, convergent neuroscience and progress in understanding autism spectrum disorder. Nat. Rev. Neurosci. 23, 323–341 (2022).

10. Parikshak, N. N. et al. Integrative functional genomic analyses implicate specific molecular pathways and circuits in autism. Cell 155, 1008–1021 (2013).

11. Willsey, A. J. et al. Coexpression networks implicate human midfetal deep cortical projection neurons in the pathogenesis of autism. Cell 155, 997–1007 (2013).

12. Gandal, M. J. et al. Broad transcriptomic dysregulation occurs across the cerebral cortex in ASD. Nature 611, 532–539 (2022).

13. Velmeshev, D. et al. Single-cell genomics identifies cell type-specific molecular changes in autism. Science 364, 685–689 (2019).

14. Wamsley, B. et al. Molecular cascades and cell type-specific signatures in ASD revealed by single-cell genomics. Science 384, eadh2602 (2024).

15. Bhaduri, A. et al. An atlas of cortical arealization identifies dynamic molecular signatures. Nature 598, 200–204 (2021).

16. Cadwell, C. R., Bhaduri, A., Mostajo-Radji, M. A., Keefe, M. G. & Nowakowski, T. J. Development and arealization of the cerebral cortex. Neuron 103, 980–1004 (2019).

17. Qian, X. et al. Spatial transcriptomics reveals human cortical layer and area specification. Nature 644, 153–163 (2025).

18. Aivazidis, A. et al. A spatial transcriptomic atlas of autism-associated genes identifies convergence in the developing human thalamus. bioRxivorg 2025.11.05.685843 (2025) doi:10.1101/2025.11.05.685843.

19. Ball, G. et al. Molecular signatures of cortical expansion in the human foetal brain. Nat. Commun. 15, 9685 (2024).

20. Wang, L. et al. Molecular and cellular dynamics of the developing human neocortex. Nature 1–10 (2025).

21. Sanders, S. J. et al. Insights into autism spectrum disorder genomic architecture and biology from 71 risk loci. Neuron 87, 1215–1233 (2015).

22. Gandal, M. J. et al. Transcriptome-wide isoform-level dysregulation in ASD, schizophrenia, and bipolar disorder. Science 362, eaat8127 (2018).

23. Li, M. et al. Integrative functional genomic analysis of human brain development and neuropsychiatric risks. Science 362, eaat7615 (2018).

24. Wen, C. et al. Cross-ancestry atlas of gene, isoform, and splicing regulation in the developing human brain. Science 384, eadh0829 (2024).

25. Koopmans, F. et al. SynGO: An evidence-based, expert-curated knowledge base for the synapse. Neuron 103, 217–234.e4 (2019).

26. Litman, A. et al. Decomposition of phenotypic heterogeneity in autism reveals underlying genetic programs. Nat. Genet. 57, 1611–1619 (2025).

27. Greig, L. C., Woodworth, M. B., Galazo, M. J., Padmanabhan, H. & Macklis, J. D. Molecular logic of neocortical projection neuron specification, development and diversity. Nat. Rev. Neurosci. 14, 755–769 (2013).

28. Harris, K. D. & Shepherd, G. M. G. The neocortical circuit: themes and variations. Nat. Neurosci. 18, 170–181 (2015).

29. Tasic, B. et al. Shared and distinct transcriptomic cell types across neocortical areas. Nature 563, 72–78 (2018).

30. Hawrylycz, M. J. et al. An anatomically comprehensive atlas of the adult human brain transcriptome. Nature 489, 391–399 (2012).

31. Matoba, N. et al. Common genetic risk variants identified in the SPARK cohort support DDHD2 as a candidate risk gene for autism. Transl. Psychiatry 10, 265 (2020).

32. Bakken, T. E. et al. Comparative cellular analysis of motor cortex in human, marmoset and mouse. Nature 598, 111–119 (2021).

33. Hodge, R. D. et al. Conserved cell types with divergent features in human versus mouse cortex. Nature 573, 61–68 (2019).

34. Allen Institute for Brain Science. MapMyCells [Software]. RRID: SCR_024672. https://knowledge.brain-map.org/mapmycells/process?refTaxonomyId=10xGene (2025).

35. Wu, S. J. et al. Cortical somatostatin interneuron subtypes form cell-type-specific circuits. Neuron 111, 2675–2692.e9 (2023).

36. Tsyporin, J. et al. Competing programs shape cortical sensorimotor-association axis development. Nature 1–12 (2026).

37. Desikan, R. S. et al. An automated labeling system for subdividing the human cerebral cortex on MRI scans into gyral based regions of interest. Neuroimage 31, 968–980 (2006).

38. Mohajeri, K. et al. Transcriptional and functional consequences of alterations to MEF2C and its topological organization in neuronal models. Am. J. Hum. Genet. 109, 2049–2067 (2022).

39. Tsai, Y.-C., et al. Morphogen-guided neocortical organoids recapitulate regional areal identity and model neurodevelopmental disorder pathology. bioRxivorg (2025) doi:10.1101/2025.09.02.672952.

40. Aibar, S. et al. SCENIC: single-cell regulatory network inference and clustering. Nat. Methods 14, 1083–1086 (2017).

41. Reyna, M. A., Leiserson, M. D. M. & Raphael, B. J. Hierarchical HotNet: identifying hierarchies of altered subnetworks. Bioinformatics 34, i972–i980 (2018).

42. Dumrongprechachan, V., Salisbury, R. B., Butler, L., MacDonald, M. L. & Kozorovitskiy, Y. Dynamic proteomic and phosphoproteomic atlas of corticostriatal axons in neurodevelopment. Elife 11, (2022).

43. Kaplanis, J. et al. Evidence for 28 genetic disorders discovered by combining healthcare and research data. Nature 586, 757–762 (2020).

44. Li, Y. et al. Splicing-dependent trans-synaptic SALM3-LAR-RPTP interactions regulate excitatory synapse development and locomotion. Cell Rep. 12, 1618–1630 (2015).

45. Bourgeron, T. From the genetic architecture to synaptic plasticity in autism spectrum disorder. Nat. Rev. Neurosci. 16, 551–563 (2015).

46. de la Torre-Ubieta, L., Won, H., Stein, J. L. & Geschwind, D. H. Advancing the understanding of autism disease mechanisms through genetics. Nat. Med. 22, 345–361 (2016).

47. Zoghbi, H. Y. Postnatal neurodevelopmental disorders: meeting at the synapse? Science 302, 826– 830 (2003).

48. Rakic, P. Specification of cerebral cortical areas. Science 241, 170–176 (1988).

49. Jorstad, N. L. et al. Transcriptomic cytoarchitecture reveals principles of human neocortex organization. Science 382, eadf6812 (2023).

50. O’Leary, D. D. Do cortical areas emerge from a protocortex? Trends Neurosci. 12, 400–406 (1989).

51. He, X. et al. Integrated model of de novo and inherited genetic variants yields greater power to identify risk genes. PLoS Genet. 9, e1003671 (2013).

52. Zhang, M. J. et al. Polygenic enrichment distinguishes disease associations of individual cells in single-cell RNA-seq data. Nat. Genet. 54, 1572–1580 (2022).

53. de Leeuw, C. A., Mooij, J. M., Heskes, T. & Posthuma, D. MAGMA: generalized gene-set analysis of GWAS data. PLoS Comput. Biol. 11, e1004219 (2015).

54. Hao, Y. et al. Dictionary learning for integrative, multimodal and scalable single-cell analysis. Nat. Biotechnol. 42, 293–304 (2024).

55. Scrucca, L., Fraley, C., Murphy, T. B. & Raftery, A. E. Model-Based Clustering, Classification, and Density Estimation Using Mclust in R. (Chapman & Hall/CRC, Philadelphia, PA, 2023).

56. Street, K. et al. Slingshot: cell lineage and pseudotime inference for single-cell transcriptomics. BMC Genomics 19, (2018).

57. Markello, R. D. et al. Standardizing workflows in imaging transcriptomics with the abagen toolbox. Elife 10, e72129 (2021).

58. Arnatkeviĉiūtė, A., Fulcher, B. D. & Fornito, A. A practical guide to linking brain-wide gene expression and neuroimaging data. NeuroImage 189, 353–367 (2019).

59. Fischl, B., Sereno, M. I., Tootell, R. B. & Dale, A. M. High-resolution intersubject averaging and a coordinate system for the cortical surface. Hum. Brain Mapp. 8, 272–284 (1999).

60. Fulcher, B. D., Arnatkeviciute, A. & Fornito, A. Overcoming false-positive gene-category enrichment in the analysis of spatially resolved transcriptomic brain atlas data. Nat. Commun. 12, 2669 (2021).

61. Cornblath, E. J. et al. Temporal sequences of brain activity at rest are constrained by white matter structure and modulated by cognitive demands. *Commun*. Biol. 3, 261 (2020).

62. Markello, R. D. & Misic, B. Comparing spatial null models for brain maps. Neuroimage 236, 118052 (2021).

63. Markello, R. D. et al. Neuromaps: Structural and functional interpretation of brain maps. Nat. Methods 19, 1472–1479 (2022).

64. Dear, R. et al. Cortical gene expression architecture links healthy neurodevelopment to the imaging, transcriptomics and genetics of autism and schizophrenia. Nat. Neurosci. 27, 1075–1086 (2024).

65. Langfelder, P. & Horvath, S. Eigengene networks for studying the relationships between co-expression modules. BMC Syst. Biol. 1, 54 (2007).

66. Ashburner, M. et al. Gene ontology: tool for the unification of biology. The Gene Ontology Consortium. Nat. Genet. 25, 25–29 (2000).

67. Liu, L., Tang, L., Dong, W., Yao, S. & Zhou, W. An overview of topic modeling and its current applications in bioinformatics. Springerplus 5, 1608 (2016).

68. Silge, J. & Robinson, D. Tidytext: Text mining and analysis using tidy data principles in R. J. Open Source Softw. 1, 37 (2016).

69. Roberts, M. E., Stewart, B. M. & Airoldi, E. M. A model of text for experimentation in the social sciences. J. Am. Stat. Assoc. 111, 988–1003 (2016).

70. Durinck, S., Spellman, P. T., Birney, E. & Huber, W. Mapping identifiers for the integration of genomic datasets with the R/Bioconductor package biomaRt. Nat. Protoc. 4, 1184–1191 (2009).

71. Mi, H. et al. Protocol Update for large-scale genome and gene function analysis with the PANTHER classification system (v.14.0). Nat. Protoc. 14, 703–721 (2019).

72. Uhlén, M. et al. Proteomics. Tissue-based map of the human proteome. Science 347, 1260419 (2015).

73. Samocha, K. E. et al. A framework for the interpretation of de novo mutation in human disease. Nat.Genet. 46, 944–950 (2014).

74. Benjamini, Y. & Hochberg, Y. Controlling the false discovery rate: A practical and powerful approach to multiple testing. J. R. Stat. Soc. Series B Stat. Methodol. 57, 289–300 (1995).

