## Supplemental text and figures for "Autism genes converge on three functional programs organized by neuronal subclass, developmental timing, and cortical patterning"

**This PDF file includes**

Supporting Methods and Results
Figures S1 to S30
Legends for Tables S1 to S18
SI References

**Other supporting materials for this manuscript include the following**

Tables S1 to S18

### Cell type enrichment

#### *MapMyCell*

To characterize the anatomical distribution of MGE-SST subtypes, we mapped upper-layer and deep-layer MGE-SST cells to the 10x Whole Human Brain Atlas using MapMyCell [1]. Major clusters representing at least 2% of the total cell population are shown, and the five clusters most strongly enriched for each subtype are highlighted on UMAP with cluster annotations summarized.

#### *Differential expression analysis of MGE-SST subtypes*

Differential expression between MGE-SST upper-layer and deep-layer subtypes was performed using pseudobulk aggregation followed by a generalized linear mixed model implemented in *glmmSeq* package (v0.5.7), with subtype as the fixed effect of interest, log2-transformed age as a covariate, and donor as a random effect (~ subtype + log2_age + (1 | dataset)). Raw counts were aggregated across cells by subtype and donor using *aggregateAcrossCells* from *scuttle* package (v1.14.0), and pseudobulk samples with fewer than 50 cells were discarded. Lowly expressed genes were removed using *filterByExpr* from *edgeR* package (v4.2.2). Library size normalization was performed using TMM scaling factors computed with *calcNormFactors*, and tagwise dispersions were estimated using *estimateDisp*. A likelihood ratio test was performed by comparing against a reduced model excluding the subtype term (~ log2_age + (1 | dataset)). *P* values from the likelihood ratio test were adjusted using the Benjamini-Hochberg procedure. Log2 fold changes (log2FC) were derived from model-predicted values for the deep-layer subtype coefficient. Genes with |log2FC| > 1 and FDR < 0.05 were considered differentially expressed.

#### *Characterization of MGE-SST subtype features*

MapMyCell revealed subtype-specific neuropeptide signatures within MGE-SST lineage (**Fig. S3**; **Table S2**). Natriuretic peptide C (NPPC) and proenkephalin (PENK) were exclusively enriched in the upper-layer subtype. Within the deep-layer subtype, the majority of clusters maintained a GABAergic profile. However, two clusters (MGE_252 and MGE_254) exhibited a dual-transmitter phenotype, co-expressing GABA and VGLUT3 (SLC17A8). Differential gene expression analysis further revealed that *NPY, TACR1, ANOS1, SYTL5, KIRREL3, PENK,* and *ADRA1A* were significantly upregulated in the ASD-enriched upper-layer SST subtype (**Fig. S4**; **Table S3**).

### Identification of ASD-associated modules and biological programs

#### *Module identification using logistic regression*

Because Transmission and De Novo Association (TADA) scores are driven partly by genes associated specifically with comorbid developmental delay/intellectual disability (DD/ID), we ran a sensitivity analysis for ASD-associated module identification using logistic regression with a binary outcome. The analytic framework was identical to the main analysis (employing the same stepwise conditional forward selection procedure), but the continuous *Z*-score-transformed TADA FDR outcome was replaced with a binary indicator of ASD gene status, defined at four TADA FDR thresholds: FDR < 0.001, FDR < 0.01, FDR < 0.05, and FDR < 0.1. Because the binary outcome does not have as much information as the continuous *Z*-score, this analysis is expected to have lower statistical power than the main linear regression approach and is a qualitative consistency check rather than a direct replication.

The logistic regression sensitivity analysis identified 10, 12, 14, and 18 independently associated modules at FDR thresholds of < 0.001, < 0.01, < 0.05, and < 0.1, respectively, representing a subset of the 28 modules identified by the main linear regression analysis. Core modules including *CUX1*$^{(+)}$, *NR3C1*$^{(+)}$, *SOX11*$^{(+)}$, and *MEF2C*$^{(+)}$ were independently significant across thresholds, and odds ratios remained stable as the threshold was relaxed (**Fig. S9**). However, many of the modules identified at one TADA FDR threshold were not significant at others, demonstrating the advantage of the linear regression model reported in the main text.

***Topic modeling methods***

Topic modeling is an unsupervised method that decomposes a collection of "documents" into a set of latent "topics," where each topic is a probability distribution over "words" and each document is represented as a mixture of topics. In our application, each module's set of enriched GO terms was treated as a document and the individual GO terms as words, such that topics correspond to recurring biological themes shared across modules. The structural topic model estimates two key quantities: the per-topic word distribution ($\beta$), which defines each topic by the GO terms most characteristic of it, and the per-document topic distribution ($\gamma$), which gives the proportion of each module attributable to each topic.

Models were fit with spectral initialization [2], the `stm` package default [3]. Briefly, spectral initialization identifies “anchor” words that occur almost exclusively within a topic from the matrix of word co-occurrences, and uses them to derive a starting estimate of the topic-word distributions before variational expectation-maximization. This fit is deterministic; reported word-topic probabilities (beta; **Fig. 2B**) are derived from this spectral fit.

The number of topics ($k$) was selected based on the trade-off between semantic coherence and exclusivity in the spectral fit model (**Fig. S10**; **Table S7A**). Semantic coherence measures how often the highest-probability words in a topic co-occur within the same document (here, GO term annotations per module), where higher values indicate more internally consistent topics. Exclusivity measures the degree to which the highest-probability words in a topic are unique to that topic rather than shared across topics; higher values indicate more distinct, non-overlapping topics. Models with few topics tend to have high coherence but low exclusivity (topics are broad and overlapping), while models with many topics tend to have high exclusivity but lower coherence (topics become too narrow). $k = 3$ maximizes semantic coherence among $k = 2$–$10$, while exclusivity increases monotonically with $k$ and so never identifies an interior optimum (**Fig. S10**). The final choice was based on the interpretability of the topics: $k = 3$ captures three biologically interpretable themes—chromatin regulation, neuronal morphogenesis, and synaptic transmission—without over-splitting the GO term space.

ASD-associated modules were then assigned to topics by consensus across repeated model fits. Given that spectral fit is deterministic, it does not provide estimates of how stable a module-topic assignment is. We therefore refit the $k = 3$ model 1000 times with random initialization; each module was assigned to its most frequent topic across restarts (**Fig. 2B**).

**Spatial enrichment patterns of ASD gene sets**

Unless stated otherwise, regional enrichment, the anterior-posterior (A-P) axis, and spin-based significance were computed as in the main **Methods** (spin test: parcel-wise Cornblath rotations at 41k fsaverage density, Spearman correlation). Occipital regions include the four occipital Desikan-Killiany parcels (pericalcarine, lingual, cuneus, lateral occipital). The "occipital enrichment score" of a gene set is each gene's regional expression *Z*-score averaged over these four regions, then averaged across the set.

***Robustness of the overall ASD adult cortical enrichment gradient***

<u>Occipital enrichment exceeds a gene property-matched null.</u> Rare variants tend to impact genes that are longer and more constrained [4,5]. Because gene length and expression level covary with regional transcriptomic profiles [6,7], and because gene-property structure is a known source of false-positive enrichment in spatial transcriptomic analyses [8], we compared the observed occipital enrichment to a null matched on these properties. Background genes were binned into tertiles on each of coding-sequence length, constraint (loss-of-function observed/expected upper bound fraction score; LOEUF [9]), and $\log_{10}$(mean expression) and crossed up to 27 strata. 5000 null sets were drawn to match the ASD set's per-stratum composition; significance was calculated as the one-sided empirical *P* (#[null ≥ observed] + 1)/(N + 1). Occipital enrichment exceeded both the unmatched null ($P = 2\times10^{-4}$) and the property-matched null ($P = 0.011$; **Fig. S7B**), indicating an effect beyond gene length, constraint, and expression level.

<u>The effect is distributed across many genes.</u> To test whether occipital enrichment is driven by a few genes that are highly expressed in occipital regions, we ranked ASD genes by their per-gene occipital score, removed the top *k* (*k* = 0:30), and recomputed the occipital enrichment of the remaining genes. Enrichment remained well above chance (the mean occipital score of the background) throughout, indicating a distributed rather than few-gene effect (**Fig. S7C**).

<u>The gradient is not confined to occipital regions.</u> To test whether the A-P gradient is driven solely by the occipital pole, we removed the four occipital regions and recomputed the A-P spin test on the remaining 30 cortical regions (1000 rotations). The gradient was essentially unchanged (full $\rho = 0.74$, $P_{spin} = 0.027$; occipital-removed $\rho = 0.65$, $P_{spin} = 0.037$; **Fig. S7D**), indicating a cortex-wide posterior bias rather than an occipital-only effect.

***Robustness of biological program-specific gradients***

<u>Relationship to dominant expression gradients.</u> To test whether the functional program ASD gene set spatial enrichment patterns aligned with the dominant axes of cortical gene expression, we correlated the ASD enrichment map with the three transcriptomic components of adult AHBA cortex from [10] (C1-C3), using a spin test (5000 spins). All three programs showed a positive A-P enrichment gradient, but only GR was significant when accounting for spatial autocorrelation ($P_{spin}$ = 0.005; **Fig. S14A**). Interestingly, SYN and MORPH were both significantly aligned with AHBA C3 ($P_{spin}$ = 0.044 and 0.016, respectively), which itself is enriched for synaptic programs and genes implicated in schizophrenia [10].

We additionally repeated all three analyses from the overall ASD gene set for the three GO-defined programs. GR maintained its A-P enrichment after occipital removal ($P_{spin}$ = 0.041, 1000 rotations), while SYN and MORPH became more aligned with A-P following occipital removal ($P_{spin}$ = 0.047 and 0.053, respectively; **Fig. S14B**). Furthermore, GR occipital enrichment was still highly significant after matching for gene properties (matched *P* = 0.001), whereas SYN was only nominal (*P* = 0.085) and MORPH was not enriched (*P* = 0.875) (unmatched $P <= 2\times10^{-4}$ for GR/SYN; MORPH unmatched *P* = 0.200; **Fig. S14C**). Concordantly, the drop top-*k* analysis showed GR occipital enrichment was distributed across many genes and SYN partially so, whereas MORPH fell to chance (**Fig. S14D**). Together these indicate that the robust, gene-property-independent posterior-occipital signal is carried specifically by the GR program; the SYN gradient is largely attributable to spatial autocorrelation and shared gene properties.

#### *Convergence of ASD gene spatial enrichment with pancortical transcriptomic dysregulation*

We asked whether regional ASD gene enrichment co-localizes with regional ASD transcriptomic dysregulation across the 11 cortical Brodmann regions of [11]. DK regions were mapped to the 11 Brodmann regions using the study's Allen-atlas region matching table, and enrichment was averaged across each Brodmann region. Per-region dysregulation was quantified as the number of genes differentially expressed between ASD and controls with ($|\log_2 FC| > 1$ and FDR < 0.05). We correlated ASD gene enrichment with dysregulation across the 11 regions using Spearman's *ρ*. Significance was assessed with a gene-set resampling null: 10,000 size-matched and brain-expressed gene sets were each mapped to the 11 regions and standardized per DK region across the null draws, then correlated with the dysregulation map; permutation *P* was one-sided.

ASD gene enrichment and transcriptomic dysregulation both peaked in the primary visual cortex (BA17) and were positively correlated in GR (*ρ* = 0.54) and overall ASD (*ρ* = 0.42; **Fig. S15A**). Against the resampling null, these correlations were directionally consistent but did not reach significance (one-sided permutation *P* = 0.15, 0.23, 0.35, and 0.43 for GR, ASD, SYN, and MORPH; **Fig. S15B**). Given only 11 regions, we interpret this as a directional convergence between rare-variant gene expression topography and transcriptomic pathology rather than an independent significant association.

#### *Gene-set spatial enrichment in the developing cortex*

To examine brain region-specific enrichment patterns in the developing human cortex, we obtained laminar fetal cortical expression data from the µBrain atlas [12]. This atlas provides laser microdissection microarray expression of 10,024 genes across 27 cortical regions and five transient tissue zones—ventricular (VZ), subventricular (SVZ), intermediate (IZ), subplate (SP), and cortical plate (CP)—sampled from the left hemisphere of four mid-gestation donors (15-21 post-conception week). Each gene's expression was *Z*-scored across all tissue samples (cortical region x layer x donor).

For each biological program (GR, SYN, and MORPH), we tested spatial enrichment relative to a background gene set using linear mixed-effects models. Genes were labeled as program members or background, and the program versus background difference in expression (i.e., program enrichment above background) was estimated as an estimated-marginal-means contrast using the R package emmeans v2.0.4 [13]. Regional enrichment within each layer was obtained by fitting scaled_expression ~ program_member[0,1] x region + (1 | gene) separately within each of the five zones and extracting the per-region program versus background contrast. Laminar (zone) enrichment was obtained separately by fitting scaled_expression ~ program_member[0,1] x zone + (1 | gene) + (1 | region:zone) across the regions sampled in all five zones and extracting the per-zone contrast. Contrasts were BH-corrected across regions within each layer or across zones. Each model was run using two background definitions: (1) the full measured µBrain transcriptome (all non-ASD genes) to test enrichment relative to all expressed genes, and (2) the ASD gene set (TADA FDR < 0.001; $n$ = 165 present in µBrain; GR $n$ = 63; MORPH $n$ = 55; SYN $n$ = 47) to test enrichment relative to ASD genes. Analysis results are reported in **Table S11A-B**.

Relative to all expressed genes, ASD genes across all three programs were broadly enriched in the cortical plate (**Fig. S16A**; **Table S11A**). Against the ASD gene background, however, GR and MORPH genes were significantly enriched across ventricular and subventricular zones (FDR < 0.05; **Fig. S16A**; **Table S11B**). In contrast, SYN remained enriched across the cortical plate, especially among hippocampal and paleocortical regions. Notably, only GR genes showed germinal zone enrichment in the posterior primary visual cortex, paralleling the adult gradient. These laminar patterns were consistent with each program's developmental trajectory (**Fig. 2D-E**).

To provide orthogonal evidence for the spatial associations identified in the µBrain analysis using an atlas that includes both cortical and subcortical regions, we examined spatial enriched gene sets derived from a prenatal spatial transcriptomic atlas [14], including germinal zone (GZ, $n$ = 26), cortical plate (CP, $n$ = 5), and thalamus (THL, $n$ = 44), that overlapped our 253 ASD genes. Representative ASD genes of different programs showed distinct regional expression patterns during mid-gestation (**Fig. S16B**), including GR gene *CREBBP*, which was enriched in GZ, SYN gene *STXBP1*, which showed preferential expression in THL, and MORPH gene *MEF2C*, which was enriched in CP.

Following Aivazidis et al., ASD genes were classified as exhibiting robust or non-robust spatial-factor expression according to their maximal observed expression across regions. Among genes with robust spatial-factor expression, GR genes were predominantly associated with GZ, SYN genes with THL, whereas MORPH genes were distributed across GZ, CP, and THL (**Fig. S16C**). Fisher's exact tests comparing GZ and THL showed that GR was significantly enriched in GZ (21/26, 81% vs. 15/44, 34%; OR = 7.85, 95% CI: 2.29–32.21; FDR = $2.85 \times 10^{-4}$), whereas SYN was significantly enriched in THL (0/26, 0% vs. 18/44, 41%; FDR = $1.47 \times 10^{-4}$). MORPH did not differ significantly between GZ and THL (5/26, 19% vs. 11/44, 25%; OR = 0.72, 95% CI: 0.17–2.65; $P$ = 0.77). Because only five ASD genes were assigned to CP, statistical comparisons involving this region were not performed owing to insufficient power.

**Localization of ASD DNM burden to SynGO synaptic compartments**

***Overall enrichment of synaptic genes compared to other brain-expressed genes***

To assess the overall enrichment of synaptic genes relative to other brain-expressed genes [15] (*n* = 15,348, downloaded from https://www.proteinatlas.org/search/NOT+tissue_category_rna%3Abrain%3Bnot+detected), we compared *de novo* mutation (DNM) counts in ASD probands vs unaffected siblings across SynGO-annotated genes and a mutation-rate-matched background. For each of 1000 iterations, we drew a random sample of brain-expressed non-SynGO genes equal in size to the SynGO gene set (*n* = 1533 intersecting with the brain-expressed background). To ensure the sampled background matched the mutation rate profile of SynGO genes, and thus that enrichment reflects differential DNM burden rather than mutations, we binned all brain-expressed genes into deciles based on background mutation rate distribution of SynGO genes, then sampled non-SynGO genes with probability proportional to those bin frequencies. For each iteration, we applied a Fisher's exact test to the 2x2 table of proband and sibling DNM counts in the SynGO genes versus the sampled background, and recorded the odds ratio ([proband synaptic / sibling synaptic] / [proband background / sibling background]). We report the mean odds ratio and 95% confidence interval across all 1000 iterations. The same approach was applied to transcription factor genes [16] and epigenetic regulator genes [17] for comparison. Transcription factors that were also annotated as epigenetic regulators were reassigned to the epigenetic regulator set, making the two sets mutually exclusive.

Across 1,000 random sampling experiments, synaptic genes were 1.32 times more likely to carry damaging DNMs than other brain-expressed genes ($P < 2.2\times10^{-16}$), confirming that the synapse is a preferential target of rare variants in ASD (**Fig. S30A**). For comparison, gene sets encoding transcription factors and epigenetic regulators showed somewhat higher enrichment—1.55-fold (1470 genes; $P < 2.2\times10^{-16}$) and 1.70-fold (645 genes; $P < 2.2\times10^{-16}$), respectively—consistent with the chromatin regulation signal identified in our module analyses.

We additionally ran the same analysis using ASD genes (TADA FDR < 0.001) assigned to each biological program as the gene sets. All three program-defined gene sets showed substantially higher enrichment than the broader functional categories above, consistent with the concentration of rare variant signal in high-confidence ASD genes. SYN-assigned genes showed 4.12-fold enrichment (*n* = 56; 95% CI [4.07, 4.18]; $P < 2.2\times10^{-16}$), MORPH-assigned genes showed 3.68-fold enrichment (*n* = 69; 95% CI [3.60, 3.75]; $P < 2.2\times10^{-16}$), and GR-assigned genes showed 4.21-fold enrichment (*n* = 128; 95% CI [4.14, 4.28]; $P < 2.2\times10^{-16}$), each relative to a mutation-rate-matched brain-expressed background (**Fig. S30B**).

***Justification for including all genes in each compartment, regardless of TADA significance***

To confirm that compartment-level analyses were not driven exclusively by individually significant ASD genes, we partitioned synaptic genes into three bins by TADA FDR (FDR < 0.001, 0.001-0.05, and 0.05-1.0). DNM rates exceeded chance expectation in all three bins for both "process in synapse" (BP) (rate ratio [RR] = 8.91, $Z_{all}$=14.30, $P = 2.1\times10^{-46}$; RR = 2.60, $Z_{all}$ = 5.48, $P = 4.2\times10^{-8}$; RR = 1.36, $Z_{all}$ = 3.92, $P = 9.0\times10^{-5}$, respectively) and "synapse" (CC) (RR = 8.87, $Z_{all}$ = 13.23, $P = 5.7\times10^{-40}$; RR = 2.25, $Z_{all}$ = 4.15, $P = 3.4\times10^{-5}$; rate ratio = 1.28, $Z_{all}$ = 2.89, $P$ = 3.8×10⁻

[3], respectively), confirming that *de novo* variation in synaptic genes contributes to ASD liability even below the threshold of individual gene-level significance.

***Compartment-level rate ratio and unadjusted DNM enrichment across BP and CC***

Across both BP and CC, rate ratio tended to diminish from child to parent nodes, reflecting the attenuation of strong signal in peripheral nodes as larger, more heterogeneous compartments inherit genes from multiple children (**Table S13A-B**). However, this pattern was not observed in BP for pathways involving presynaptic membrane potential and (post)synaptic organization, which maintained similarly elevated rate ratios across topological levels, suggesting these represent biological enrichment rather than artifacts of tree inheritance.

For CC, cytosol compartments on both sides of the synapse produced substantially greater signal than ribosomal compartments: presynaptic cytosol ($Z_{all}$ = 3.63, $P = 1.4\times10^{-4}$; RR = 4.54; 51 genes) vs postsynaptic cytosol ($Z_{all}$ = 5.70, $P = 7.8\times10^{-7}$; rate ratio = 15.01; 49 genes), with 25 genes shared between these compartments (**Table S13A**). For BP, peripheral nodes with the strongest signal included structural constituent of postsynaptic density ($Z_{all}$ = 5.70, $P = 5.9\times10^{-9}$; rate ratio = 10.71; 12 genes) and ligand-gated ion channel activity involved in regulation of presynaptic membrane potential (22 genes), while synaptic metabolism showed only modest enrichment ($Z_{all}$ = 1.43, $P$ = 0.076; rate ratio = 1.47; 103 genes) (**Table S13B**). Notably, ribosomal, metabolic, and translational compartments showed consistently low enrichment across both BP and CC, suggesting these synaptic functions are not predominant targets of rare variation in ASD.

***Selective constraint alone does not explain compartment-level enrichment***

Gene-level selective constraint against loss-of-function can be quantified by loss-of-function observed/expected upper bound fraction (LOEUF) score, the upper bound of the observed/expected ratio of predicted loss-of-function variants in human populations, where lower values indicate stronger constraint [9]. Finally, we evaluated mean LOEUF scores per compartment as a measure of evolutionary constraint, and assessed whether constraint levels were consistent with the patterns of DNM enrichment across compartments. Mean compartment LOEUF was inversely related to ASD enrichment (Spearman $\rho = -0.36$ for CC, $\rho = -0.27$ for BP; compartments with >=10 genes), and the subdomains with the strongest enrichment (presynaptic active zone and postsynaptic specialization for CC, synapse organization for BP) were among the most constrained. Several compartments with little or no enrichment were among the most tolerant, including synaptic vesicle (rate ratio = 2.31; $Z_{adj}$ = 0.17; 148 genes; mean LOEUF = 0.68) and neuronal dense core vesicle (rate ratio = 0.99; $Z_{adj} = -1.73$; 40 genes; mean LOEUF = 0.92; **Fig. S20**).

Constraint alone, however, did not predict ASD enrichment. Synapse maturation was highly constrained (mean LOEUF = 0.45; 34 genes) yet showed no excess of proband DNMs (rate ratio = 0.78; $Z_{adj} = -2.33$); postsynaptic cytoskeleton organization was similarly constrained and showed only modest enrichment (mean LOEUF = 0.38; 26 genes; rate ratio = 1.52; $Z_{adj} = -0.35$). In contrast, transmitter-gated ion channel activity regulating postsynaptic membrane potential was comparatively tolerant (mean LOEUF = 0.61; 38 genes) but was one of the most strongly enriched compartments (rate ratio = 4.23; $Z_{adj} = 3.02$). Constraint therefore contributes to, but does not fully account for, the distribution of ASD mutational burden across synaptic compartments.

***Distribution of per-gene rate enrichment across compartments ($R_P$)***

To assess whether compartment-level enrichment signals were concentrated in a small number of highly mutated genes or distributed across the compartment, we computed $R_P$ as the fraction of genes in each compartment for which the per-subject mutation rate in probands was higher than that in unaffected siblings. We evaluated deviations from the genome-wide expectation of $R_P = 0.45$ (meaning, 45% of all synaptic genes show an excess of mutations in probands) using 95% confidence intervals (CI; **Fig. S19**). An additional 41% (BP) to 43% (CC) of all synaptic genes had no mutations in any subject, meaning that most genes either show an excess of mutations in probands or are uninformative (very few show an excess in siblings).

For CC, compartments exceeding the upper 95% CI bound included presynaptic active zone and its cytoplasmic components and integral membrane components, as well as postsynaptic specialization and density (consistent with the HotNet subnetworks identified in the main text; **Fig. S19**). Compartments falling below expectation included presynaptic and postsynaptic ribosomal compartments, neuronal dense core vesicle, and presynaptic endosome.

For BP, compartments exceeding expectation included genes regulating presynaptic membrane potential and its ligand-gated ion channel activity, structural constituent of synapse, axo-dendritic transport, as well as genes involved in the modification of postsynaptic structure and actin cytoskeleton. Compartments falling below expectation were all related to synaptic translation, both pre- and postsynaptically, supporting the conclusion that translational machinery is not a primary target of rare variation in ASD. Notably, no genes were shared among the $R_P$ outlier compartments within either BP or CC (aside from parent-offspring dependencies), confirming that these are independent signals instead of the same set of genes driving enrichment across several compartments.

The full distribution of $Z_{all}$, $Z_{adj}$, rate ratio, and $R_P$ across all BP and CC compartments is provided in **Table S13A-B**.

***Genes with multiple synaptic functions carry the strongest ASD enrichment***

Postsynaptic specialization contained the highest number of ASD genes ($n$ = 30), but as the HotNet with the highest number of total genes ($n$ = 416), this represents the lowest fraction of total (7%; **Fig. 3B**). Presynaptic ion channels and active zone harbored the highest percent of ASD genes ($n$ = 10 of 48 total [21%] and $n$ = 16 of 109 total [15%]), respectively. Interestingly, we observed many shared ASD genes across HotNets. For example, *GRIN2B* was found in presynaptic active zone, presynaptic ion channels, presynaptic membrane, and postsynaptic organization, while *GRIN2A* was in the same HotNets except for presynaptic active zone. Similarly, other high-confidence ASD genes, including *SCN8A*, *SCN2A*, *GRIN1*, and *GRIA2*, among others, were also found in multiple HotNets.

To quantitatively address this observation of gene sharing across compartments, we partitioned SynGO genes by the major child nodes of the root compartment and computed the rate ratio for each non-overlapping set (**Methods**). For CC, rate ratio estimates were similar across partitions (rate ratio ranging from 2.36 to 2.55), indicating that pre- and postsynaptic structural genes are roughly equal targets of ASD rare variant burden (**Supplement**). In contrast, partitioning in BP revealed greater differences. By synaptic location, rate ratio was intermediate for genes exclusively presynaptic (rate ratio [95% CI] = 2.92 [2.07, 4.12]), lower for those exclusively postsynaptic (1.40 [0.93, 2.09]), and higher for those shared across both compartments (4.89 [2.50, 9.56]; **Fig. S21**). Genes found solely in signaling or organizational compartments showed modest enrichment (rate ratio = 1.89 [1.14, 3.14] and 1.86 [1.39, 2.49], respectively), whereas the largest rate ratio arose from the 45 genes shared between synaptic signaling and organization (rate ratio = 5.91 [2.91, 12.0]). These findings indicate that genes with multiple synaptic functions tend to be the most prominent targets of ASD rare variant burden, though this is not necessarily true of all such genes (**Fig. S21**).

***Presynaptic and postsynaptic HotNets implicate distinct cellular contexts***

To characterize cell type enrichment for genes in each HotNet subnetwork, we quantified cell-level gene set enrichment using the same AUCell-based framework. As expected, HotNet AUC scores were elevated in mature neurons relative to immature neurons and non-neuronal cells, increasing progressively along the pseudotime trajectory (**Fig. S22A-C**). Among mature neurons, EN-IT neurons showed stronger enrichment than non-IT neurons, particularly in L2/3-, L4-, and L5-IT neurons, with mature IN subtypes also showing enrichment (**Fig. 3C**). Postsynaptic organization exhibited the weakest and least cell-type-specific enrichment pattern across all HotNets, indicating broader and more uniform expression across cell types.

We then asked whether HotNet program activity within individual cells is associated with ASD genetic vulnerability by assessing the within-cluster correlation between HotNet AUC scores and scDRS in lineage clusters showing significant ASD relevance. Both presynaptic HotNets (active zone and membrane) correlated most strongly with scDRS in L4-IT neurons (two-sided Monte Carlo test, FDR $< 0.05$), indicating that these IT neurons with higher presynaptic program activity are associated with greater ASD vulnerability (**Fig. S23A**). Within L4-IT neurons, these associations strengthened postnatally. Both were non-significant in the third trimester, and the presynaptic active zone association became significant from infancy through adolescence, and the presynaptic membrane association became significant during adolescence. (**Fig. S23B**). In contrast, postsynaptic organization correlated with scDRS in IPC-EN and EN-Newborn, with similar magnitude correlations across both immature and mature INs. Within IPC-EN and EN-Newborn, these associations strengthened prenatally (first and second trimester) and became non-significant by the third trimester (**Fig. S23B**).

Together, these analyses reveal compartment-specific patterns of ASD vulnerability, with presynaptic Hotnet programs tracking ASD burden in postnatal mature IT neurons.

***DNM enrichment of CC and BP partitions***

Beyond the variation in the number of genes per compartment and the obvious parent-child dependence, compartments in disparate portions of the tree share genes. To address this issue, we compute rate ratios for the major child compartments of CC and BP after partitioning genes according to whether they occur in both compartments or whether they are unique to one. For CC, these partitions make little difference for the rate ratios, which range from 2.36 to 2.55. Compartments assigned to the synapse alone show less signal (rate ratio = 1.60), although these compartments also have fewer genes (26 for synaptic membrane; 22 for synaptic cleft; and 7 for extrasynaptic space). In contrast, for BP (**Fig. S21**), comparisons show greater differences.

Given the variation in major child compartments for BP's "process in synapse", we next contrast mutation rates in cases only in these unique gene sets (**Table S18**). To do so, we first compute the rate of mutations found in cases per set (total number of mutations over cases divided by number of genes); then compute the ratio of these rates for independent gene sets; as well as a *P* value of the contrasted rates under a binomial assumption. The highest mutation rate was observed for genes shared between synaptic signalling and organisation (4.69), followed by those shared between pre- and postsynaptic compartments (4.62). However, this pattern was not universal: genes shared between postsynaptic and signalling showed only marginally elevated rates relative to either alone (1.48 versus 1.24 and 1.18, respectively).

To investigate the differing patterns for presynaptic versus postsynaptic between independent sets for CC versus BP, we further partitioned all SynGO genes into BP-only ($n$ = 108 genes), CC-only ($n$ = 457), and both CC and BP ($n$ = 968 genes) sets. The CC-only set demonstrated higher ASD enrichment compared to BP (CC-only: RR = 2.72, $Z_{all}$ = 7.43, $P = 1.1\times10^{-13}$; BP-only: RR = 1.99, $Z_{all}$ = 3.12, $P$ = 0.0018; CC and BP: RR = 2.36, $Z_{all}$ = 11.55, $P = 7.6\times10^{-31}$). This CC signal was driven by a concentration of calcium and potassium ion channel genes, including *CACNA1C*, *CACNB1*, *KCNB1*, and *KCNQ2*, as well as *BCL11A*, *SCN8A*, and *PTEN* (52 mutations in probands, 1 in controls), all of which are categorized as postsynaptic in SynGO. This pattern explains why the signal is stronger for presynaptic (or both pre- and postsynaptic) in the BP gene set than it is for CC, in which signal is stochastically equal across pre- and postsynaptic gene sets.

### Gene set DNM burden analyses stratifying by DD/ID status

***Methods***

To quantify the contributions of ASD genes to damaging DNM burden across biologically defined gene sets, and to examine whether these contributions differ by DD/ID comorbidity status, we computed observed-to-expected (O/E) ratios for DNMs across three proband groups: all ASD probands ($N$ = 38,680), ASD probands with recorded DD/ID ($ASD_D$, $n$ = 24,839) and those without ($ASD_{DA}$, $n$ = 13,841). Unaffected siblings ($N$ = 9,567) served as a negative control. As with the SynGO analyses, variant classes included PTV, Mis2, and Mis1.

Expected mutation counts were derived from per-gene, per-variant-class mutation rates computed by annotating all exonic SNVs in the GRCh38 reference genome with per-site mutation rates from gnomAD v4.1, stratified by trinucleotide sequence context, allele, and methylation level, using the same variant consequence definitions applied to the ASD cohort. Rates were summed by gene and variant class, excluding autosomal sites with mean gnomAD genome coverage below 10x (and non-PAR chrX/Y sites below 1x; full details: https://github.com/ksatterstrom/ASD-Genetics/blob/main/Mutation_rate_calculations_for_ASC_2024-07-16_24.ipynb). For each gene set and cohort, the expected count was $E = \Sigma \mu_i * 2 * N$, where $\mu_i$ is the sum of mutation rates across all PTV, Mis2, and Mis1 sites in gene $i$, and $N$ is the number of individuals in the cohort.

The O/E ratio was computed as O/E = Σobs / Σ$E$, with exact 95% confidence intervals derived using the Garwood method [4]. Pairwise differences in DNM enrichment were assessed using a two-sided conditional binomial test, in which the null probability $E_A / (E_A + E_B)$ reflects the expected proportion of mutations attributable to gene set A under equal enrichment. Rate ratios (RRs) were estimated as $(obs_A / E_A) / (obs_B / E_B)$ to compare O/E ratios between cohorts for each gene set, with 95% confidence intervals estimated by the log-normal delta method.

The spatial burden analysis used gene sets derived from a spatial transcriptomics factor analysis of the prenatal human brain [14]. Genes were retained if they satisfied a robust expression criterion and were enriched in factors corresponding to three brain regions: germinal zone (Factor 2; $n$ = 26 genes), cortical plate (Factor 3; $n$ = 5 genes), and thalamus (Factor 5; $n$ = 44 genes). Factors corresponding to subplate (Factor 1) and caudate/putamen (Factor 4) were excluded due to their small sizes ($n$ = 2 genes). For genes with loadings across multiple retained factors, each gene was assigned to the factor with the highest loading. The genes with identical maximum loadings across factors (*ADNP*, *TLE3*, *WAC*) were excluded from all gene sets. Pairwise comparisons of O/E ratios between gene sets within each cohort were performed to estimate their relative burden.

***Spatial enrichment stratifying by DD/ID status***

First, given that the cell-type enrichment analyses were limited to cortical cells from V1 and PFC, we asked whether ASD vulnerability extends to the developing thalamus, a major source of thalamocortical input to these regions [18]. We quantified the burden of DNMs in spatially-enriched gene sets from a prenatal spatial transcriptomic atlas [14], including germinal zone (GZ, $n$ = 26), cortical plate (CP, $n$ = 5), and thalamus (THL, $n$ = 44) gene sets overlapping our 253 ASD genes (**Table S15A**). All three gene sets showed significantly higher O/E ratios and DNM burden in ASD probands compared to unaffected siblings (**Fig. S27A; Table S15B**).

Stratifying by DD/ID status, all three gene sets were significantly enriched in both $ASD_D$ and $ASD_{DA}$ probands relative to siblings (**Fig. S27B**). The CP-enriched gene set showed strongest burden in both $ASD_D$ and $ASD_{DA}$ probands relative to siblings. CP-enriched and thalamus-enriched genes showed a trend toward greater enrichment in $ASD_D$ than $ASD_{DA}$ probands, compared to GZ-enriched genes, though confidence intervals overlapped across regions (**Table S15C**). Pairwise comparisons of O/E ratios across spatial gene sets revealed that in $ASD_D$ probands, both CP-enriched genes and thalamus-enriched showed significantly higher burden than GZ-enriched genes (**Fig. S27C**; **Table S15D**). Conversely, no significant differences were observed between any spatial gene sets in $ASD_{DA}$ probands, suggesting THL-enriched genes pose similar risk to ASD without comorbid DD/ID to GZ- and CP-enriched genes.

***Module enrichment stratifying by DD/ID status***

We next examined whether the modules associated with ASD rare variant were preferentially enriched in probands with or without DD/ID. Of the 28 independently associated modules, 28 showed significantly elevated DNM burden in $ASD_D$ probands relative to $ASD_{DA}$ probands, with relative risk ratios ranging from 1.5 to 4.0 (all FDR < 0.05 except for *ZNF281*$^{(+)}$; **Fig. S28A-B**). This DDID-predominant pattern held across all 1,151 modules, as 564 showed significantly higher DNM burden in $ASD_D$ than $ASD_{DA}$ probands ($RR_{D:DA}$ > 1, FDR < 0.05), while only 2 showed the reverse ($RR_{D:DA}$ < 1, FDR < 0.05), indicating that $ASD_D$ probands carry greater rare variant burden broadly across co-expression and regulon gene programs (**Fig. S28C-D**; **Table S16A-B**). The most significant $ASD_D$ enrichments relative to $ASD_{DA}$ were observed in *CUX1*$^{(+)}$, *VEZF1* (M11-iso), and *NR3C1*$^{(+)}$, all three of which are characterized by predominantly genomic regulation (GR) pathways as determined by topic modeling. *TCF4*$^{(+)}$, a small 14-gene regulon including established ASD genes *NRXN1*, *ARID1B*, and *SETD5*, had the highest $O/E_D$ and $ASD_D$:$ASD_{DA}$ RRs ($O/E_D$ = 19.1 [15.2, 23.4]; $RR_{D:DA}$ = 4.04 [2.78, 5.86]; FDR = $1.76\times10^{-14}$; **Fig. S28A**).

While $ASD_D$ burden was significantly greater than $ASD_{DA}$ burden across virtually all modules (**Fig. S28C-D**), $ASD_{DA}$ probands also showed significant enrichment above siblings for 26 of the 28 independently associated modules, with rate ratios of $ASD_{DA}$ vs siblings [$RR_{DA:sib}$] ranging from 1.36 to 7.70 (all FDR < 0.05 except *ZNF281*$^{(+)}$, *MZF1*$^{(+)}$, and *LHX9*$^{(+)}$; **Fig. S28A-B**; **Table S16B**). This finding indicates that these gene programs capture ASD vulnerability across the full phenotypic spectrum, not exclusively in probands with comorbid DD/ID. The strongest $ASD_{DA}$ enrichments (quantified as rate ratio vs siblings) were observed in *TCF4*$^{(+)}$, *BHLHE22*$^{(-)}$, *SRCIN1* (M84-gene), *SOX11*$^{(+)}$, and *THRB*$^{(-)}$, though confidence intervals were wide for *TCF4*$^{(+)}$ and *BHLHE22*$^{(-)}$, reflecting their small gene set sizes (*n* = 14 and *n* = 30, respectively). Notably, four of these five modules were enriched in the MORPH category (excepting *SRCIN1*, which was SYN), in contrast to the GR-dominant enrichment observed among the most higher-DDID modules, suggesting that neuronal morphogenesis and differentiation pathways are disproportionately implicated in ASD vulnerability without comorbid DD/ID.

***SynGO enrichment stratifying by DD/ID status***

Finally, we asked whether synaptic gene ontology enrichments differ by DD/ID comorbidity status by performing O/E analysis across all SynGO pathways (**Fig. S29A**; **Table S17A-B**). Overall, probands with comorbid DD/ID carried ~53.5% of synaptic mutations, with $ASD_D$ probands showing a 2-fold higher mutation rate than $ASD_{DA}$ probands. Despite the lower mutation rate for $ASD_{DA}$, these individuals still showed a 1.81-fold and 1.94-fold higher mutation rate than controls for genes in BP (RR = 1.81, $Z_{all}$ = 7.76, $P = 8.3\times10^{-15}$) and CC (RR = 1.94, $Z_{all}$ = 9.38, $P = 6.7\times10^{-21}$), supporting the finding that synaptic machinery is disrupted across the full ASD phenotypic spectrum.

A core set of presynaptic and postsynaptic pathways was significantly enriched in both $ASD_D$ and $ASD_{DA}$ probands relative to siblings. As expected given the higher rate of mutations in $ASD_D$

probands, the majority of SynGO pathways significant in either cohort showed greater enrichment in $ASD_D$ than $ASD_{DA}$ probands, as quantified by the $RR_{D:DA}$ of the O/E ratio (consistent with the broader pattern observed across the GO database; **Fig. S29B**). These pathways spanned both presynaptic and postsynaptic compartments, the three most significant being integral component of presynaptic membrane (GO:0099056; $O/E_D$ = 6.56 [5.78, 7.43]; $RR_{D:DA}$ = 2.60 [2.15, 3.16]; FDR = $1.70\times10^{-20}$), postsynaptic density intracellular component (GO:0099092; $O/E_D$ = 9.61 [8.14, 11.3]; $RR_{D:DA}$ = 3.45 [2.63, 4.53]; $RR_{D:DA}$ = $2.00\times10^{-18}$), and modulation of chemical synaptic transmission (GO:0050804; $O/E_D$ = 6.12 [5.25, 7.09]; $RR_{D:DA}$ = 3.01 [2.36, 3.83]; FDR = $7.02\times10^{-18}$; **Table S17A**).

Synapse adhesion between pre- and post-synapse (GO:0099560) was the only pathway to show significant enrichment in $ASD_{DA}$ probands but not $ASD_D$ probands relative to siblings ($O/E_{DA}$ = 2.24 [1.60, 3.05]; FDR = $3.90\times10^{-5}$; $O/E_D$ = 1.51 [0.84, 2.49]; FDR = 0.16; **Fig. S29A**; **Table S17A**), suggesting that trans-synaptic adhesion represents a distinct axis of ASD vulnerability independent of DD/ID comorbidity. This was replicated across the broader GO database (BP, CC, and MF ontologies; **Fig. S29B**), in which synapse membrane adhesion was again the only pathway showing significant enrichment in $ASD_{DA}$ but not $ASD_D$ probands relative to siblings, though the DDID-predominant pattern remained evident across the vast majority of GO pathways.

Together, these findings indicate that rare variant burden is broadly elevated in $ASD_D$ relative to $ASD_D$ probands across module, synaptic, and spatial gene sets; however, a small number of pathways related to neural circuit organization and trans-synaptic connectivity show preferential enrichment in ASD without comorbid DD/ID.

### Supplementary Figures

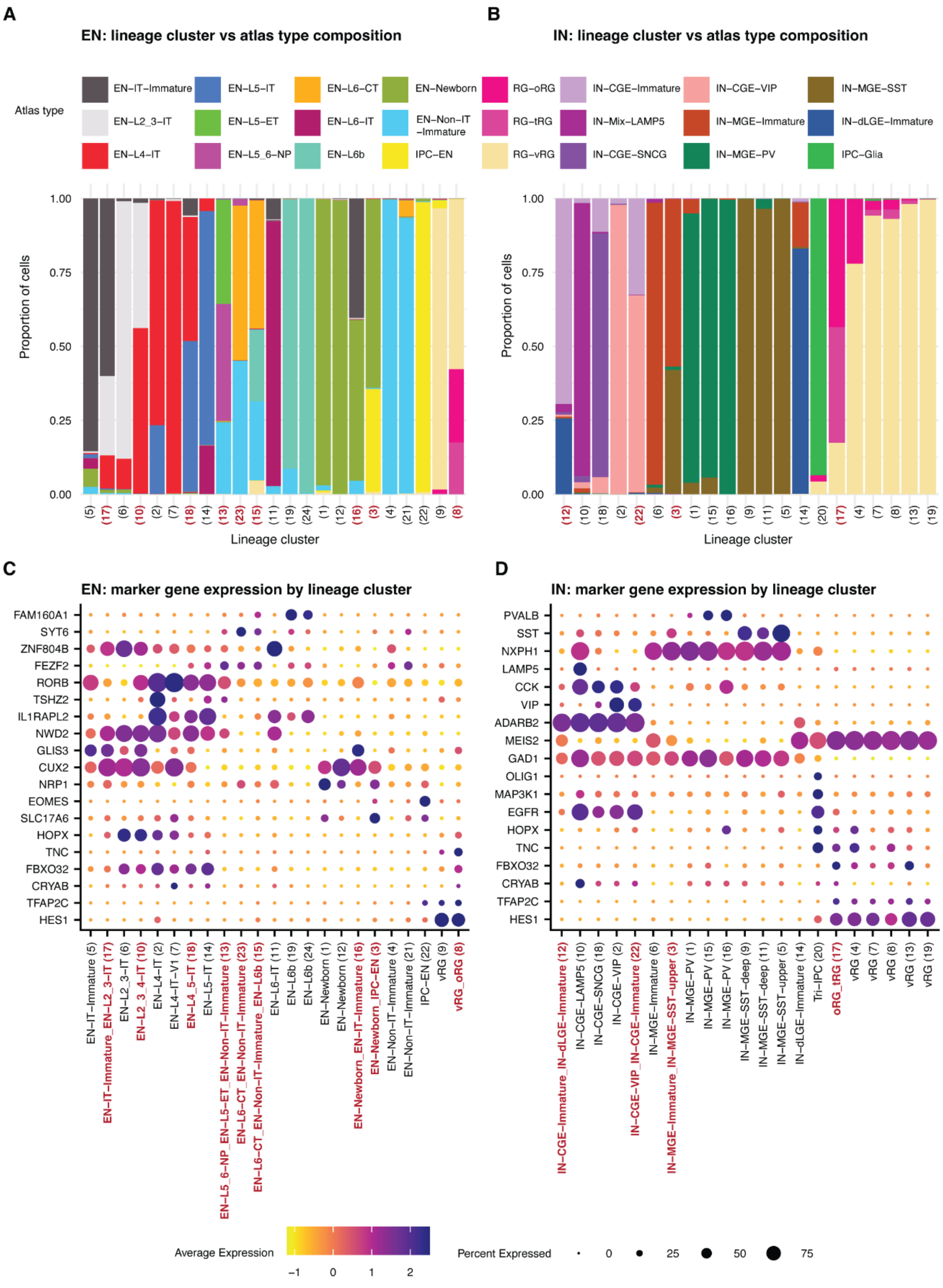

**Fig. S1. Annotation of excitatory neuron (EN) and inhibitory neuron (IN) lineage clusters.**
**(A,B)** Proportion of cells assigned to each snMultiome atlas cell-type annotation [19] within each EN **(A)** and IN **(B)** lineage cluster, identified via *mclust* clustering in the 8D WNN-UMAP embedding (see Methods). Cluster numbers are indicated in parentheses below each bar; numbers shown in red denote clusters

assigned a combined label (no single atlas type exceeded 70% of constituent cells). **(C,D)** Average expression and percentage of cells expressing marker genes across EN **(C)** and IN **(D)** lineage clusters, labeled by their composition-based names (cluster numbers in parentheses, matching A and B). Cluster names shown in red, bold text indicate clusters assigned a combined label.

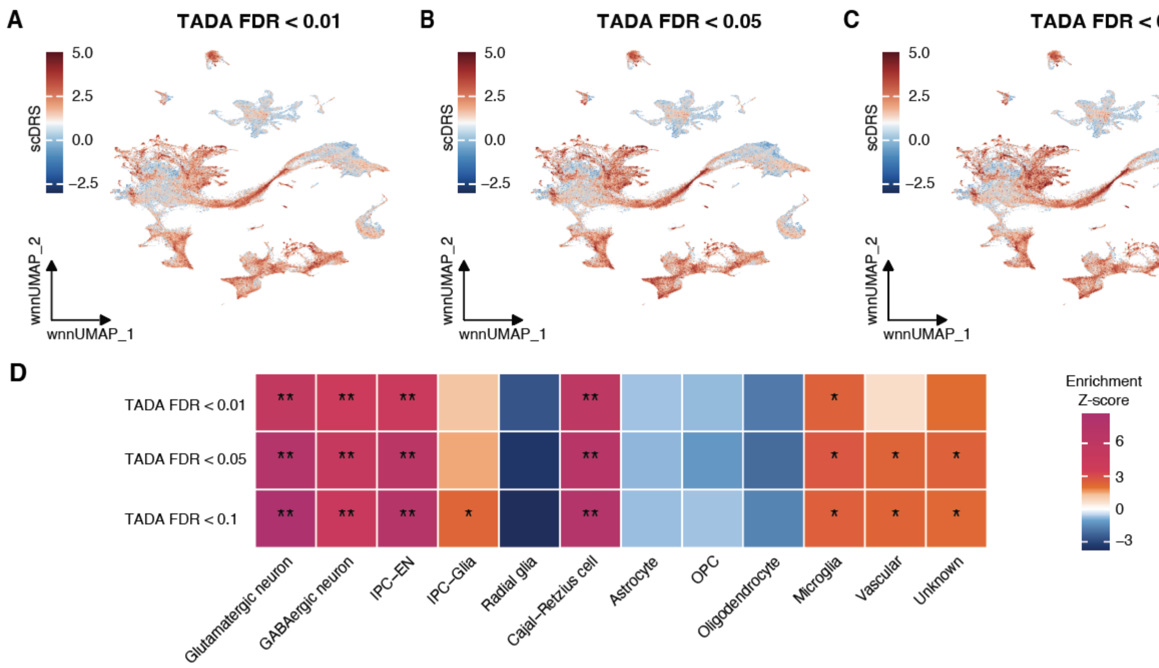

**Fig. S2. Sensitivity analyses of cell-type enrichment using expanded ASD gene sets.**
**(A-C)** UMAP visualizations of normalized scDRS scores using ASD genes at TADA FDR thresholds: FDR < 0.01 ($n$ = 416; A), FDR < 0.05 ($n$ = 696; B), and FDR < 0.1 ($n$ = 951; C). **(D)** Enrichment $Z$-scores for each subclass. Significance: ** FDR < 0.01; * FDR < 0.05.

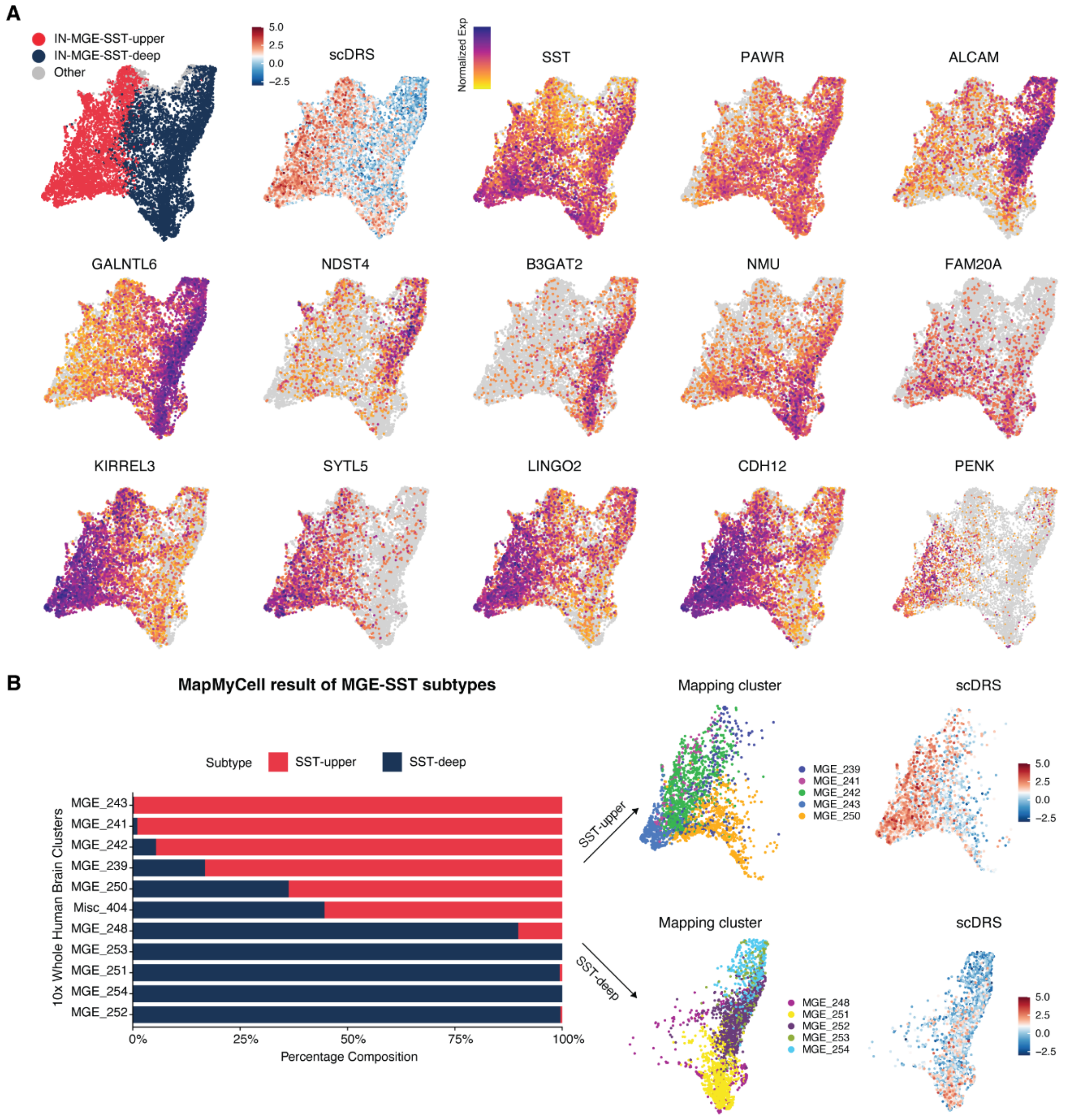

**Fig. S3. Differential ASD enrichment within the IN-MGE-SST cluster.**
**(A)** UMAP of IN-MGE-SST cells colored by subtype identity, normalized scDRS, and normalized expression of representative marker genes. **(B)** Mapping MGE-SST subclusters to the 10x Whole Human Brain Atlas via MapMyCell[1] uncovered subtype-specific molecular signatures. Left: Proportional composition of SST-upper and SST-deep subtypes across major 10x Whole Human Brain Atlas clusters. Right: UMAP of the five clusters most strongly enriched for SST-upper and SST-deep.

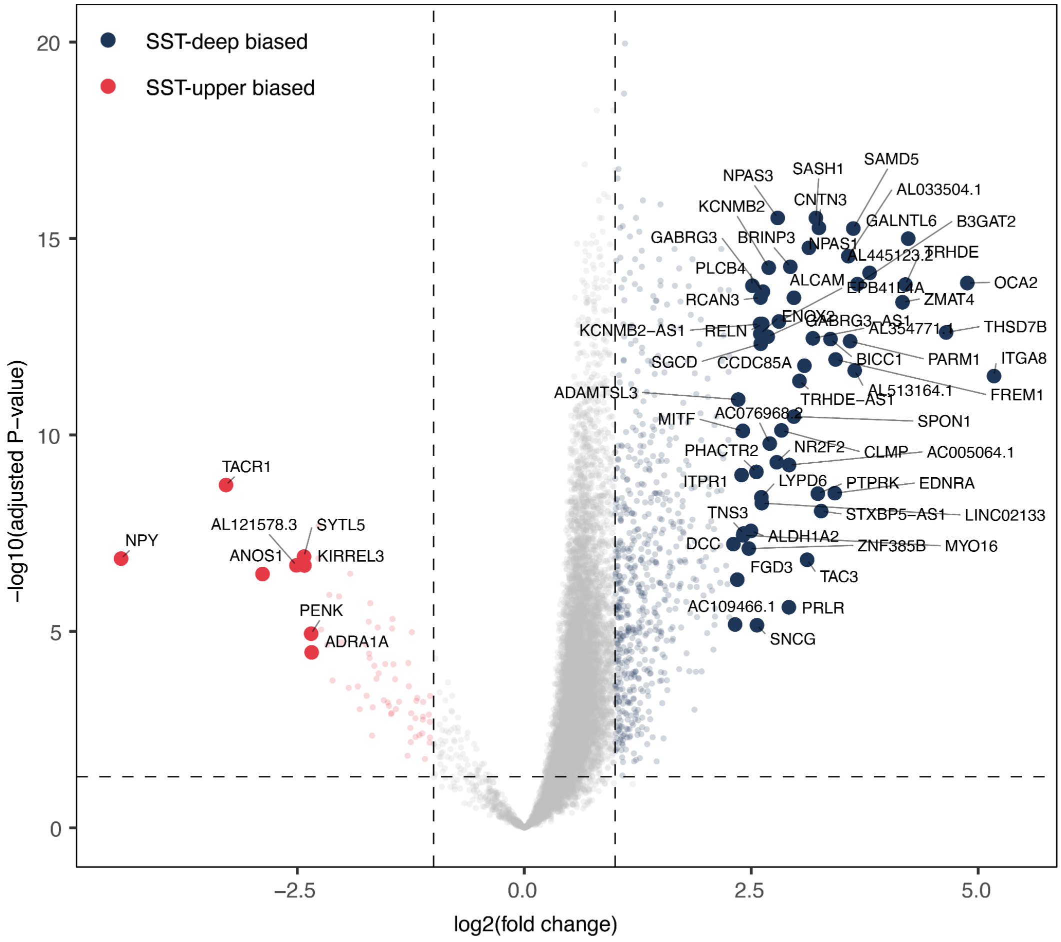

**Fig. S4. Differential gene expression between MGE-SST upper-layer and deep-layer subtypes.** Genes with |$\log_2$ fold change| > 2.3 and FDR < 0.005 are highlighted in the volcano plot.

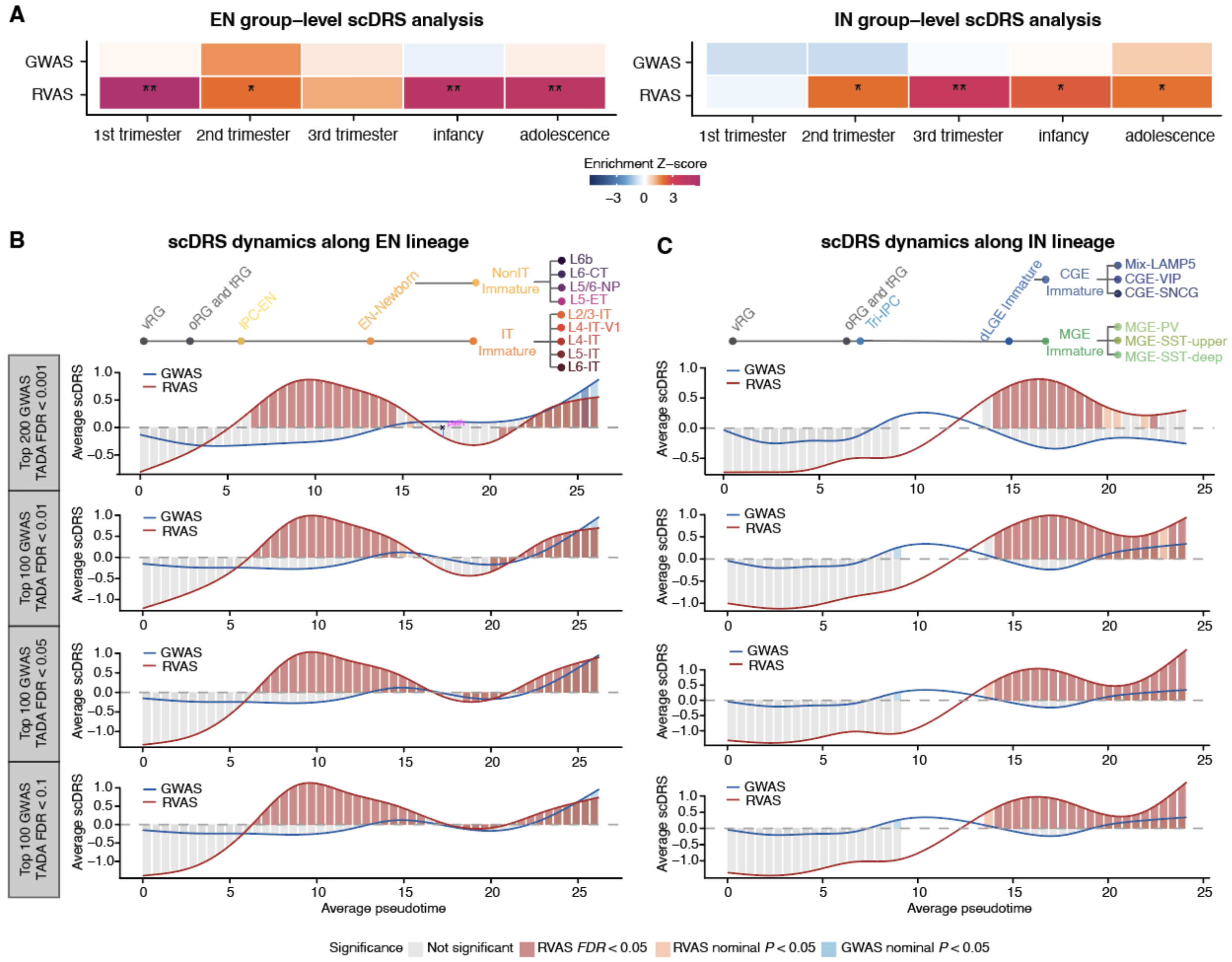

**Fig. S5. Developmental and pseudotemporal enrichment of ASD vulnerability in excitatory neuron (EN) and inhibitory neuron (IN) lineages.**

**(A)** Group-level scDRS enrichment of rare (RVAS) and common (GWAS) variant-based gene sets across developmental stages. Heatmaps showing enrichment Z-scores from group-based scDRS analysis for RVAS (rare variant) and GWAS (common variant) gene sets across five developmental stages in the excitatory neuron (EN, top) and inhibitory neuron (IN, bottom) lineages. Significance: * FDR < 0.05, ** FDR < 0.01. **(B-C)** Sensitivity analysis for developmental stage enrichment of ASD genes. Temporal comparison of RVAS and GWAS enrichment along EN (**B**) and IN (**C**) pseudotime. Schematic ordering of lineage states along the pseudotime axis with corresponding cell type annotations. Average scDRS per pseudotime bin for RVAS and GWAS, curve color indicates bin-level significance.

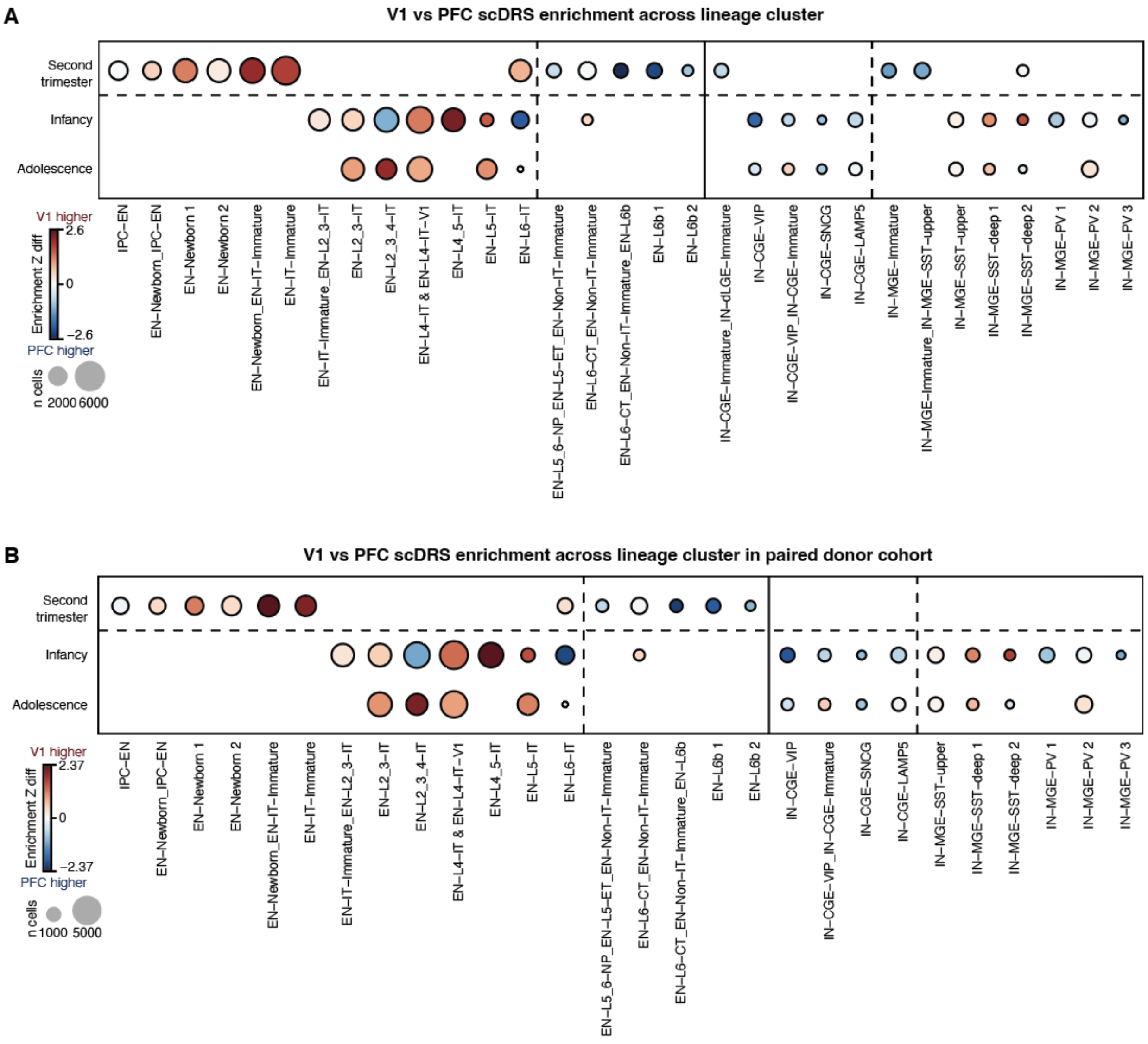

**Fig. S6. Group-level scDRS enrichment difference between V1 and PFC, by lineage cluster and developmental stage.**

**(A)** Full dataset. **(B)** Paired-donor subset, restricted to donors with matched PFC and V1 samples, to control for inter-individual variability. Each point represents one lineage cluster (EN or IN) at one developmental stage (T2, second trimester; Infancy; Adol, adolescence; third trimester excluded due to sparse data). The shared EN-L4-IT cluster and the region-specific EN-L4-IT-V1 cluster were pooled into a single group for this analysis. Point color indicates the scDRS group-level enrichment Z-score difference between V1 and PFC (Z_V1 − Z_PFC); point size indicates the total number of cells (PFC + V1 combined). Only cluster × stage × region combinations with ≥150 cells in both PFC and V1 are shown. Positive values (red-toned) indicate V1-biased enrichment; negative values (blue-toned) indicate PFC-biased enrichment.

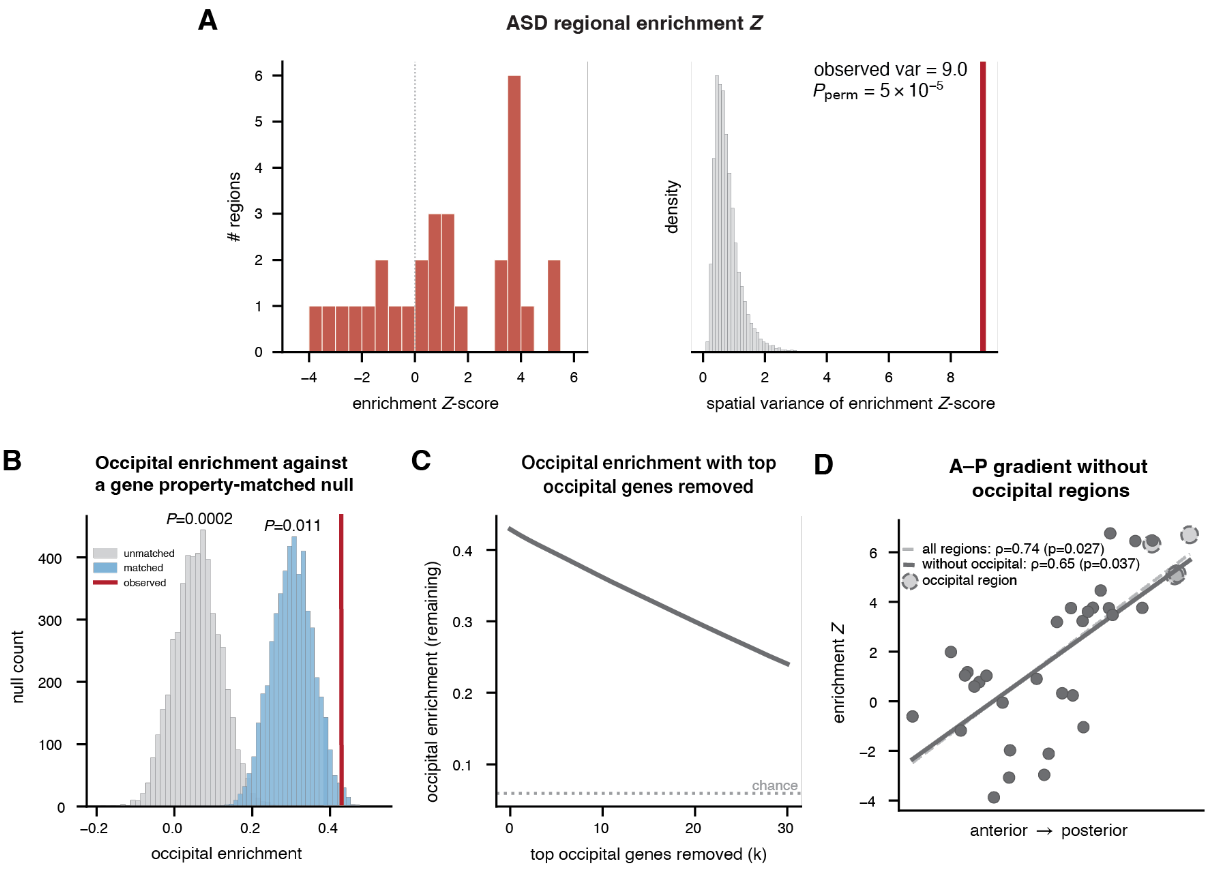

**Fig. S7. The ASD anterior-posterior enrichment gradient is robust to spatial and gene property confounds.**

**(A)** The ASD gene spatial enrichment is not uniform across the cortex. Left: Distribution of regional enrichment *Z*-scores across the 34 cortical Desikan-Killiany regions. Right: The spatial variance of enrichment *Z* across the 34 regions (red line = observed enrichment variance) compared with 20,000 random size-matched sets. The observed variance exceeds every null set ($P_{perm} = 5\times10^{-5}$), indicating that ASD enrichment is regionally patterned rather than uniform across the cortex. **(B)** Occipital enrichment remains above chance as the genes with the highest expression in the occipital regions are removed, indicating the signal is not driven by a handful of high-effect genes. **(C)** Occipital enrichment (red line) exceeds both an unmatched null (gray) and a null matched on gene expression, coding sequence length, and constraint (LOEUF; blue), indicating the enrichment effect extends beyond these gene properties. **(D)** The A-P gradient is robust to the exclusion of four occipital regions (pericalcarine, lateral occipital, lingual, cuneus), suggesting it is a cortex-wide property. Dashed line: full $\rho = 0.74$, $P_{spin} = 0.027$; Solid line: occipital-removed $\rho = 0.65$, $P_{spin} = 0.037$.

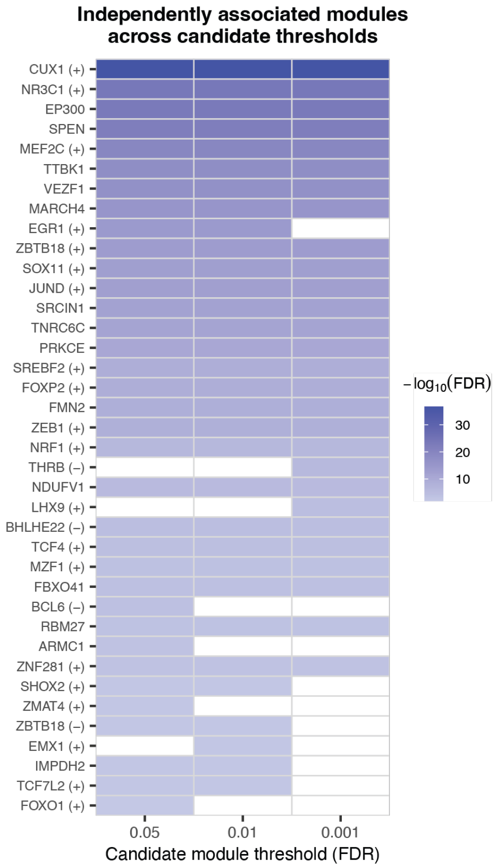

**Fig. S8. The set of independently associated modules is robust to the candidate module significance threshold.**
Each column represents a stepwise conditional analysis run with a different candidate inclusion threshold (FDR < 0.05, 0.01, or 0.001), holding the conditional selection threshold fixed (likelihood-ratio $P < 0.001$). Rows are all modules that were independently associated with ASD rare variant vulnerabilities in at least one run, labeled by their top hub gene (co-expression modules) or transcription factor (regulons). Cell fill encodes each module's marginal association strength ($-\log_{10}(\mathrm{FDR})$); white indicates a module not selected at that threshold. Rows are ordered by marginal significance (most significant at top). Because the marginal statistic is independent of the candidate threshold, a given module's shade is constant across the columns in which it is retained; consistency across columns therefore reflects stability of the selected set.

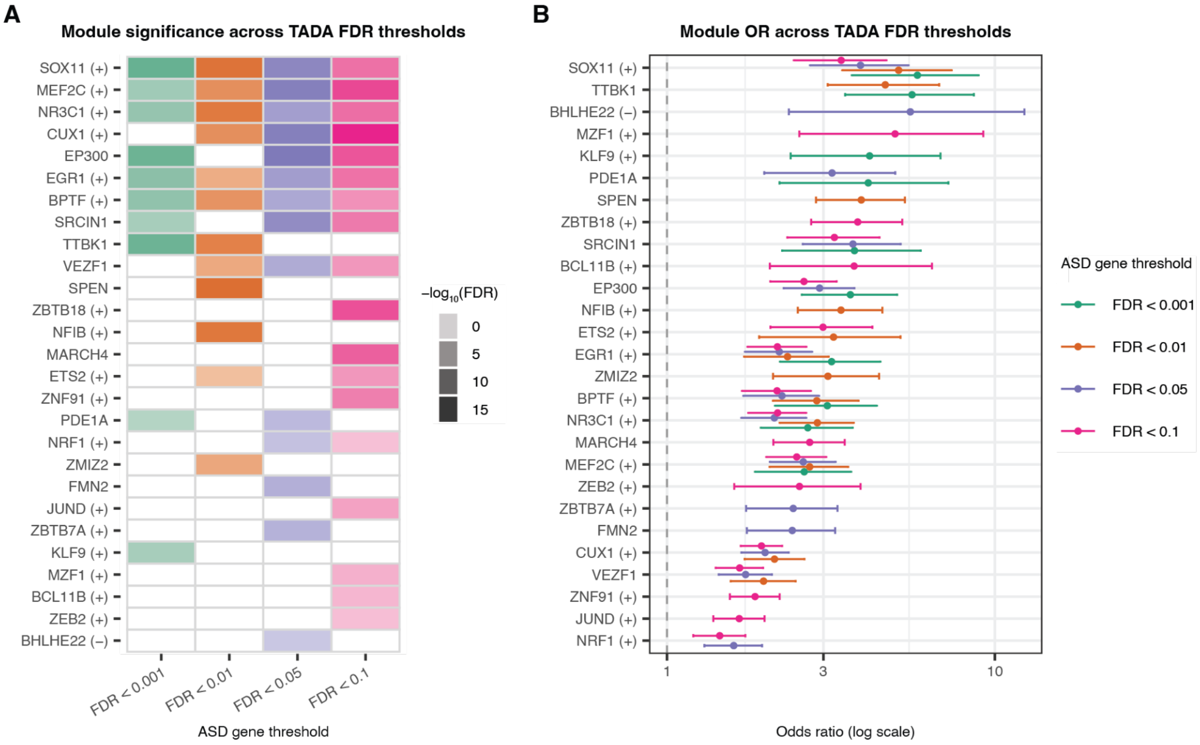

**Fig. S9. Module curation using logistic regression across TADA FDR thresholds.**
**(A)** Significance of association (shown as $-\log_{10}$(FDR)) for each module across four ASD gene definition thresholds. White cells indicate modules not independently associated at that threshold. **(B)** Odds ratios and 95% CIs for all modules independently associated with ASD at any of the four FDR thresholds, shown on a log scale. Each module is shown once per threshold at which it was independently significant, with points color-coded by threshold.

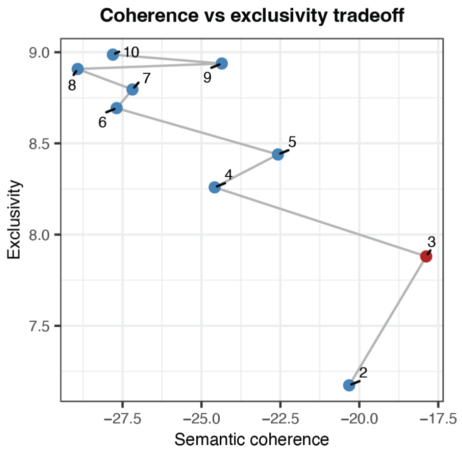

**Fig. S10. Selection of the number of topics (K) for the gene module topic model.**
Semantic coherence (i.e., how often high-probability words occur in the same document) and exclusivity (i.e., the degree to which high-probability words in a topic are unique to that topic) are shown for topic models fitted with $k$ = 2-10 topics (label = $k$). $k$ = 3 maximizes semantic coherence among $k$ = 2-10, exclusivity increases monotonically with $k$ and so never identifies an interior optimum, and the final choice was based on the interpretability of the topics.

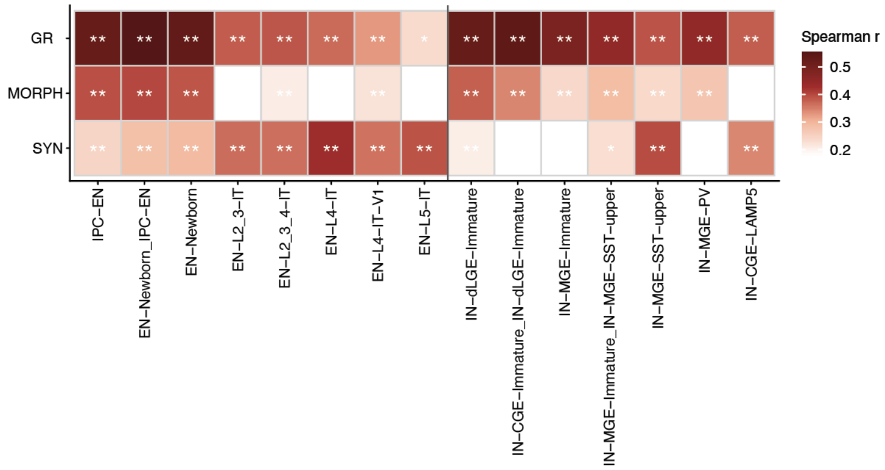

**Fig. S11. Correlation between cell-level biological program AUC scores and scDRS across ASD relevant lineage clusters.**

Network cell-type associations mirror expression trajectories, with SYN network genes associated with mature neurons and GR and MORPH networks enriched for immature cell types. Significance assessed by two-sided Monte Carlo test: * FDR < 0.05; ** FDR < 0.01.

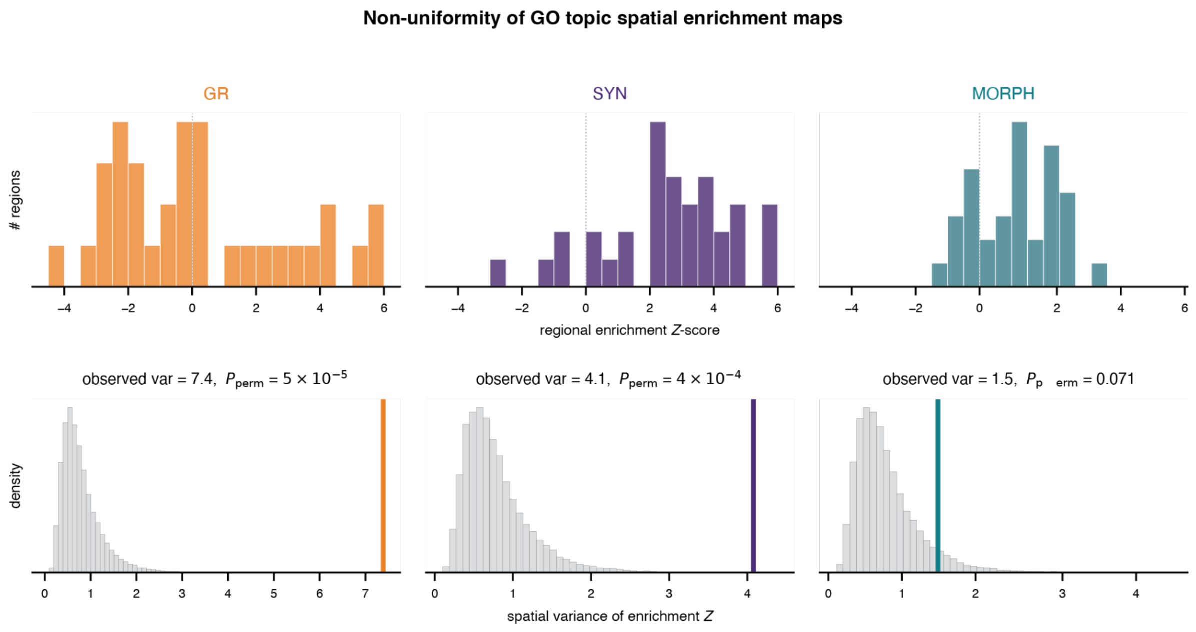

**Fig. S12. Cortical enrichment of the ASD biological programs is spatially non-uniform.**
For each GO-derived program, regional enrichment across the 34 cortical Desikan-Killiany regions was compared to 20,000 random size-matched gene sets. Top: distribution of regional enrichment *Z*-score. Bottom: the spatial variance of enrichment *Z* across the 34 regions (colored line = observed) compared with the null distribution of variances from the same 20,000 random gene sets; $P_{perm}$ is the fraction of random sets whose variance meets or exceeds the observed value. GR and SYN enrichment is significantly non-uniform ($P_{perm} = 5\times10^{-5}$ and $4\times10^{-4}$), whereas MORPH is borderline ($P_{perm} = 0.07$).

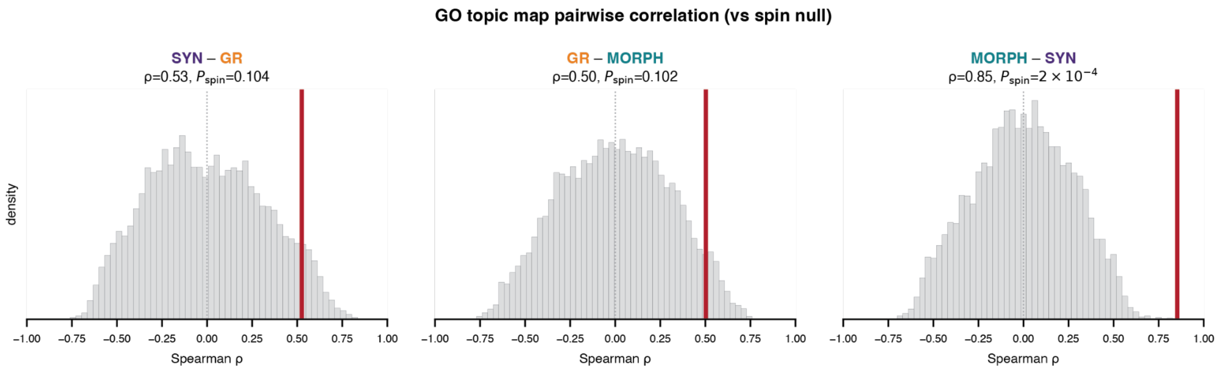

**Fig. S13. SYN and MORPH cortical enrichment patterns are spatially correlated beyond chance, while GR is less related.**

Shown are the pairwise Spearman correlations between the biological program cortical enrichment maps across the 34 Desikan-Killiany cortical regions (red line), each compared to spatial autocorrelation-preserving null distribution (5000 rotations). SYN and MORPH are strongly and significantly correlated ($\rho$ = 0.85, $P_{spin}$ = 2×10$^{-4}$), whereas GR correlates moderately with each and not beyond spatial autocorrelation (SYN-GR $\rho$ = 0.53, $P_{spin}$ = 0.10; GR-MORPH $\rho$ = 0.50, $P_{spin}$ = 0.10).

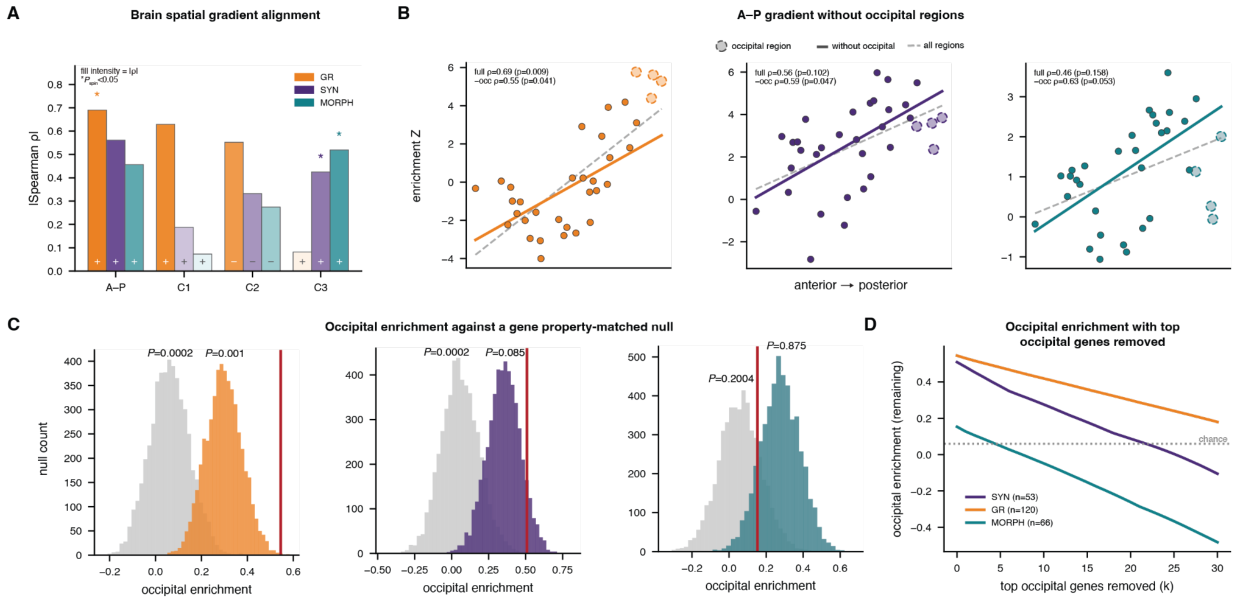

**Fig. S14. Program-specific cortical enrichment gradients are robust, with GR showing the strongest posterior bias.**
**(A)** Alignment of each program's spatial enrichment map with the A-P axis and expression components C1-C3 [10]. Bar height and fill intensity indicate the magnitude of Spearman correlation, +/- indicate direction of correlation, * $P_{spin}$ < 0.05. **(B)** Posterior enrichment in GR persists after the removal of occipital regions; SYN is marginally significant after occipital removal and MORPH not at all, indicating weaker, occipitally-influenced gradients. *P* values are spin corrected using 1000 rotations. **(C)** Occipital enrichment relative to unmatched (gray) and gene property-matched (program color) nulls. GR is strongly enriched when corrected for gene properties; SYN and MORPH are not. **(D)** Occipital enrichment in the GR program is distributed across many genes, whereas MORPH is not enriched above chance. SYN shows some distributed occipital enrichment, but this is likely due to spatial autocorrelation (panel A) and gene properties (panel C).

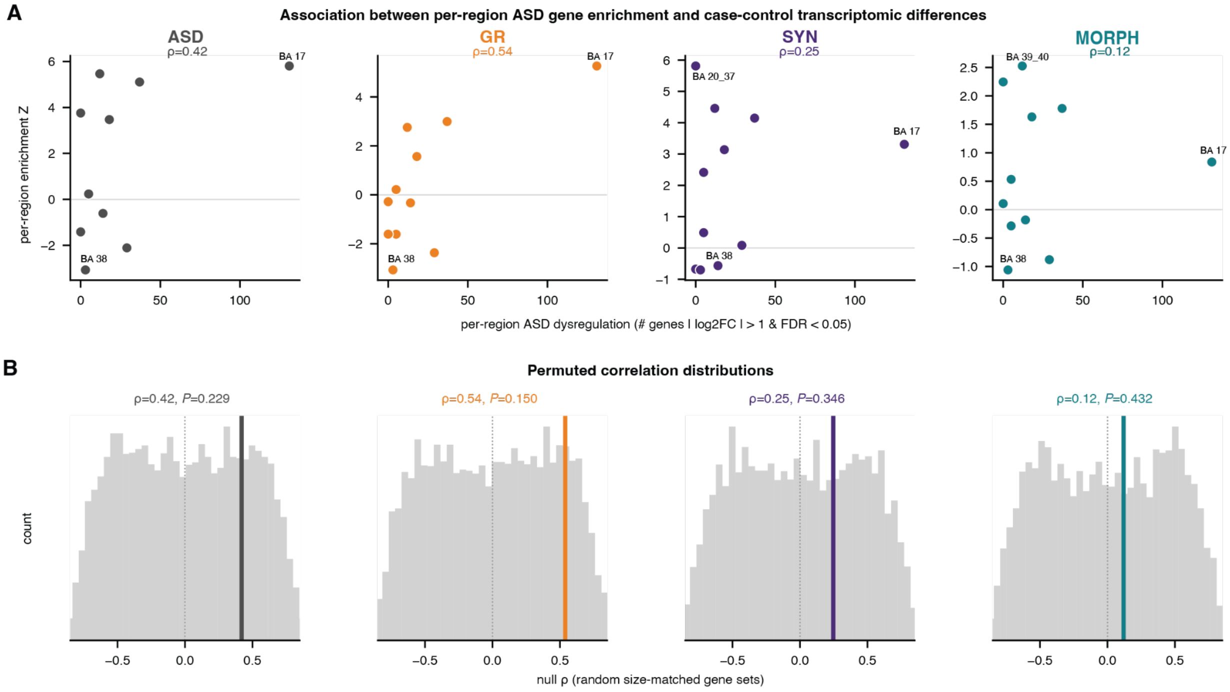

**Fig. S15. ASD gene spatial enrichment is directionally concordant with regional cortical transcriptomic dysregulation in ASD.**

**(A)** In GR-assigned genes (and to some extent, ASD overall), per-region ASD gene enrichment tracks per-region ASD dysregulation across (genes with $|\log_2 FC| > 1$ and FDR < 0.05; ASD vs control) across 11 Brodmann regions[11], peaking in primary visual cortex (BA17). **(B)** Against a null of size-matched random random gene sets, the GR and overall-ASD correlations are directionally consistent but do not reach significance at $n = 11$ regions (one-sided permutation $P = 0.15$ and 0.23, respectively).

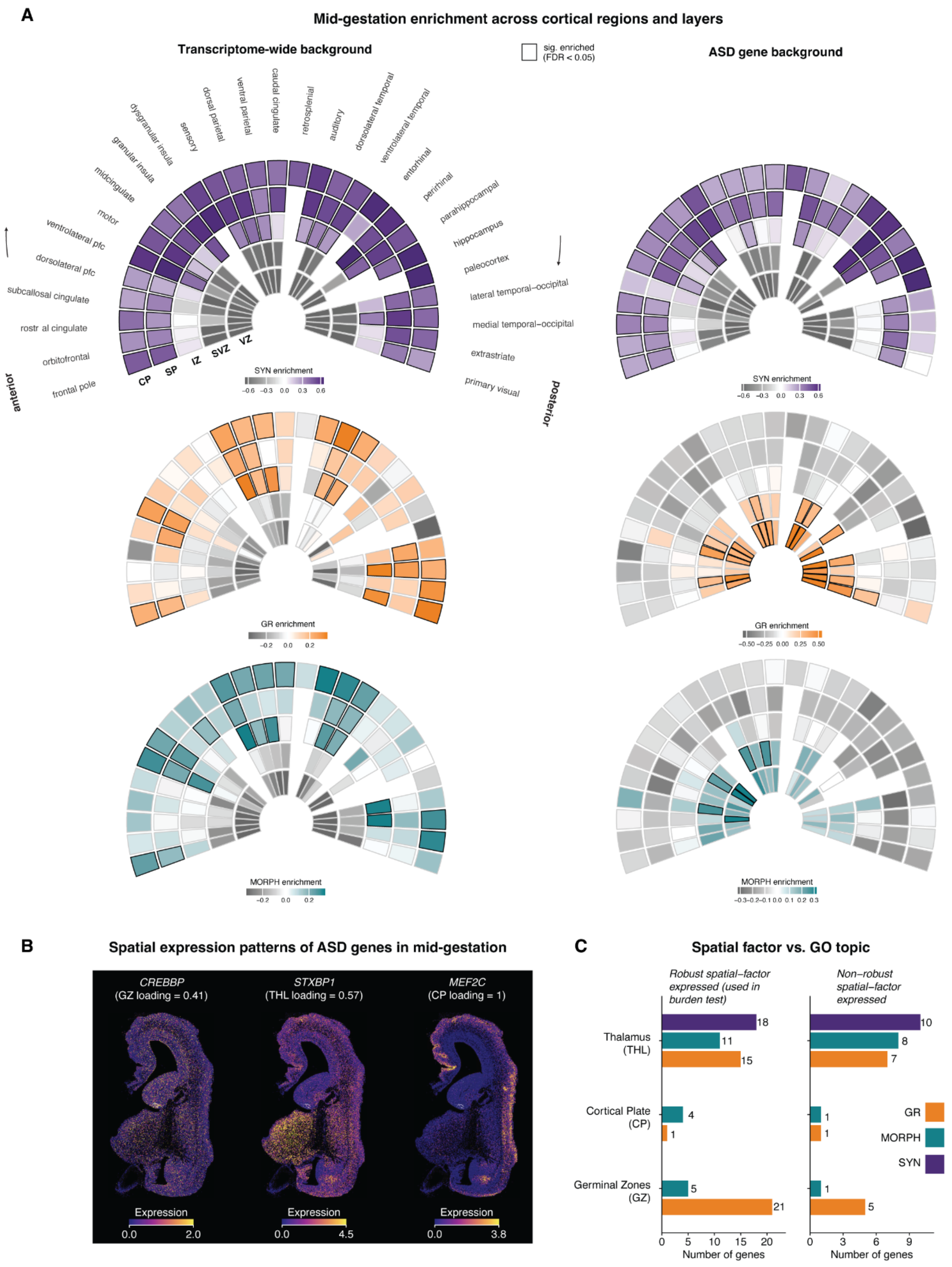

**Fig. S16. Prenatal spatial enrichment patterns of ASD genes by biological program.**

**(A)** Regional and laminar enrichments of per-program ASD genes across the prenatal cortex. Each wedge represents one laminar zone x cortical region, with the radius indicating depth (ventricular zone [VZ], subventricular zone [SVZ], intermediate zone [IZ], subcortical plate [SP], cortical plate [CP]) and angle indicating region ordered anterior-to-posterior. Fill is the enrichment of the program's genes relative to all expressed genes (left) and other ASD genes (right), estimated using a linear mixed effects model. Wedges that are significantly enriched above background (BH FDR < 0.05) are outlined in black. **(B)** Spatial expression of representative ASD genes in the prenatal human brain from the STAGE atlas (stageatlas.org, [14]). **(C)** Distribution of ASD genes across GZ, CP, and THL spatially enriched gene sets. Genes were classified as exhibiting robust or non-robust spatial-factor expression according to their maximal observed expression across regions [14].

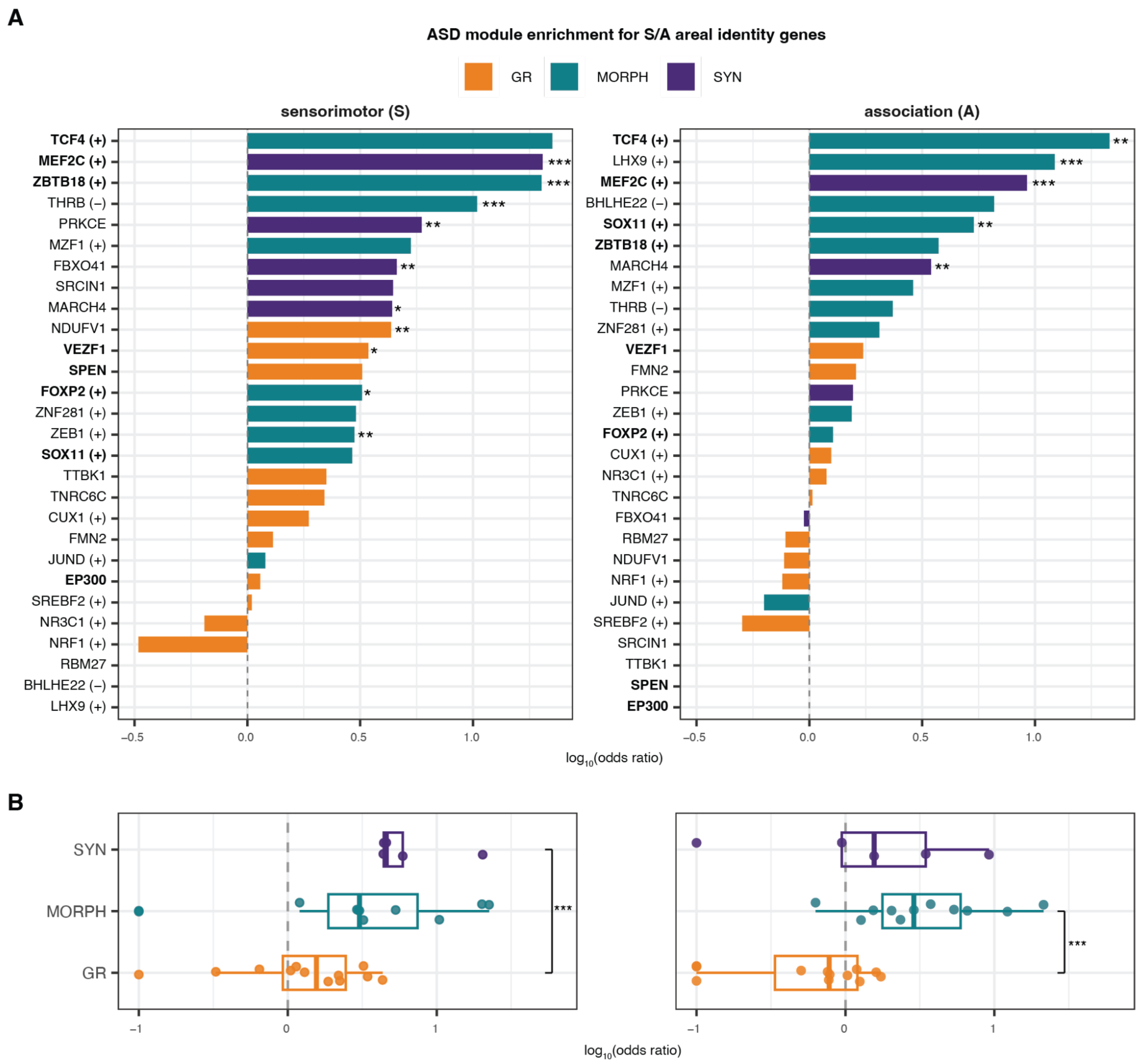

**Fig. S17. Enrichment of ASD modules for sensorimotor-association (S-A) areal identity genes.** All ASD-associated co-expression modules and regulons were tested for hypergeometric overlap with the sensorimotor (S; *N* = 43 genes) and association (A; *N* = 171 genes) gene sets defined by Tsyporin et al. [20]. **(A)** Per-module enrichment for the S (left) and A (right) gene lists, shown as $\log_{10}$(odds ratio) from a one-sided hypergeometric test. The dashed line marks OR = 1 (no enrichment); asterisks denote Benjamini–Hochberg FDR computed across all modules within each gene list (*FDR < 0.05, **FDR < 0.01, ***FDR < 0.001); bold module labels indicate modules whose hub gene or transcription factor is itself an ASD gene. **(B)** Distribution of module enrichments grouped by biological program for the S and A lists. Brackets show pairwise two-sided Wilcox rank-sum test between programs (*$P$ < 0.05, **$P$ < 0.01, ***$P$ < 0.001; non-significant comparisons are not shown).

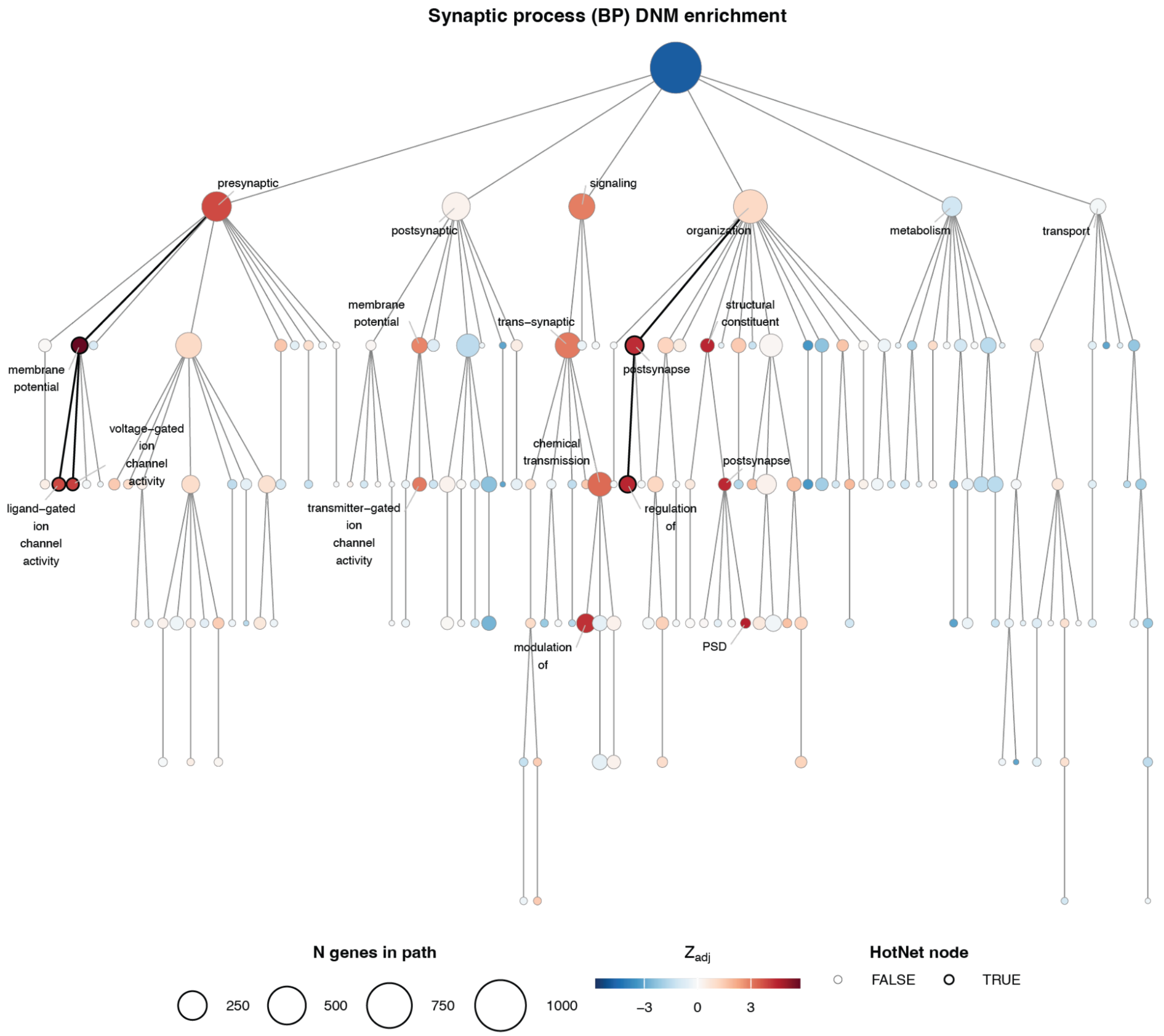

**Fig. S18. Synaptic process HotNet tree for DNM enrichment.**
Each node represents a SynGO term, and node color corresponds to $Z_{adj}$ (enrichment statistic adjusting for gene set size). Nodes outlined in black and connected by dark edges are members of Hierarchical HotNet subnetworks, or groups of compartments with coordinated elevated signal per gene. HotNet nodes and direct children of the root node are labeled, as well as nodes with $Z_{adj} > 2.8$ that are not part of a HotNet. The BP HotNets and constituent GO paths are: Presynaptic ion channels (left; GO:0099505, GO:0099507, GO:0099508) and Postsynaptic organization (GO:0099173, GO:0099175). Abbreviations: PSD = postsynaptic density.

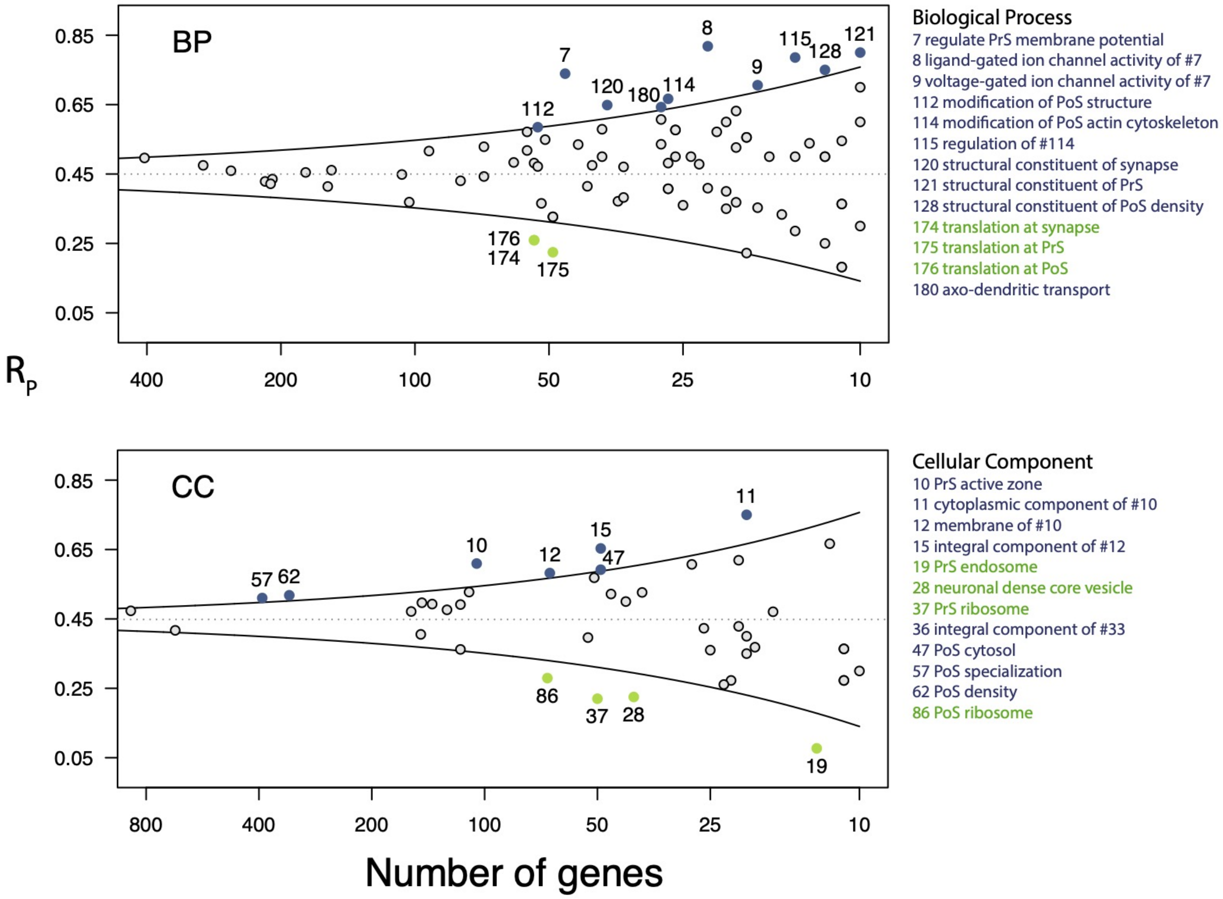

**Fig S19. Average compartment $R_p$ vs number of genes.**

Fraction of genes in compartment with mutation rate greater for cases than for controls ($R_p$) versus number of genes in compartment. BP = biological process of the synapse; CC = cellular component of the synapse.

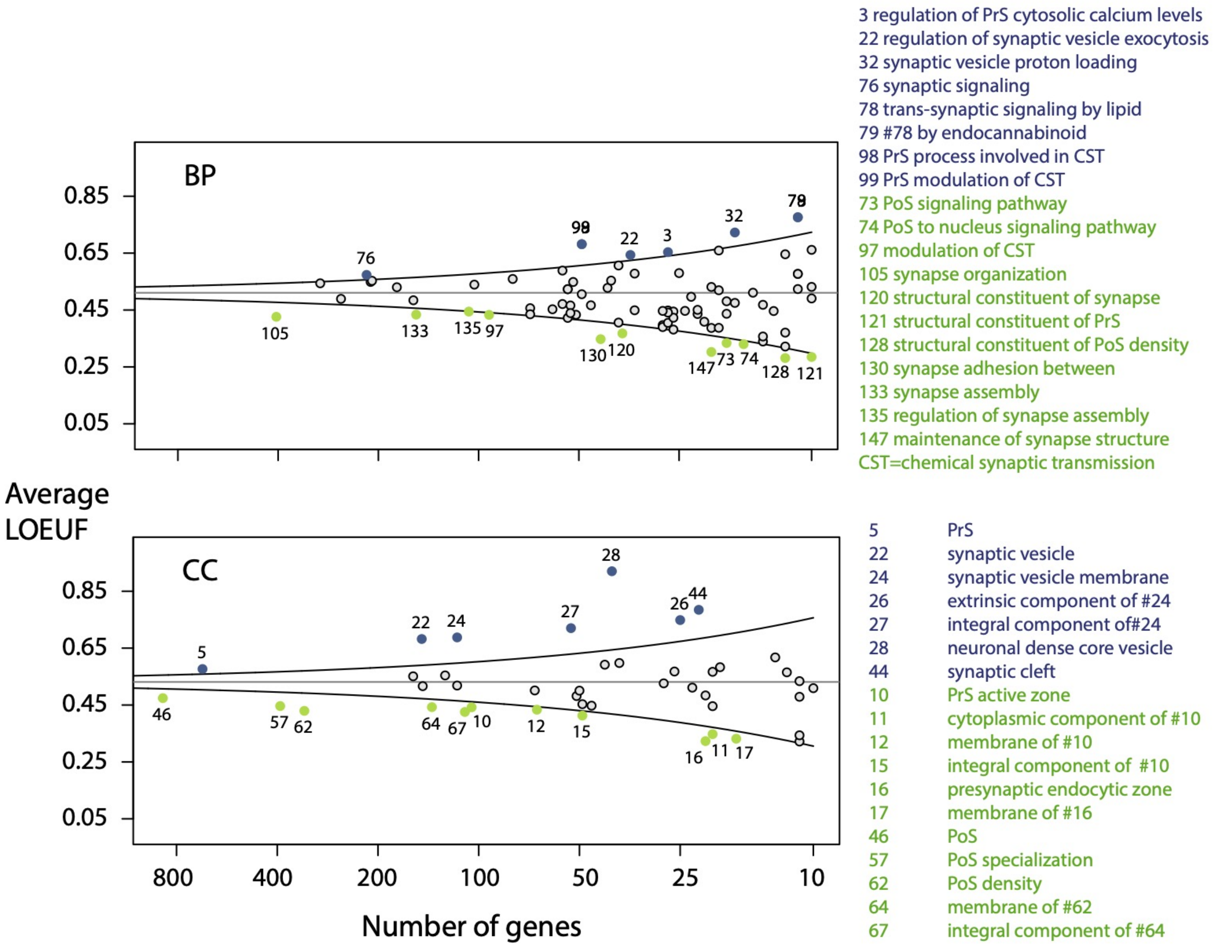

**Fig S20. Average compartment LOEUF vs number of genes.**

Average LOEUF versus number of genes per compartment for. Mean LOEUF over all genes in the SynGO frameworks BP and CC are 0.51 and 0.53, respectively. Using these means and their standard error, we compute the 95% confidence intervals of the mean for each descendant compartment. BP = biological process of the synapse; CC = cellular component of the synapse.

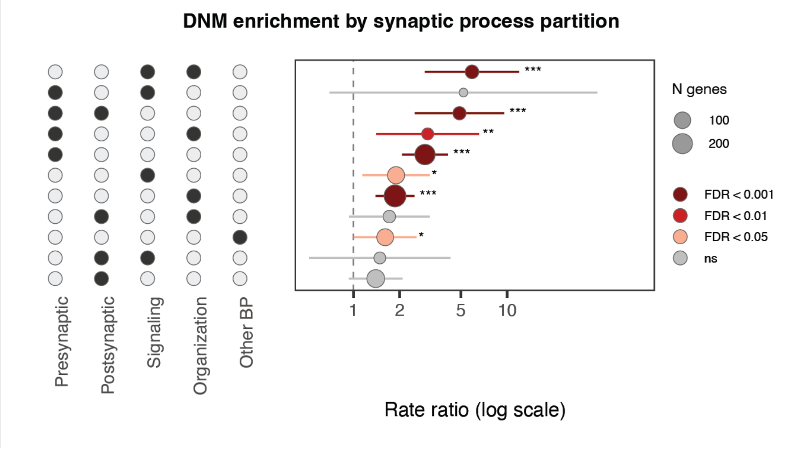

**Fig. S21.** ***De novo*** **mutation enrichment by synaptic process partition.**
Rate ratio of DNMs in ASD probands relative to unaffected siblings for non-overlapping SynGO BP gene partitions defined by the major process categories: presynaptic, postsynaptic, synaptic signaling, synapse organization, and metabolism/transport (Other). Genes annotated to two categories are assigned to the corresponding intersection partition (e.g., Pre+Post; **Methods**). Left: Dot matrix showing the SynGO process category membership of each partition. Right: Rate ratio is shown on a log scale; dot size indicates the number of genes in each partition and error bars show 95% confidence intervals (Poisson approximation). Dot color and asterisks both show the FDR-corrected significance of a one-sided binomial test ($H_0$: equal proband and sibling mutation rates): light red (* FDR < 0.05), medium red (** FDR < 0.01), dark red (*** FDR < 0.001); gray indicates non-significant.

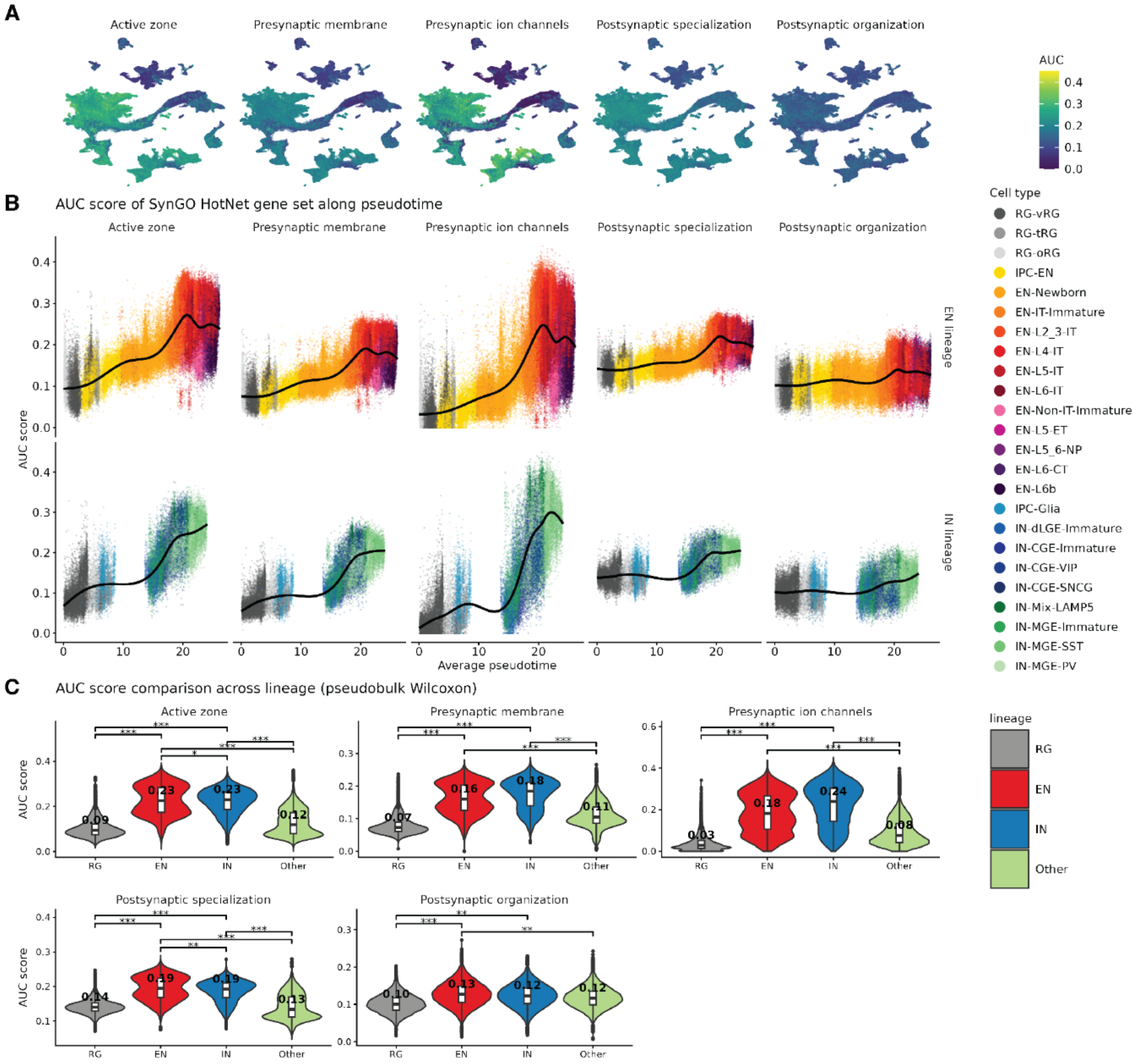

**Fig. S22. AUCell enrichment of SynGO HotNet gene sets across cell types and developmental trajectories.**
**(A)** UMAP visualization of AUC scores for five SynGO HotNet gene sets across all cells. **(B)** AUC scores along pseudotime for EN (top) and IN (bottom) lineages, colored by cell type. Black curves represent GAM-smoothed trends. X-axis indicates average pseudotime rank. **(C)** Pseudobulk Wilcoxon comparison of AUC scores across major lineages (RG, EN, IN, Other) for each HotNet gene set. Median AUC values are shown within each violin. Significance brackets indicate pairwise comparisons (* FDR < 0.05; ** FDR < 0.01; *** FDR < 0.001).

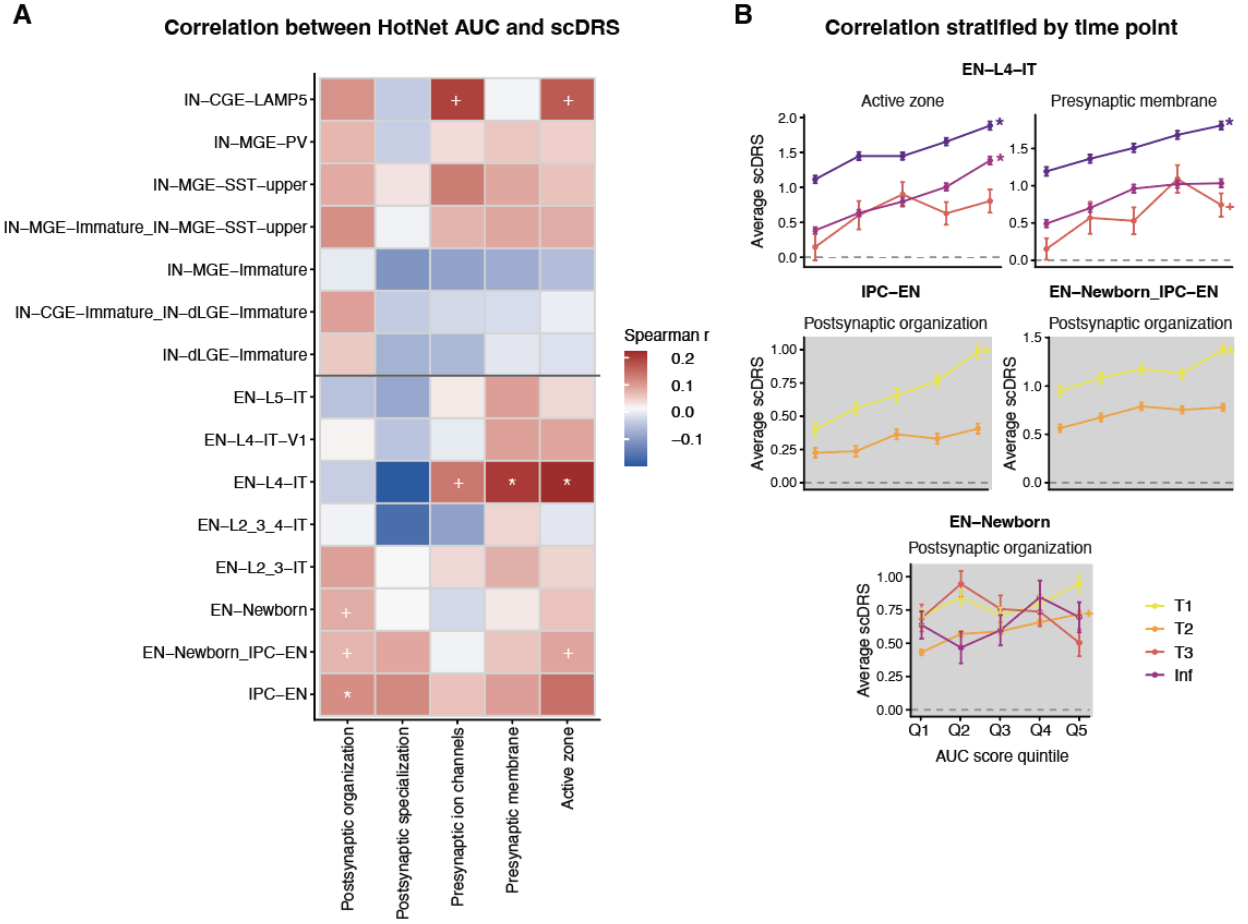

**Fig. S23. Correlation of HotNet AUC scores and scDRS in ASD-relevant cell types**

**(A)** Heatmap showing Spearman correlations between AUC scores for SynGO HotNet subnetworks and scDRS across scDRS-significant cell type clusters (as found in Fig. 1), pooling cells across all developmental stages. **(B)** AUC score quintile plots showing mean scDRS (± SE) for cells binned by AUC score quintile, assessed separately within each developmental stage. Results are stratified by cell type and HotNet subnetworks. T1 = First trimester; T2 = Second trimester; T3 = Third trimester; Inf = Infancy; Adol = Adolescence. In (A), correlations were computed for each cell type by pooling cells across all developmental stages; in (B), correlations for HotNets with significance in (A) were computed separately for each cell type within each developmental stage. Significance was assessed by a two-sided Monte Carlo test comparing the observed correlation against a null distribution derived from 1,000 matched control gene sets. Only cell groups with at least 150 cells were tested, and *P* values were Benjamini-Hochberg corrected. * FDR < 0.05, $^{+}$ P < 0.05.

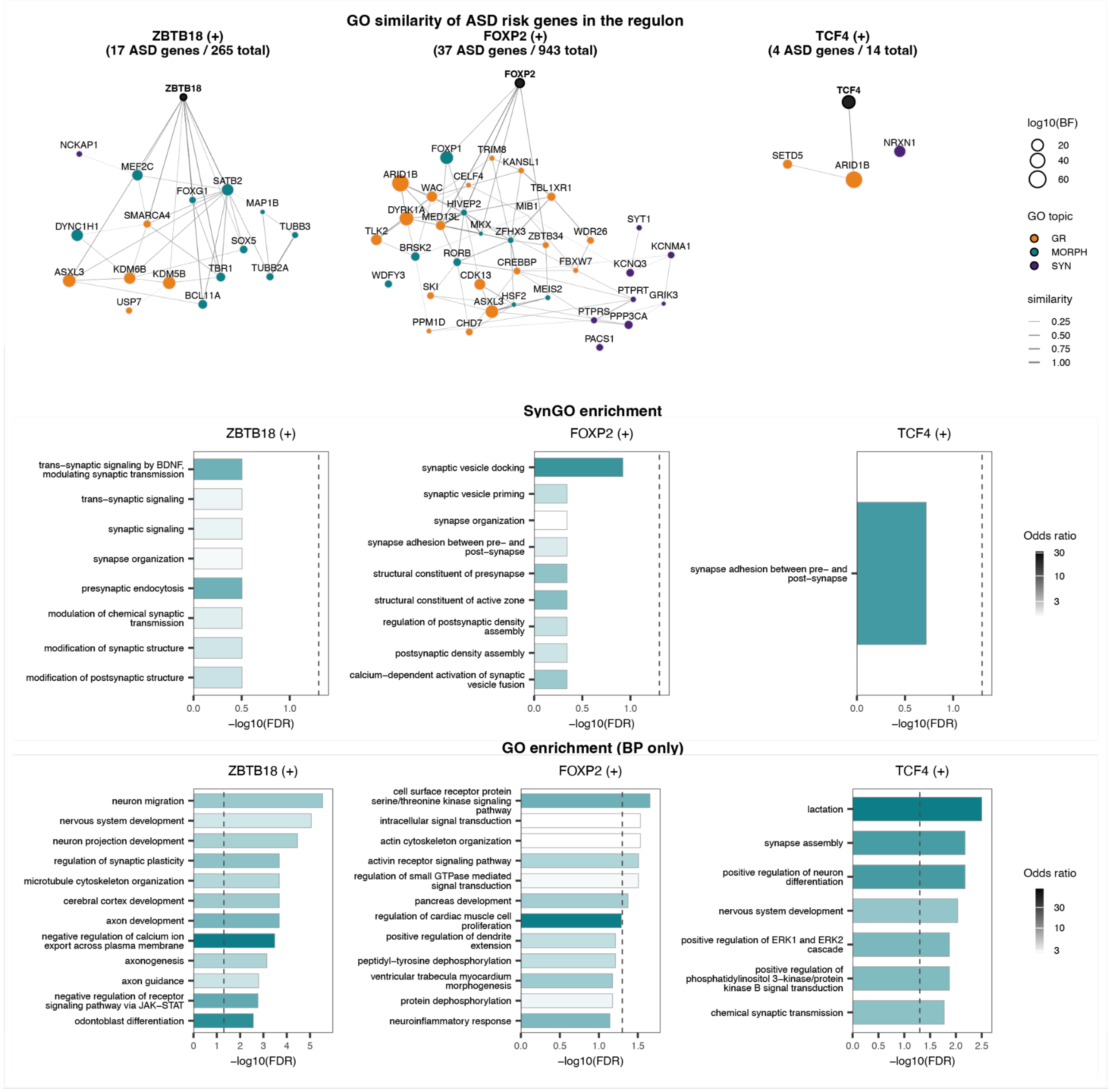

**Fig. S24. Regulons driven by ASD gene transcription factors.**
Top: GO similarity graphs of each regulon. Legend as in Fig. 2D, with nodes colored by biological program assignment. Middle: Nominally significant SynGO enrichments (*P* < 0.05) for each regulon. Vertical dashed line indicates FDR = 0.05; fill represents Fisher's test odds ratio. Bottom: Same as middle but for GO BP enrichments.

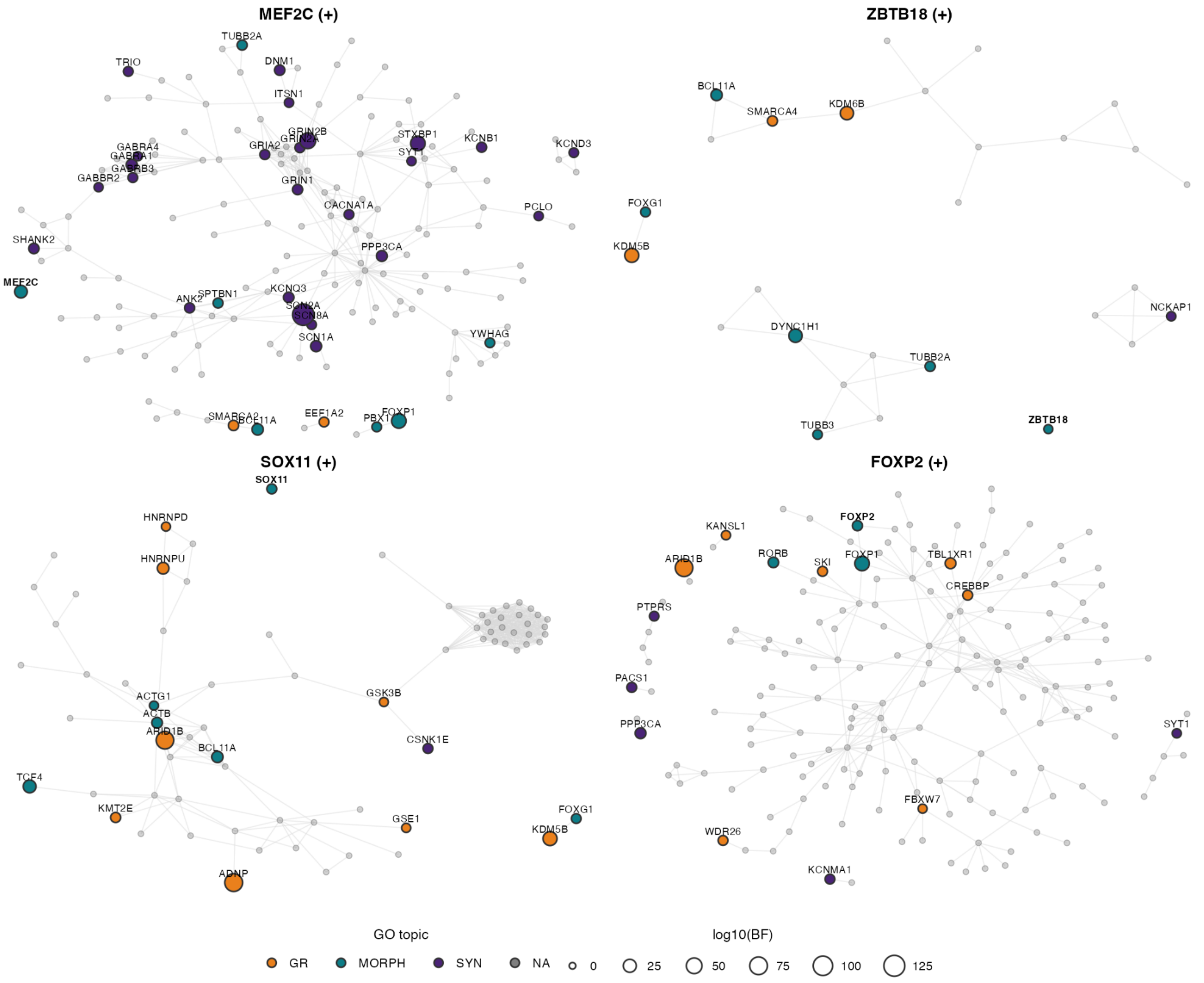

**Fig. S25. High confidence protein-protein interactions within regulons with ASD gene TFs.** STRING (v12) [21] physical interactions among protein products of $MEF2C^{(+)}$, $ZBTB18^{(+)}$, $SOX11^{(+)}$, and $FOXP2^{(+)}$ genes, showing only high-confidence (STRING combined score > 0.7) and only connected components that contain at least one ASD gene (TADA FDR < 0.001). Node fill indicates each gene's biological program assignment; gray nodes are regulon genes that are not ASD genes. Node size corresponds to ASD association strength ($\log_{10}$(BF)).

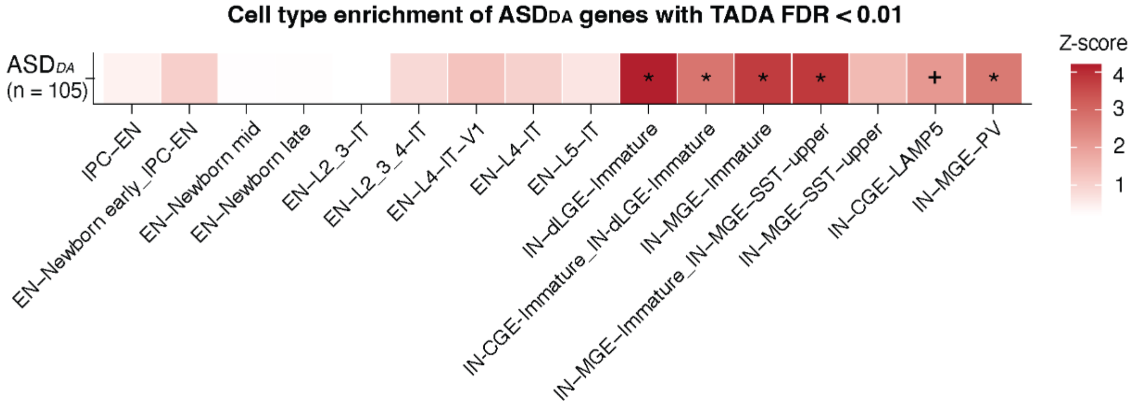

**Fig. S26. Sensitivity analysis of cell type enrichment for $ASD_{DA}$ genes.**
Heatmap shows cluster level enrichment when the TADA FDR threshold for defining $ASD_{DA}$ genes was relaxed to FDR < 0.01 (*n* = 105). Significance: * FDR < 0.05; $^{+}$ *P* < 0.05.

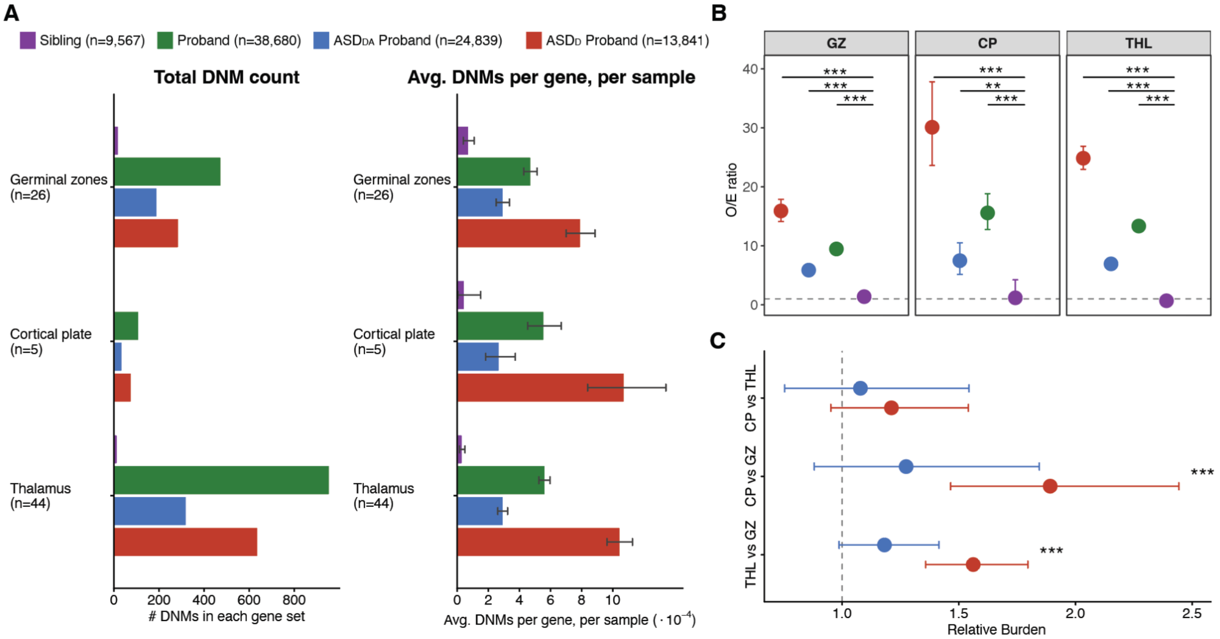

**Fig. S27. Spatial gene set enrichment of ASD genes stratified by DD/ID comorbidity.**
**(A)** Total count of DNMs observed in each regional gene set across four cohorts (Left). Average number of DNMs per gene per sample ($\times10^{-4}$) across regional gene sets, stratified by cohort (Right). Error bars represent 95% confidence intervals. **(B)** O/E ratio of DNMs in ASD genes in cortical plate (CP, $n$ = 5), germinal zone (GZ, $n$ = 26), and thalamus (THL, $n$ = 44). Significance indicates FDR-corrected comparison against unaffected siblings. The dashed line indicates O/E = 1. **(C)** Pairwise comparisons of O/E ratio across spatial gene sets in $ASD_D$ and $ASD_{DA}$ probands, expressed as relative burden with 95% confidence intervals.Significance: ** FDR < 0.01; ***FDR < 0.001.

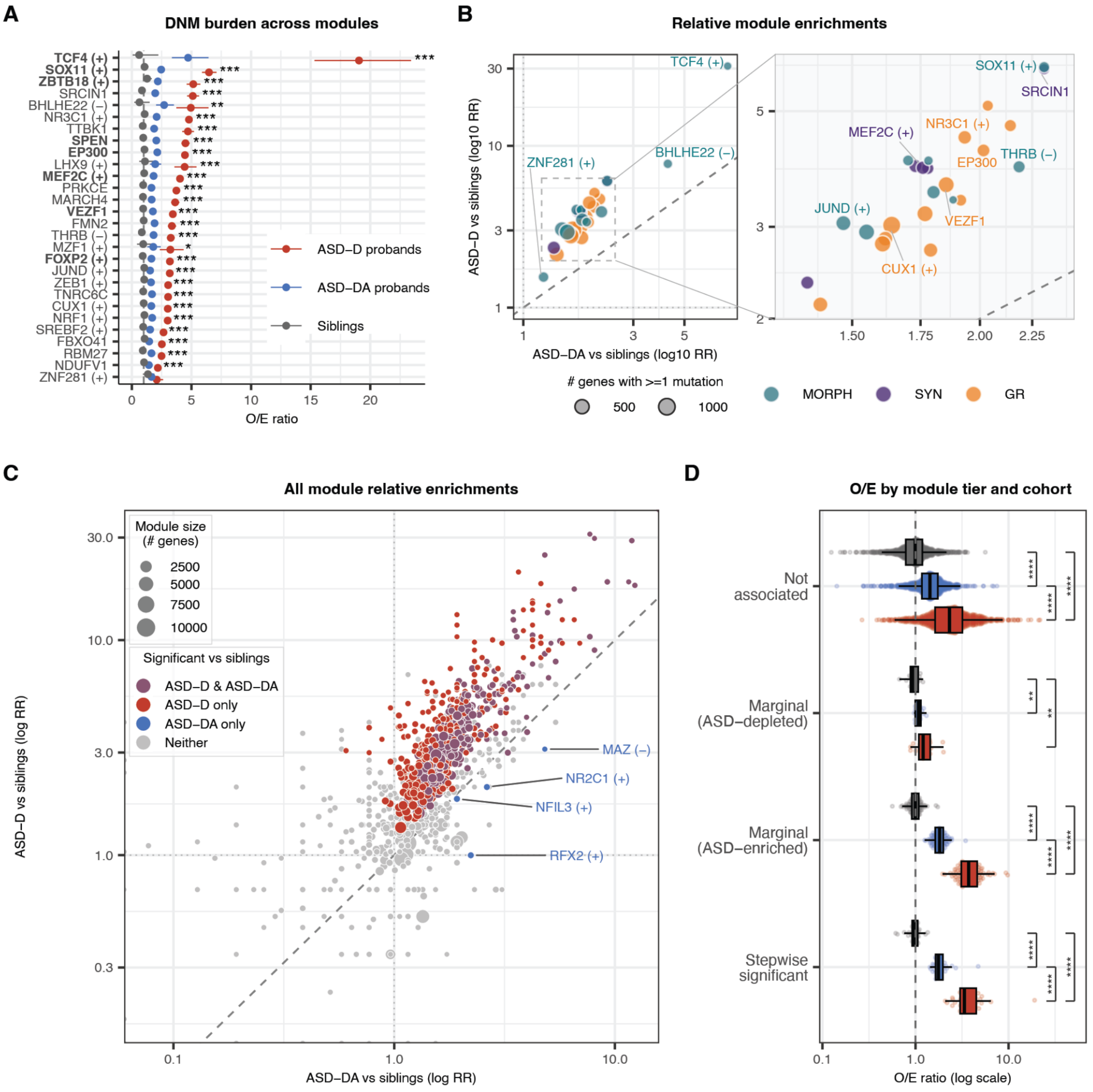

**Fig. S28. Module DNM enrichment stratified by DD/ID comorbidity.**
**(A)** O/E ratio of damaging DNMs for each of the 28 independently ASD-associated modules across three cohorts: ASD$_D$ probands (red; *n* = 13,841, including 1,194 from the Kaplanis et al. DD/ID cohort), ASD$_{DA}$ probands (blue; *n* = 24,839), and unaffected siblings (gray; *n* = 9,567). Error bars show 95% Garwood exact Poisson confidence intervals, the dashed line marks O/E = 1. Asterisks indicate significance of the ASD$_D$ vs ASD$_D$ pairwise comparison (BH-corrected; *FDR < 0.05, **FDR < 0.01, ***FDR < 0.001). As with **Fig. 2**, module labels show the hub gene (co-expression modules) or regulatory factor and direction of effect (regulons; + activating, − repressive). **(B)** Rate ratio (RR) of ASD$_D$ vs siblings (y-axis) and ASD$_{DA}$ vs siblings (x-axis) for each of the 28 ASD-associated modules, on a log scale. The dashed diagonal indicates equal enrichment in both cohorts. Points are colored by biological program and sized by the number of module genes carrying >=1 DNM. **(C)** Scatter plot of RR of damaging DNMs (PTVs, Mis2, Mis1) in ASD$_D$ probands

versus siblings (y-axis) and $ASD_{DA}$ probands versus siblings (x-axis), shown on a log scale, for all 1,151 gene co-expression modules and transcriptional regulons. Each point represents one module, size is proportional to the number of genes. Color indicates significance versus siblings. The dashed diagonal line indicates equal enrichment in $ASD_D$ and $ASD_{DA}$ probands. Labeled modules (blue) are those significant in $ASD_{DA}$ but not $ASD_D$ probands relative to siblings. **(D)** Distribution of O/E ratios across cohorts for modules grouped by their association tier: stepwise-significant modules (independently associated with ASD after conditional forward selection; $n$ = 28), marginally ASD-enriched modules (individually significant at FDR < 0.001 but pruned in stepwise selection, with positive beta coefficient; $n$ = 96), marginally ASD-depleted modules (individually significant with negative beta coefficient; $n$ = 16), and modules not individually associated with ASD ($n$ = 1009). O/E ratios (log-scale) are shown on the x-axis; the dashed vertical line indicates O/E = 1 (no enrichment). Significance brackets indicate pairwise differences between cohorts assessed by conditional binomial test (** FDR < 0.01; **** FDR < 0.0001).

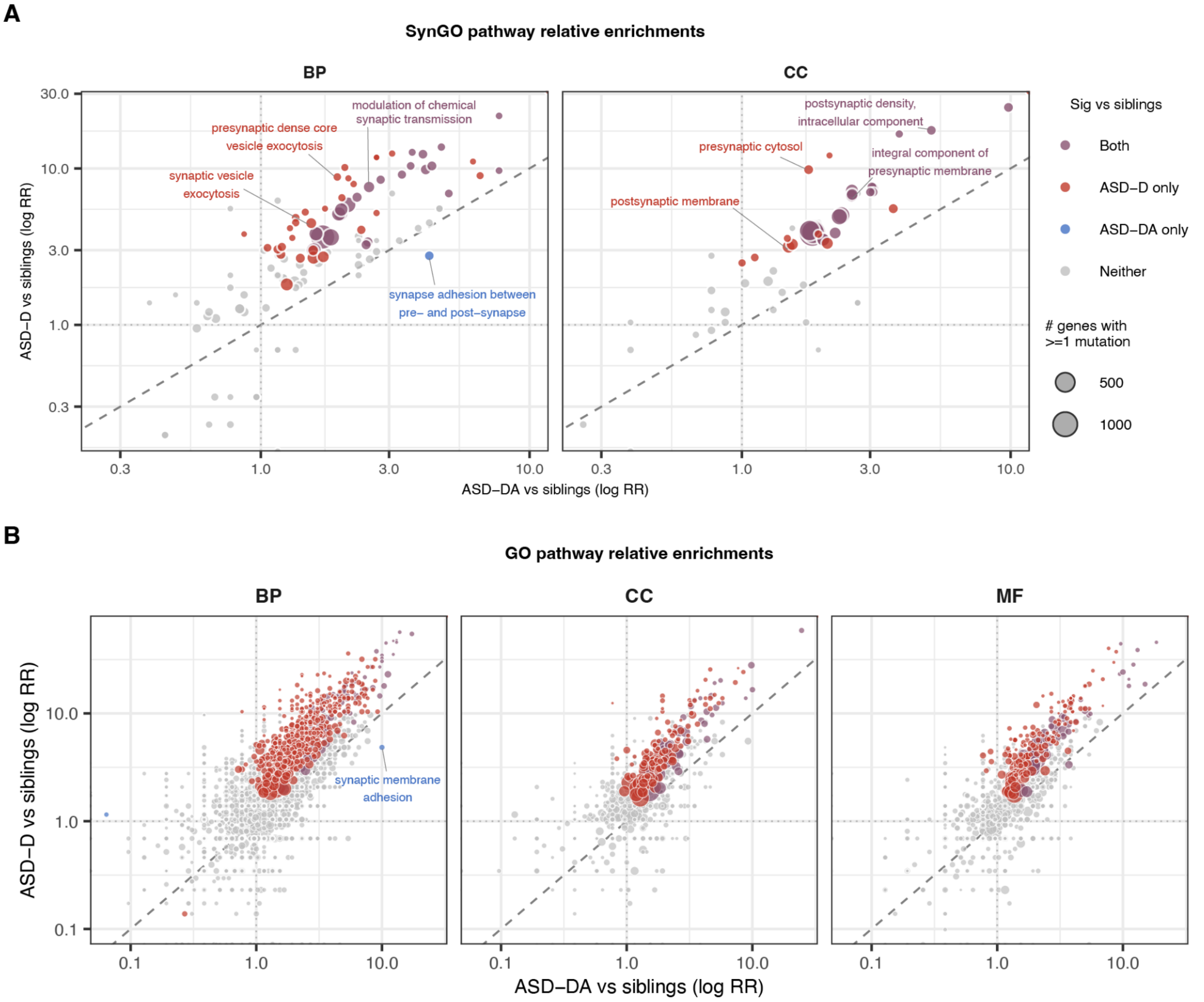

**Fig. S29. Gene ontology pathway DNM enrichment stratified by DD/ID comorbidity.**
**(A)** Rate ratio (RR) of ASD$_{D}$ vs siblings (y-axis) and ASD$_{DA}$ vs siblings (x-axis) for all SynGO pathways, separated by ontology. Points are colored by significance versus siblings: both cohorts (purple), ASD$_{D}$ only (red), ASD$_{DA}$ only (blue), or neither (gray; BH FDR < 0.05). Point size reflects pathway gene count. The dashed diagonal indicates equal enrichment in both cohorts; selected pathways are labeled. **(B)** DNM burden across all GO pathways. Legend as in panel A**.** BP = biological process; CC = cellular component; MF = molecular function.

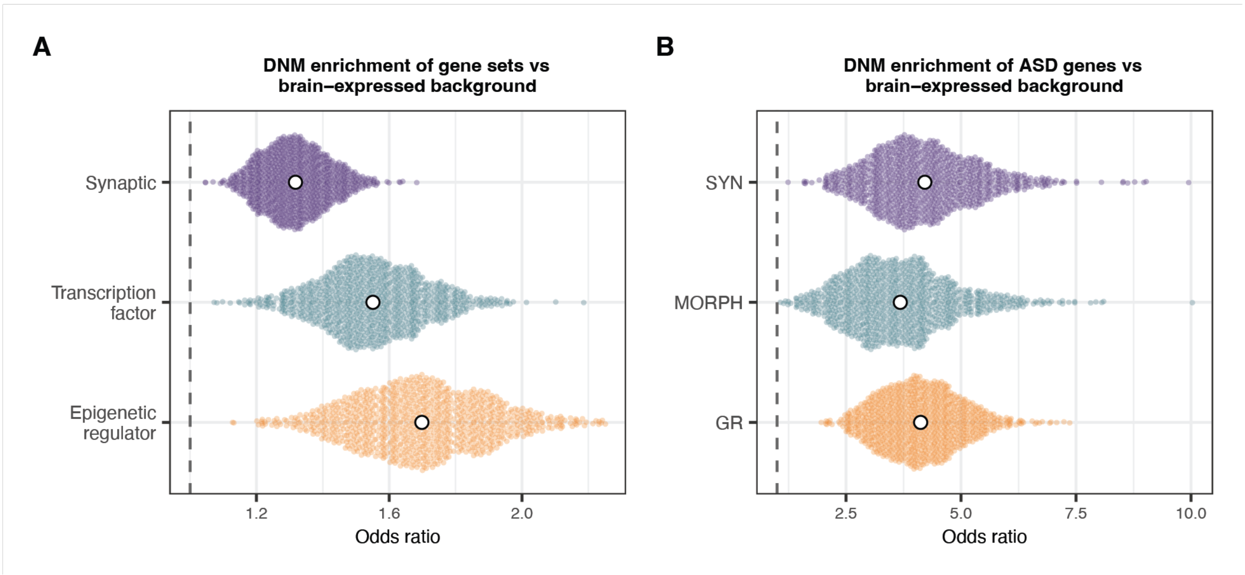

**Fig. S30. Gene sets DNM enrichment relative to brain-expressed background.**
**(A)** Overall enrichment of synaptic, transcription factor, and epigenetic regulator gene sets for ASD *de novo* mutations (DNMs), each compared against a mutation-rate-matched resample of brain-expressed genes (1,000 iterations). Colored dots show the odds ratio (OR) for each resample; white points show the mean OR and 95% CI of the mean. Synaptic OR [95% CI]: 1.32 [1.31, 1.32]; Transcription factors: 1.55 [1.54, 1.56]; Epigenetic regulators: 1.70 [1.69, 1.71]. **(B)** Overall enrichment of ASD genes (TADA FDR < 0.001) assigned to each biological program for ASD *de novo* mutations (DNMs). Panel legend is the same as panel A. SYN OR [95% CI]: 4.12 [4.07, 4.18]; MORPH: 3.68 [3.60, 3.75]; GR: 4.21 [4.14, 4.28].

### Supplementary Table Legends

**Table S1. scDRS group-level enrichment analyses (rare-variant/RVAS gene set). (A)** Group-level scDRS enrichment statistics for cell subclasses. **(B)** Group-level scDRS enrichment statistics for excitatory (EN) and inhibitory (IN) lineage clusters. **(C)** Group-level scDRS enrichment statistics across developmental stages for EN and IN lineages. **(D)** Group-level scDRS enrichment statistics across pseudotime bins for the EN lineage. **(E)** Group-level scDRS enrichment statistics across pseudotime bins for the IN lineage. **(F)** Group-level scDRS enrichment statistics for PFC and V1, pooled across developmental stages in 11 paired donors.

**Table S2. MapMyCell mapping of MGE-SST subtypes to the 10x Whole Human Brain Atlas.** Subtype-specific molecular signatures and mapped reference cell types for the upper-layer and deep-layer MGE-SST subtypes.

**Table S3. Differential gene expression between MGE-SST subtypes.** Genes differentially expressed between the upper-layer (ASD-enriched) and deep-layer (ASD-unenriched) MGE-SST subtypes.

**Table S4. scDRS group-level enrichment analyses (common-variant/GWAS gene set). (A)** Group-level scDRS enrichment statistics across developmental stages for EN and IN lineages**. (B)** Group-level scDRS enrichment statistics across pseudotime bins for the EN lineage. **(C)** Group-level scDRS enrichment statistics across pseudotime bins for the IN lineage**.**

**Table S5. Regional enrichments of the ASD gene set across the adult cortex.** Per-region expression enrichment of the 253 high-confidence ASD genes across 34 Desikan-Killiany cortical regions [22], from the Allen Human Brain Atlas microarray dataset [23]. Includes enrichment *Z*-scores (relative to a size-matched random-gene-set null) and spin-permutation *P* values (BH-corrected across regions).

**Table S6. Gene module association with ASD rare variant vulnerability. (A)** Statistics for the 28 modules independently associated with ASD after conditional forward selection. **(B)** Marginal association statistics for all 1,151 modules included in the analysis.

**Table S7. Functional and cell-type enrichments of ASD-associated modules. (A)** Gene Ontology overrepresentation results for each module. **(B)** Cell-type specificity of each module, shown as the proportion of AUCell [24] high-scoring cells per developmental cell type [19].

**Table S8. GO topic model defining three functional programs. (A)** Top GO terms per topic, used to annotate topics according to their biological function. Ten words for each of prob (highest probability; words with the highest overall frequency within the topic), frex (words that are both frequent and exclusive to this topic), lift (words that are more common in this topic than in the rest of the dataset), and score (weighted measure emphasizing words that are frequent and high-contrast against other topics). **(B)** Per-module topic proportions (gamma) and dominant-program assignments (`primary_program`).

**Table S9. Assignment of ASD genes to programs and functional subclusters. (A)** Mutually exclusive program assignment for each of the 253 ASD genes. **(B)** GO-based functional subclusters within the GR program. **(C)** GO-based functional subclusters within the MORPH program.

**Table S10. Program-level regional enrichment across the adult cortex.** Per-region expression enrichment for each functional program across 34 Desikan-Killiany cortical regions (adult Allen Human Brain Atlas), as in **Table S5**. **(A)** GR. **(B)** SYN. **(C)** MORPH.

**Table S11. Program-level spatial enrichment in the developing cortex.** Expression enrichment of each functional program (GR, MORPH, SYN) across compartments and areas of the developing human cortex (μbrain atlas) [12], tested against two null backgrounds. **(A)** transcriptome-wide background. **(B)** ASD gene set background.

**Table S12. ASD genes overlapping sensorimotor-association patterning transcription factors**. MORPH-assigned ASD genes and transcription factors that overlap the areal-patterning factors used by Tsyporin et al. [20] to define the sensorimotor–association axis, with each gene's description and assigned pole.

**Table S13. SynGO compartment-level de novo mutation enrichment and module-synapse enrichments.** Damaging DNM burden of ASD probands versus unaffected siblings across the SynGO synaptic gene ontology. (**ReadMe**) column definitions. **(A)** Cellular (synaptic) Component compartments. **(B)** Biological (synaptic) Process compartments. **(C)** Module-synapse (SynGO CC root term) enrichment results using overrepresentation analysis (ORA). **(D)** Module-HotNet enrichment results using ORA.

**Table S14. Stratification of high-confidence ASD genes by DD/ID-associated mutation burden and associated scDRS enrichment analyses. (A)** Classification of high-confidence ASD genes into $ASD_{DA}$ and $ASD_{DH}$/$ASD_{DL}$ groups, based on per-proband mutation rate and mutation fraction relative to the background expectation $p_0$. **(B)** Group-level scDRS enrichment statistics for EN and IN lineage clusters, for the $ASD_{DH}$, combined $ASD_{DL}$+$ASD_{DA}$, and $ASD_{DA}$ gene sets. **(C)** Group-level scDRS enrichment statistics across developmental stages for the $ASD_{DH}$, combined $ASD_{DL}$+$ASD_{DA}$, and $ASD_{DA}$ gene sets.

**Table S15. DNM burden of spatially enriched gene sets in the developing thalamus and cortex. (A)** Germinal zone (GZ), cortical plate (CP), and thalamus (THL) gene sets from a prenatal spatial transcriptomic atlas [14], overlapping the 253 high-confidence ASD genes. **(B)** O/E ratios and DNM burden for GZ-, CP-, and THL-enriched gene sets in ASD probands versus unaffected siblings. **(C)** Pairwise comparisons of O/E ratios across GZ-, CP-, and THL-enriched gene sets. **(D)** Pairwise comparisons of O/E ratios across GZ-, CP-, and THL-enriched gene sets within $ASD_D$ and $ASD_{DA}$ probands.

**Table S16. Module-level burden stratified by DD/ID comorbidity. (A)** Observed vs. expected damaging DNM burden within each module for $ASD_D$, $ASD_{DA}$, all probands, and unaffected siblings. **(B)** Pairwise between-cohort rate ratios.

**Table S17. SynGO compartment burden stratified by DD/ID comorbidity. (A)** O/E ratios of damaging DNM burden within each SynGO compartment for the same four cohorts. **(B)** Pairwise between-cohort rate ratios.

**Table S18. SynGO BP partitions** Mutation rates and contrasts of these rates for independent subsets of genes formed from major children of process of the synapse. In the upper triangle, the ratio of de novo rates in ASD cases (column gene set divided row gene set). In the lower triangle, *P* values assume a binomial distribution for mutations. Rightmost table, number of genes per set; number of de novo mutations identified in ASD cases; and the ratio of these to form a rate. Pre =process in the presynapse; Post = process in the postsynapse; Sig = synaptic signaling; Org = synapse organization; Oth = Metabolism & Trans*p*ort.

### References

1. Allen Institute for Brain Science. MapMyCells [Software]. RRID: SCR_024672. https://knowledge.brain-map.org/mapmycells/process?refTaxonomyId=10xGene (2025).
2. Arora, S. *et al.* A Practical Algorithm for Topic Modeling with Provable Guarantees. in *International Conference on Machine Learning* 280–288 (PMLR, 2013).
3. Roberts, M. E., Stewart, B. M. & Tingley, D. Stm: An R package for structural topic models. *J. Stat. Softw.* **91**, 1–40 (2019).
4. Samocha, K. E. *et al.* A framework for the interpretation of de novo mutation in human disease. *Nat. Genet.* **46**, 944–950 (2014).
5. Fu, J. M. *et al.* Rare coding variation provides insight into the genetic architecture and phenotypic context of autism. *Nat. Genet.* **54**, 1320–1331 (2022).
6. Zylka, M. J., Simon, J. M. & Philpot, B. D. Gene length matters in neurons. *Neuron* **86**, 353–355 (2015).
7. Sugino, K. *et al.* Mapping the transcriptional diversity of genetically and anatomically defined cell populations in the mouse brain. *Elife* **8**, e38619 (2019).
8. Fulcher, B. D., Arnatkeviciute, A. & Fornito, A. Overcoming false-positive gene-category enrichment in the analysis of spatially resolved transcriptomic brain atlas data. *Nat. Commun.* **12**, 2669 (2021).
9. Karczewski, K. J. *et al.* The mutational constraint spectrum quantified from variation in 141,456 humans. *Nature* **581**, 434–443 (2020).
10. Dear, R. *et al.* Cortical gene expression architecture links healthy neurodevelopment to the imaging, transcriptomics and genetics of autism and schizophrenia. *Nat. Neurosci.* **27**, 1075–1086 (2024).
11. Gandal, M. J. *et al.* Broad transcriptomic dysregulation occurs across the cerebral cortex in ASD. *Nature* **611**, 532–539 (2022).
12. Ball, G. *et al.* Molecular signatures of cortical expansion in the human foetal brain. *Nat.*

*Commun.* **15**, 9685 (2024).

13. Lenth, R. V. & Piaskowski, J. emmeans: Estimated Marginal Means, aka Least-Squares Means. https://CRAN.R-project.org/package=emmeans (2026).
14. Aivazidis, A. *et al.* A spatial transcriptomic atlas of autism-associated genes identifies convergence in the developing human thalamus. *bioRxivorg* 2025.11.05.685843 (2025) doi:10.1101/2025.11.05.685843.
15. Uhlén, M. *et al.* Proteomics. Tissue-based map of the human proteome. *Science* **347**, 1260419 (2015).
16. Lambert, S. A. *et al.* The human transcription factors. *Cell* **172**, 650–665 (2018).
17. Wang, J., Shi, A. & Lyu, J. A comprehensive atlas of epigenetic regulators reveals tissue-specific epigenetic regulation patterns. *Epigenetics* **18**, 2139067 (2023).
18. Harris, K. D. & Shepherd, G. M. G. The neocortical circuit: themes and variations. *Nat. Neurosci.* **18**, 170–181 (2015).
19. Wang, L. *et al.* Molecular and cellular dynamics of the developing human neocortex. *Nature* 1–10 (2025).
20. Tsyporin, J. *et al.* Competing programs shape cortical sensorimotor-association axis development. *Nature* 1–12 (2026).
21. Szklarczyk, D. *et al.* The STRING database in 2023: protein-protein association networks and functional enrichment analyses for any sequenced genome of interest. *Nucleic Acids Res.* **51**, D638–D646 (2023).
22. Desikan, R. S. *et al.* An automated labeling system for subdividing the human cerebral cortex on MRI scans into gyral based regions of interest. *Neuroimage* **31**, 968–980 (2006).
23. Hawrylycz, M. J. *et al.* An anatomically comprehensive atlas of the adult human brain transcriptome. *Nature* **489**, 391–399 (2012).
24. Aibar, S. *et al.* SCENIC: single-cell regulatory network inference and clustering. *Nat. Methods* **14**, 1083–1086 (2017).
